# Graph-based Nanopore Adaptive Sampling with GNASTy enables sensitive pneumococcal serotyping in complex samples

**DOI:** 10.1101/2024.02.11.579857

**Authors:** Samuel T. Horsfield, Basil Fok, Yuhan Fu, Paul Turner, John A. Lees, Nicholas J. Croucher

**Author notes:** These authors contributed equally to this work (John A. Lees, Nicholas J. Croucher).

## Abstract

Serotype surveillance of *Streptococcus pneumoniae* (the pneumococcus) is critical for understanding the effectiveness of current vaccination strategies. However, existing methods for serotyping are limited in their ability to identify co-carriage of multiple pneumococci and detect novel serotypes. To develop a scalable and portable serotyping method that overcomes these challenges, we employed Nanopore Adaptive Sampling (NAS), an on-sequencer enrichment method which selects for target DNA in real-time, for direct detection of *S. pneumoniae* in complex samples. Whereas NAS targeting the whole *S. pneumoniae* genome was ineffective in the presence of non-pathogenic streptococci, the method was both specific and sensitive when targeting the capsular biosynthetic locus (CBL), the operon that determines *S. pneumoniae* serotype. NAS significantly improved coverage and yield of the CBL relative to sequencing without NAS, and accurately quantified the relative prevalence of serotypes in samples representing co-carriage. To maximise the sensitivity of NAS to detect novel serotypes, we developed and benchmarked a new pangenome-graph algorithm, named GNASTy. We show that GNASTy outperforms the current NAS implementation, which is based on linear genome alignment, when a sample contains a serotype absent from the database of targeted sequences. The methods developed in this work provide an improved approach for novel serotype discovery and routine *S. pneumoniae* surveillance that is fast, accurate and feasible in low resource settings. GNASTy therefore has the potential to increase the density and coverage of global pneumococcal surveillance.

**One sentence summary:** Pangenome graph-based Nanopore Adaptive Sampling, presented in our tool GNASTy, is a sensitive, portable and cost-effective method for *Streptococcus pneumoniae* surveillance.

## 1 Introduction

*Streptococcus pneumoniae* (the ‘pneumococcus’) is a human nasopharyngeal commensal which can cause severe diseases, such as pneumonia, bacteremia and meningitis, disproportionately affecting young children and the elderly (*1*). *S. pneumoniae* infections cause a significant global health burden, being associated with more than 800,000 deaths annually (*2*), and are the leading cause of death in children under 5 years of age (*3*, *4*). The species can be divided into >100 serotypes (*5*), each of which expresses an immunologically-distinct polysaccharide capsule that enables the bacterium to evade the host’s immune response (*6*).

Polysaccharide conjugate vaccines (PCVs) target a subset of *S. pneumoniae* serotypes that cause a substantial proportion of invasive pneumococcal disease (IPD) (*7*), driving a reduction in the global IPD burden (*3*). This is achieved through a significant perturbation of the pneumococcal population carried in the nasopharynx. Consequently, vaccine-targeted serotypes have been replaced through the expansion of already common serotypes not included in current formulations, and the emergence of previously rare or unknown serotypes, changing the frequency of antimicrobial resistance (AMR) and incidence of disease in *S. pneumoniae* (*8–10*). Ongoing serotype surveillance is critical to identify significant increases in non-vaccine serotype prevalence, particularly if such a serotype is associated with AMR or high invasiveness (*11*). Such dynamics can be monitored through analysis of nasopharyngeal samples, although the frequent carriage of multiple serotypes within a single individual, known as ‘co-carriage’ or ‘co-colonisation’, makes identification of all circulating serotypes challenging (*12*). This problem is exacerbated by the recent discovery that minority serotypes are often present at low frequency (*<* 25% of pneumococcal cells within an individual), but are still responsible for a notable proportion of transmission events (*13*). Therefore, scalable high sensitivity serotype assays that can deconvolute mixed samples, and identify novel serotypes, are necessary to update vaccine formulations and public health strategies in response to pneumococcal epidemiological dynamics (*14*).

The original methods for serotyping pneumococci assay the ability of an unknown isolate to agglutinate in the presence of different antisera that recognise known serotypes (*15*, *16*). Agglutination assays have high specificity when applied by experts, but have extensive training requirements, as precise typing requires a succession of tests with different antisera (*17*). When applied to individual colonies, such methods have a low sensitivity for detecting co-carriage, although this can be improved using latex agglutination of plate sweeps (*15*). Nevertheless, these methods cannot identify novel serotypes; these may be discovered through whole genome sequencing (WGS) approaches, which detect specific sequence variants of the Capsular Biosynthetic Locus (CBL) (*18–20*), the operon which defines pneumococcal serotype (*21*). Yet WGS of individual colonies is difficult to deploy at scale in resource-limited settings as it is expensive and time-consuming, requiring specific expertise and access to specialist laboratory equipment (*22*). The limited number of colonies from a single patient that can be feasibly sequenced limits the ability of WGS to detect co-carriage, unless a sample is subjected to deep sequencing (*13*). However, this reduces the number of samples that can be analysed, and therefore lowers overall throughput. Additionally, both agglutination assays and WGS rely on prior selective culture of *S. pneumoniae* as means of enrichment to improve sensitivity. Selective culture adds additional time, resource and expertise requirements to already complex laboratory workflows, limiting throughput, and potentially resulting in false negatives if cells fail to grow (*23*). Purely genotypic approaches, such as PCR and DNA microarrays, target CBL DNA sequences directly present in the sample and therefore do not require selective culture. These methods can identify co-carriage, and are less laborious and expensive than agglutination assays or WGS, and can therefore be used in high-throughput settings (*22*). However, these methods require CBL sequences to be specified *a priori*, and so cannot detect novel serotypes. Overall, no current serotyping method can scalably and sensitively identify both known and novel serotypes, as well as co-carriage.

Novel nucleotide sequencing approaches have the potential to allow accurate, simple and relatively inexpensive culture-free *S. pneumoniae* surveillance. Nanopore sequencing, developed by Oxford Nanopore Technologies (ONT), is a portable long-read nucleotide sequencing technology in which DNA or RNA molecules are sequenced as they move across an impermeable membrane through protein nanopores (*24*, *25*). Reads are generated in real-time, enabling on-flowcell enrichment of sequences of interest, referred to as ‘target’ DNA, via rejection of all other sequences, referred to as ‘non-target’ DNA. These methods, known collectively as Nanopore Adaptive Sampling or ‘NAS’, align the first segment of DNA fragments as they pass through a nanopore to a reference database, before sending a signal back to the sequencer to either ‘accept’, where the fragment is sequenced to completion, or ‘reject’, where voltage across the nanopore is reversed, ejecting the fragment (*26*). NAS increases target sequence yield by rejecting non-target DNA, increasing the sensitivity for detecting of sequences of interest (*26*, *27*). This makes NAS well-suited for metagenomics, the culture-free DNA sequencing-based analysis of mixed samples (*28*), such as nasopharyngeal communities. NAS has been shown to increase target yield approximately four-fold (*29*, *30*), and by extension enabling multiplexing of samples on ONT devices to increase throughput. Furthermore, increased target yield has been shown to improve accuracy of downstream analyses such as variant calling and assembly when analysing metagenomes (*27*, *31*, *32*).

NAS is available as part of the standard ONT sequencing software platform. However, there has been limited quantification of its accuracy, particularly in metagenomics. It has been previously shown that NAS sensitivity is highest when a target present in a metagenome is closely related to a sequence in the reference database (*31*, *33*). However, high genetic relatedness between non-target and target taxa in the same sample has the potential to negatively impact NAS specificity, as target and non-target reads will be more difficult to distinguish between during the rejection process. The sequence similarity between *S. pneumoniae* and other members of the *Streptococcus* genus (*34*, *35*), which are also present as part of the upper respiratory tract microbiome (*36*), is comparable with the error rate of individual ONT reads (*37*). Hence attempts to enrich for a whole *S. pneumoniae* genome may be limited by the challenge of resolving pneumococcal DNA from that of non-pathogenic streptococci. Alternatively, targeting loci which are specific to *S. pneumoniae* will improve target enrichment, as such sequence are typically absent from benign commensals. Hence CBL sequences are a promising candidate for targeted metagenomics enrichment (*7*, *21*, *38*).

Here, we develop and test methods for metagenome-based NAS with the aim of enhancing the surveillance of *S. pneumoniae*. We show that NAS can be used to implement a scalable method of serotype detection, with high sensitivity and specificity, which can quantify serotype prevalence in co-carriage samples and detect novel serotypes. Furthermore, this method has the potential to be used without the requirement of culturing samples. To improve performance when detecting novel serotypes, we further developed a graph-based bioinformatic method for NAS, named GNASTy (Graph-based Nanopore Adaptive Sampling Typing, pronounced ‘nasty’), and benchmark it against the current NAS implementation, which uses linear alignment. We show that GNASTy outperforms standard linear alignment when enriching for a novel CBL sequence not present in the reference database. This work demonstrates the advantages and caveats of NAS for application in *S. pneumoniae* surveillance, and provides a novel tool for detection and discovery novel serotyping loci in metagenomes.

## 2 Results

### 2.1 NAS performance depends on microbiome composition

We first set out to determine the taxonomic range across which NAS can effectively enrich for a target sequence, whilst still correctly rejecting non-target sequences. We hypothesised that NAS would fail to enrich for target loci when the sequence similarity between target and non-target genomes was comparable to the single-strand error rate of ONT reads (*∼* 6% (*37*)), resulting in incorrect selection of non-target DNA that ultimately reduces target enrichment.

To test this hypothesis, we generated mock communities containing mixtures of genomic DNA from *S. pneumoniae*, the serotype 23F pneumococcal isolate ATCC 700669 (*39*), referred to as ‘Spn23F’, with that of closely and distantly related non-target species. Spn23F DNA was mixed with DNA from species from a different phylum, represented by *Escherichia coli* DH5-*α*; the same genus but different species, represented by *Streptococcus mitis* SK142; and the same species but different strain, represented by *S. pneumoniae* R6 (Figure 1a). To test NAS sensitivity at low target DNA concentrations, Spn23F DNA was titrated from a proportion of 0.5 to 0.001 (50% *−* 0.1%) (Figure 1b) in non-target DNA. These proportions describe the ratio of total DNA bases within a sample which belong to target DNA. The choices of alignment parameters used for NAS performance comparisons are detailed in Supplementary Material (Section A.1). All libraries were size-selected to remove DNA fragments < 10 kb in length, as this was shown to improve enrichment (Supplementary Material, Section A.2). NAS was carried out using Readfish (*26*), targeting either the whole Spn23F genome or 23F CBL (Figure 1c). All samples were multiplexed into a combined sequencing library and run on a single flow cell to control for batch effects. Half of the ‘channels’ (a group of four pores, of which only one is sequencing at one time) sequenced the library using NAS, while the other half sequenced the same library normally without NAS (termed ‘control’). Splitting the flow cell in this way provides an internal control which is used for calculation of enrichment by composition (referred to further as ‘enrichment’, see Section 4.6) (*31*). Using enrichment allows direct comparison of NAS performance across sequencing runs which may otherwise be confounded by between-run variability. Enrichment *>* 1 indicates that a target was successfully enriched, with a greater proportion of target bases generated using NAS relative to the control.

**Figure 1:**
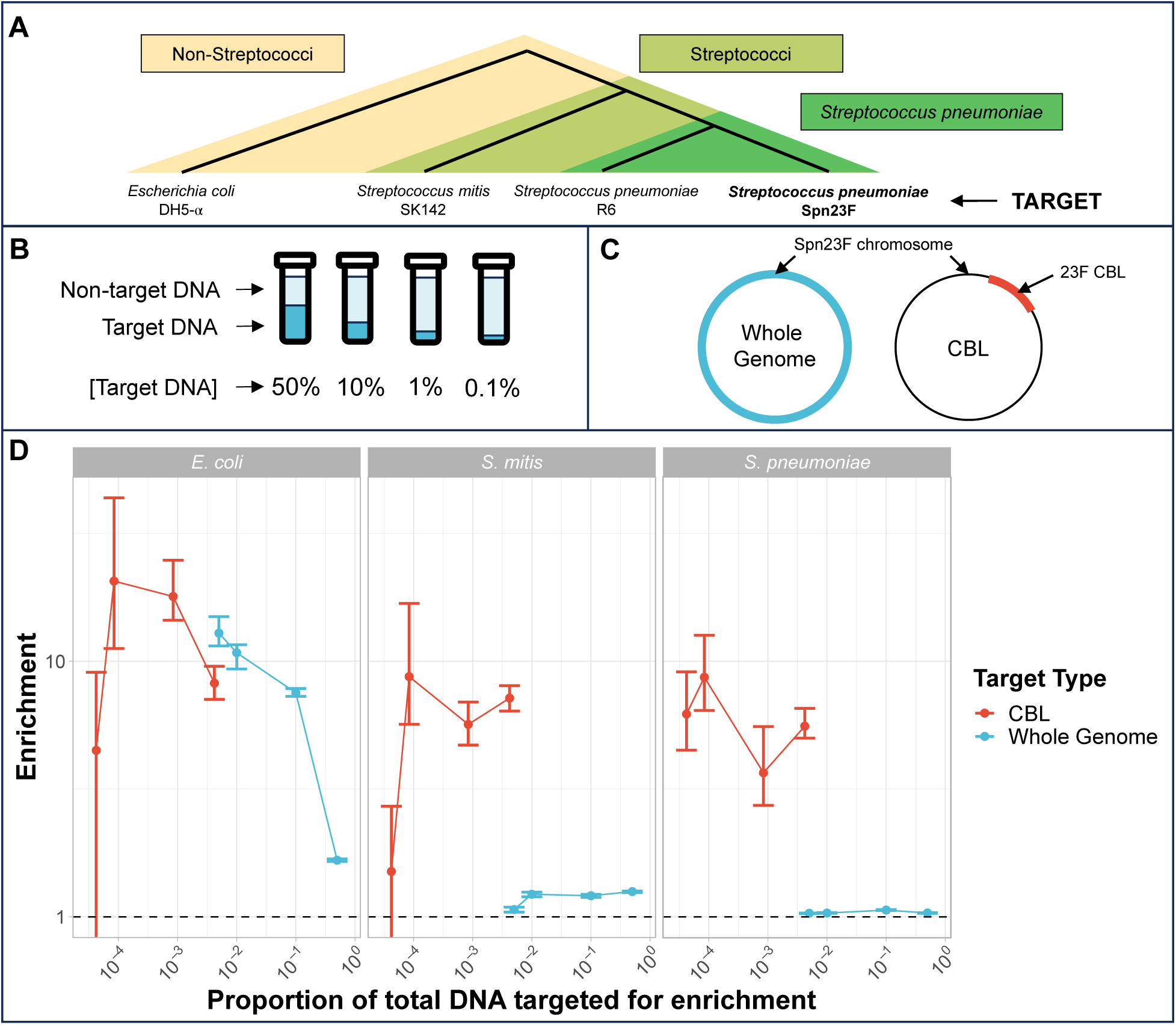
Enrichment of *S. pneumoniae* Spn23F in samples containing closely and distantly related non-target species. (**a**) Representation of evolutionary relatedness of non-target species and the target, *S. pneumoniae* Spn23F. (**b**) Experimental setup of target DNA dilution series with non-target DNA. (**c**) Representation of two enrichment experiments, either targeting the whole Spn23F genome (blue) or the 23F CBL sequence present on the Spn23F chromosome. (**d**) Enrichment results of Spn23F whole genome or 23F CBL at different concentrations of target DNA. Bar ranges are inter-quartile range of enrichment from 100 bootstrap samples of reads. Data points connected by lines are observed enrichment values for each library, with solid lines connecting target DNA diluted at different concentrations with non-target DNA. Columns describe the non-target species within each mixture. To plot on a log scale, all enrichment values had 0.01 added to them. Horizontal dashed line describes enrichment = 1 i.e. no enrichment has occurred.

Comparison of NAS performance based on enrichment is shown in Figure 1d (blue). Spn23F whole genome enrichment was highest in mixtures containing *E. coli* for all target proportions. Conversely, mixture with *S. mitis* and *S. pneumoniae* resulted in notably lower enrichment, although enrichment was slightly higher in mixtures with *S. mitis*. For example, for the 0.1 target dilution, enrichment of the Spn23F genome was 7.51, 1.20 and 1.05 for the *E. coli*, *S. mitis* and *S. pneumoniae* mixtures. Additionally, enrichment increased monotonically as Spn23F DNA proportions decreased in *E. coli* mixtures, as observed previously (*31*), whilst for mixtures with *S. mitis* and *S. pneumoniae* enrichment remained relatively constant between dilutions. These results indicate that the NAS alignment process is not able to effectively reject sequences from non-target species when their divergence is similar to the read error rate. This result has particular significance for use of NAS in *S. pneumoniae* surveillance, as the presence of commonly co-occurring streptococci in the nasopharynx greatly impacts NAS performance.

To determine the effect of non-specific enrichment on downstream analyses, we then assembled reads using MetaFlye (*40*) and analysed assembly quality using Inspector (*41*), overlaying results on the Spn23F reference genome for the 0.1 target DNA dilutions. We compared the relative read coverage and aligned contig coverage of the reference genome, as well as the presence of small (*<* 50 bp) and large (*≥* 50 bp) assembly errors (Figure 2). Greater coverage by aligned contigs indicates that read coverage, and therefore target yield, was sufficiently high to generate a contiguous assembly, whilst presence of small or large errors suggests problems with the assembly process, such as insufficient read coverage or integration of non-target reads into assemblies.

**Figure 2:**
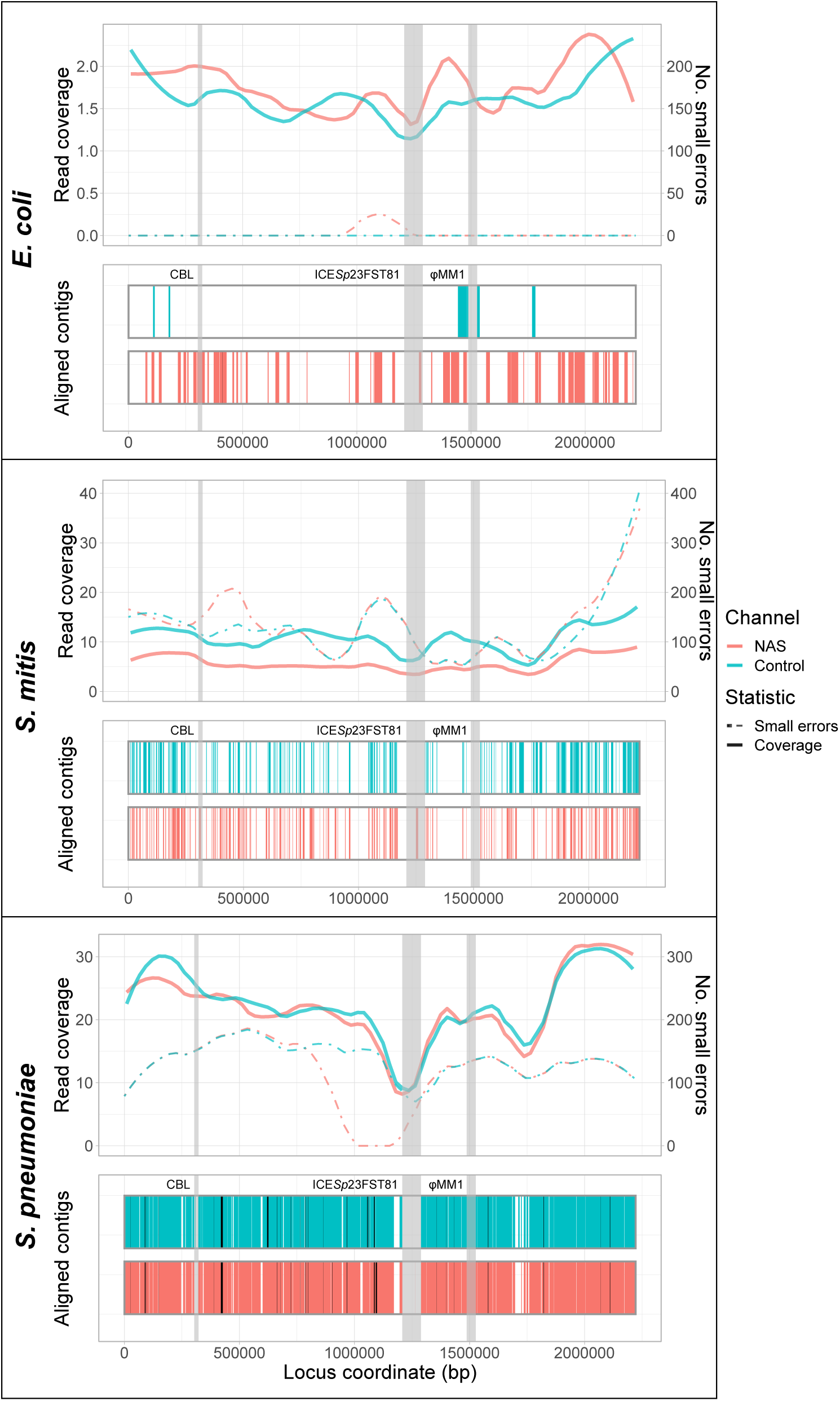
Spn23F whole genome enrichment assembly comparison. Each panel describes a Spn23F assembly generated from 0.1 Spn23F dilutions with each non-target organism. For each panel, the top plot shows the read coverage (solid), defined as the absolute number of bases aligning to a locus, and number of small errors (*≤* 50 bp, dashed), whilst the bottom plot shows aligned contigs (colours) and large errors (black bars, *>* 50 bp) in each assembly. Loci of interest are annotated by grey bars; CBL, as well as ICE*Sp*23FST81 and *ψ*MM1 prophage which are missing in Spn23F (*42*).

For the *E. coli* mixture, the Spn23F whole genome assembly contained very few errors, although read coverage was low and the assembly covered only a small portion of the Spn23F genome (Figure 2, top). The resulting assembly from NAS channels had a greater coverage of aligned contigs than that from control channels, coupled with higher read coverage across the Spn23F genome. The assemblies from the dilution with *S. mitis* had greater overall genome coverage than the equivalent *E. coli* mixture, although the respective aligned contigs were short and contained larger numbers of small errors (Figure 2, middle). Based on read coverage, which was higher in the *S. mitis* mixture over *E. coli* despite Spn23F being at equivalent concentrations, these errors are likely due to incorporation of non-target *S. mitis* reads into Spn23F assemblies, ultimately resulting in mismatches with the reference sequence. The dilution of Spn23F with *S. pneumoniae* R6 produced assemblies with the greatest coverage of aligned contigs, although the assemblies also had large numbers of both small and large errors (Figure 2, bottom). There was also a gap in assemblies at the 23F CBL; *S. pneumoniae* R6 is unencapsulated and so does not possess a CBL, meaning that these assemblies likely contained a large number of non-target *S. pneumoniae* R6 reads. Comparing NAS and control assemblies across all mixtures, both read and assembly coverage were similar for the *S. mitis* and *S. pneumoniae* mixtures between control and NAS channels, whilst for *E. coli* the NAS channels outperformed the control channels. These results highlight the inability of NAS to distinguish between closely-related target and non-target sequences, resulting in lowered enrichment and chimeric assemblies.

### 2.2 NAS can effectively enrich for pneumococcal CBL

To improve enrichment by NAS, we specifically enriched for the pneumococcal CBL, which is generally absent from streptococci other than *S. pneumoniae* (*21*). We sequenced the same library as described in Figure 1, targeting 106 distinct pneumoccocal CBL sequences using NAS (see Section 4.5 for details of sequences), measuring the enrichment of the 23F CBL present in the Spn23F genome. Targeting all known CBL sequences using NAS would be the best approach when serotyping a novel isolate in the field, as this practice would provide the highest probability of detecting and enriching for a previously observed serotype. Most CBL are approximately 20 kb, with a 2.2 Mb genome, and therefore the enrichment values were scaled by 8 *×* 10*^−^*^3^ to account for the smaller target sequence size.

We observed a notable improvement in enrichment when only targeting the 23F CBL, particularly in mixtures containing *S. mitis* and *S. pneumoniae* (Figure 1d, red). For example, for the 4 *×* 10*^−^*^3^ target dilution, CBL enrichment was 7.16 and 5.46, whilst for the 5 *×* 10*^−^*^3^ target dilution, whole genome enrichment was 1.06 and 1.02 for *S. mitis* and *S. pneumoniae* mixtures respectively. For the *E. coli* mixture, the coverage difference between NAS and control channels at the 23F CBL locus was greater when enriching for the whole Spn23F genome than for the 23F CBL, whilst the reverse was true for the *S. mitis* and *S. pneumoniae* (Figure 3). These results indicate that when a non-target species is sufficiently divergent from target species, both whole genome and CBL enrichment are viable means of serotyping (Figure 1d, red). However, directly targeting the CBL boosts NAS performance when non-target species are closely related to the target. Therefore, CBL sequences are sufficiently divergent from the rest of the *S. pneumoniae* genome, as well as other closely related genomes, to be differentiated and enriched for. Overall, CBL enrichment works consistently, independently of the population composition, whilst whole genome enrichment is dependent on concomitant non-target species. Furthermore, 23F DNA was still detectable at the lowest concentration tested, meaning that NAS can enrich for target DNA at concentrations as low as 1 in 10,000 bases (targeting 20 kb of 2.2 Mb pneumococcal genome (*∼* 1%) in a 0.01 dilution) in a mixed sample. Taken together, these results indicate that targeting CBL for identification and serotyping of *S. pneumoniae* is a viable alternative to whole genome enrichment in complex microbial samples.

**Figure 3:**
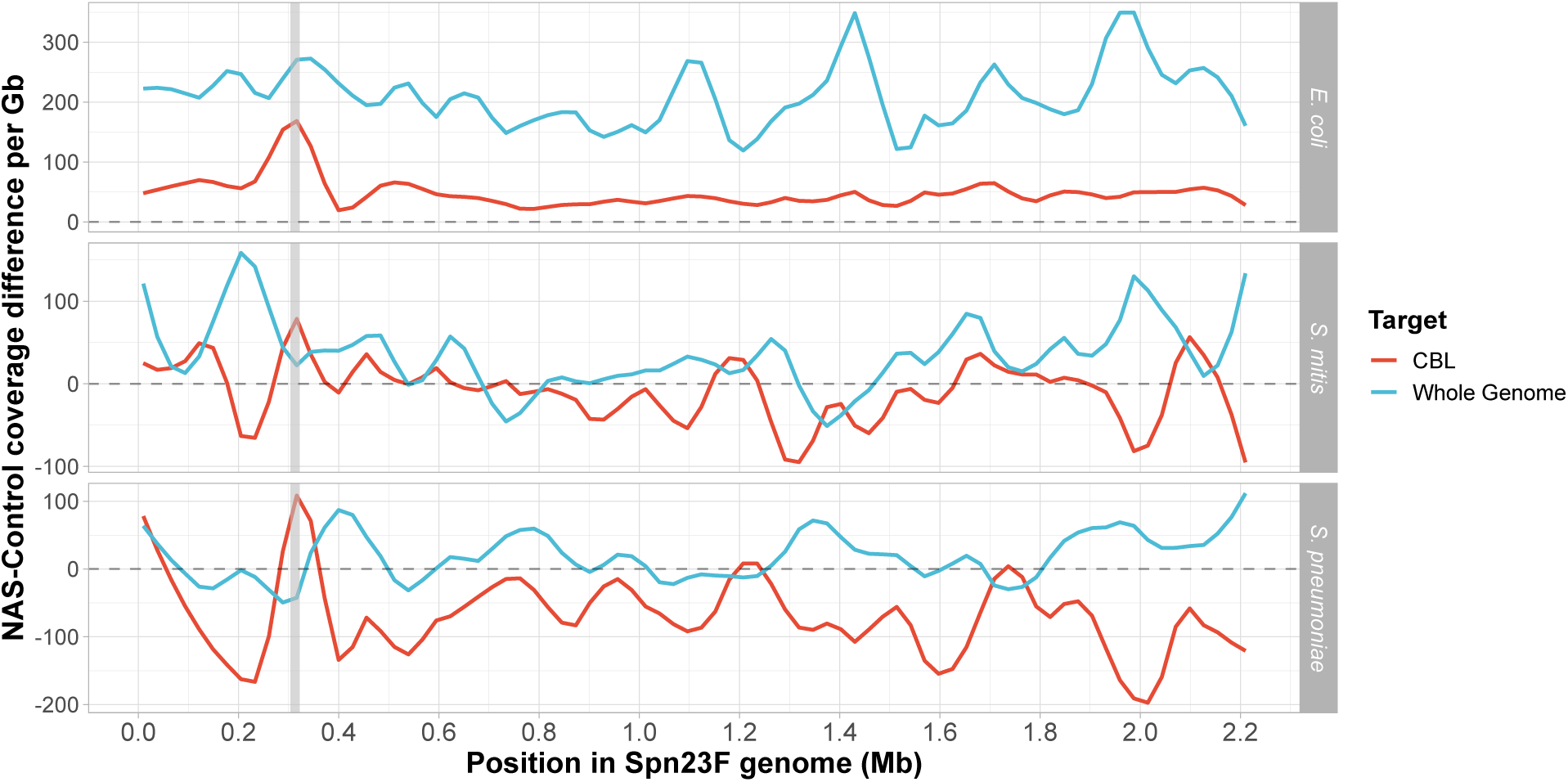
Difference in normalised coverage per locus between NAS and control channels across the Spn23F genome when targeting whole genome (blue) or CBL (red). NAS-control coverage difference per gigabase (Gb) calculated by normalising the read coverage for each locus by the amount of data generated (in Gb) for each respective sample and channel, and then negating the normalised coverage for control channels from NAS channels for each locus. The grey dashed line at 0 indicates equivalent coverage at a given locus between NAS and control channels; *>* 0 indicates NAS channels generated greater coverage, *<* 0 indicates control channels generated greater coverage. Data shown for 0.1 dilutions of Spn23F only. Grey column in each plot highlights 23F CBL locus. Rows show different species for the non-target which was mixed with Spn23F in each sample.

We then generated and compared 23F CBL assemblies as before, focusing on 0.1 Spn23F dilutions, equating to 8 *×* 10*^−^*^4^ 23F CBL DNA (Figure 4). For mixtures containing *E. coli* and *S. mitis*, NAS channels generated more read coverage than control channels, resulting in more complete 23F CBL assemblies containing very few errors. For the mixture containing *S. pneumoniae* R6, 23F CBL assembly was conversely more complete for control channels, likely due to low read counts making the assembly process noisy and leading to patchy coverage for both NAS and control channels (Supplementary Table 6). Overall, NAS improved assembly quality at lower target DNA concentrations over normal sequencing, although low read count made the assembly process more variable.

**Figure 4:**
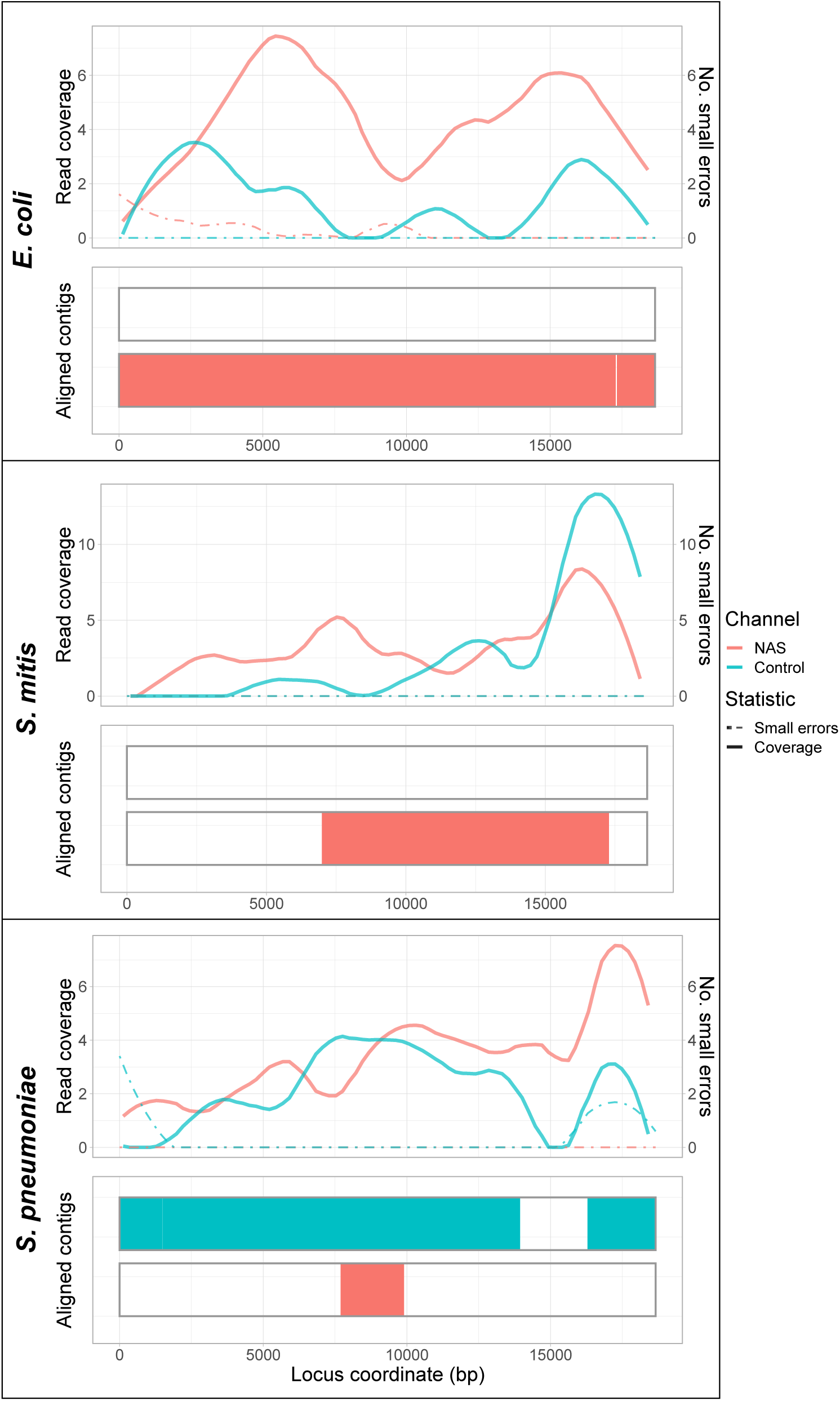
Spn23F CBL enrichment assembly comparison. Each panel describes a 23F CBL assembly generated from 0.1 Spn23F dilutions (8 *×* 10*^−^*^4^ 23F CBL proportion) with each non-target organism. For each panel, the top plot shows the read coverage (solid), defined as the absolute number of bases aligning to a locus, and number of small errors (*≤* 50 bp, dashed), whilst the bottom plot shows the aligned contigs (colours) and large errors (black bars *>* 50 bp) in each assembly.

Previous studies have shown that although NAS increases the proportion of target bases within the read dataset, it may reduce the absolute yield for an equivalent sequencing time (*26*, *31*). In these instances, normal sequencing would give increased coverage of the target genome, and therefore NAS should be avoided. To determine whether this was the case with enrichment of the whole Spn23F genome and 23F CBL, we compared the total number of bases aligning to target sequences across control and NAS channels (Supplementary Figure 25). For whole genome enrichment, the absolute yield was lower for NAS channels on average; however, this difference was not significant. For CBL enrichment, there was a significant increase in absolute yield using NAS (2.17-fold on average, p=0.0049). Therefore, when targeting sequences that are divergent from non-target DNA, NAS increases both proportional and absolute yield.

### 2.3 NAS can simultaneously enrich for multiple pneumococcal CBL in the same mixture

NAS is therefore capable of distinguishing encapsulated *S. pneumoniae* from other streptococci, but effective serotype surveillance requires the identification of multiple serotypes in cases of co-carriage. CBL are highly structurally diverse (*21*), potentially allowing differentiation of multiple CBL in co-carriage by phasing contiguous structural variants using long reads (*43*). To determine whether NAS can differentiate and enrich for multiple CBL sequences, we generated a set of mock communities where Spn23F was mixed in 50:50 proportions with other *S. pneumoniae* strains with different genotypes and serotypes (Figure 5a). We then targeted CBL sequences using NAS; however, we increased the number of times a read can be realigned to the reference sequence before it is rejected (‘maxchunks’ = 4, rather than 0) to determine whether this would improve enrichment of poorly aligned short reads.

**Figure 5:**
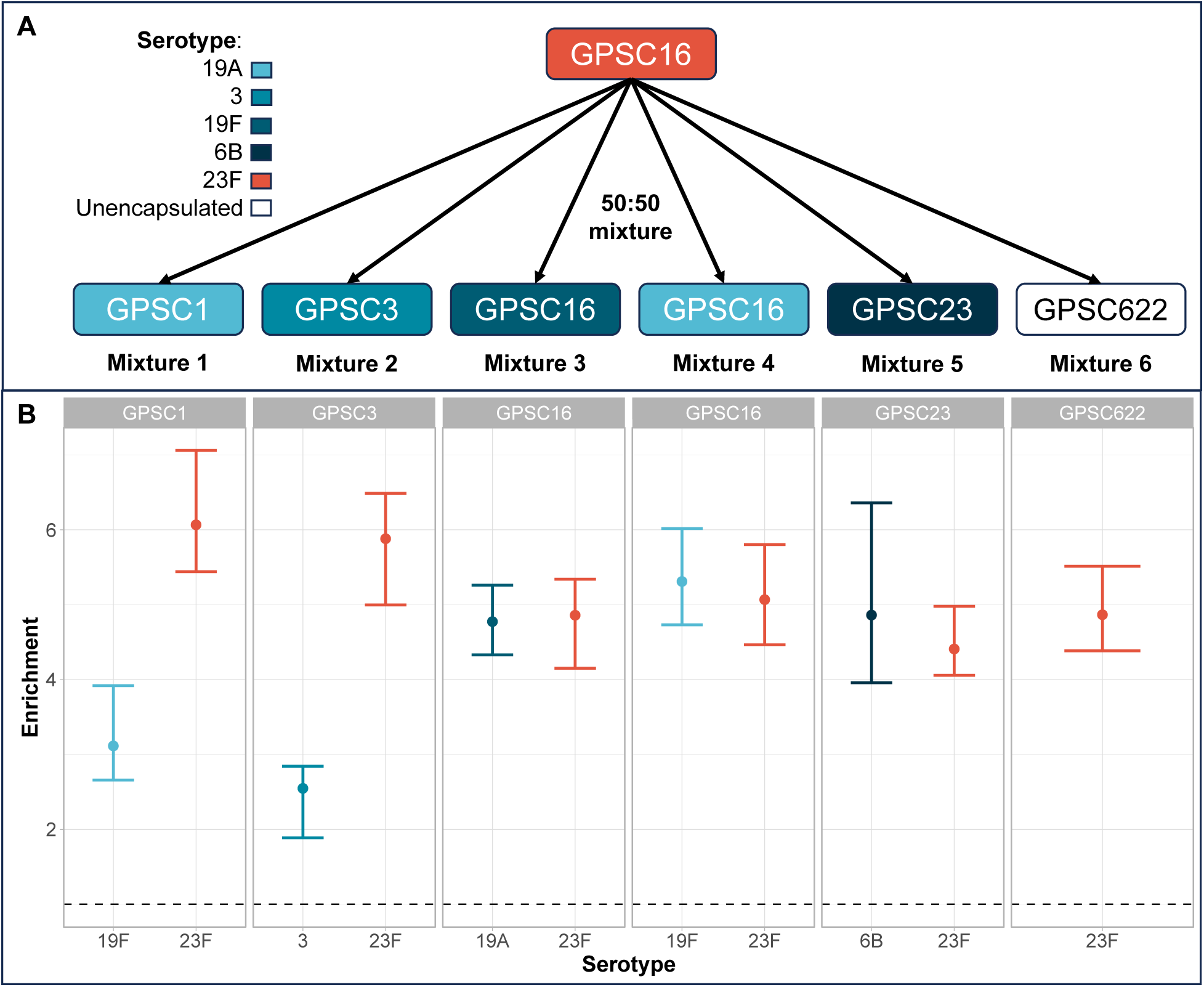
CBL enrichment in mixtures of multiple pneumococci. **a**) Experimental setup. Spn23F DNA (GPSC16, serotype 23F, red) was mixed in 50:50 proportions with other *S. pneumoniae* isolates with different serotypes (given by colour) and genotypes (given by Global Pneumococcal Sequence Cluster, GPSC). **b**) Enrichment of multiple CBL in mixtures. Bar ranges are inter-quartile range of enrichment from 100 bootstrap samples of reads. Data points are observed enrichment values for each CBL per library. X-axis and colour describes the serotype combination of the *S. pneumoniae* isolate mixed with Spn23F, columns describe the GPSC. Dashed line describes enrichment = 1 i.e. no enrichment has occurred.

All CBL sequences were enriched in across all mixtures, independent of serotype or genotype (Figure 5b), with NAS significantly increasing the yield of reads aligning to the CBL locus relative to control channels by 1.9-fold on average (p=9.5 *×* 10*^−^*^7^) (Supplementary Figure 26). Therefore, NAS can be used for targeted sequencing in cases of co-carriage, regardless of respective *S. pneumoniae* serotypes or genotypes. However, CBL enrichment was slightly lower than that observed in mixtures containing a single encapsulated isolate at equivalent concentrations. Comparing the enrichment of the 23F CBL in the 50:50 mixture with the unencapsulated strain, R6, (Figure 5b, GPSC622), with that observed previously (Figure 1d, 4 *×* 10*^−^*^3^ target dilution with *S. pneumoniae*), enrichment was reduced (4.9 vs. 5.6). Therefore, increasing the ‘maxchunks’ had a detrimental impact on enrichment and should kept at zero.

To determine whether NAS improves serotype prediction accuracy in mixed samples, we then analysed reads from mixed samples using PneumoKITy (*18*), a tool for pneumococcal serotype prediction from read data. Serotype predictions were correct for all but mixture 1 using reads from NAS channels, which missed a prediction of 19F (Table 1); however, this serotype was successfully identified in mixture 3. For control channels, three mixtures had incorrect serotype predictions, where either one or both serotypes were missed. For samples where two serotypes were correctly predicted, estimated proportions did not deviate substantially from 50%, the expected values for these mixtures, and were similar between NAS and control channels. Therefore, NAS improved the accuracy of co-carriage detection over normal sequencing.

**Table 1:** Serotype predictions from mixed samples. Each row describes a mixture from Figure 5, with the expected serotypes in the 50:50 mixtures and relative proportions estimated by PneumoKITy (*18*) for reads generated from NAS and control channels. Prediction errors are highlighted in red. ‘None’ represents the unencapsulated isolate.

| Mixture | Expected |  | NAS |  | Control |  |
| --- | --- | --- | --- | --- | --- | --- |
|  | Serotype A | Serotype B | A % | B % | A % | B % |
| 1 | 23F | 19F | 100 | 0 | 0 | 0 |
| 2 | 23F | 3 | 36.4 | 63.6 | 0 | 0 |
| 3 | 23F | 19A | 39.1 | 60.9 | 36.4 | 63.6 |
| 4 | 23F | 19F | 41.2 | 58.8 | 0 | 100 |
| 5 | 23F | 6B | 37.5 | 62.5 | 46.2 | 53.8 |
| 6 | 23F | None | 100 | 0 | 100 | 0 |

### 2.4 Optimising serotyping sensitivity from metagenomes with graph-based alignments using GNASTy

We have shown that pneumococcal CBL sequences are a more suitable target for metagenome-based serotype surveillance than whole genomes. However, high sequence divergence between CBL may limit NAS application in the discovery of previously unobserved serotypes. Although SNPs and short variants can usually be aligned to a divergent reference, larger structural variation, present between CBL of different pneumococcal serotypes, can hinder read alignment when variants are not captured in a reference database (*44*). Such variation can be captured using a pangenome graph, which is a compact representation of multiple linear DNA sequences. These are constructed by merging similar sequences into nodes, with variation between genomes represented by edges. Pangenome graphs provide a means of recapitulating unobserved structural variation, enabling greater flexibility in alignment to capture novel recombinants (*45*) and alignment across assembly gaps (*46*). We therefore explored the application of graph alignment in NAS to enrich for novel *S. pneumoniae* CBL.

We developed and implemented a read-to-graph alignment method to replace the linear alignment method currently used in NAS methods (Figure 6). Our method employs ‘pseudoalignment’, whereby short overlapping nucleotide sequences, known as ‘*k*-mers’, are matched between a read and a de Bruijn graph (DBG), a type of pangenome graph built from matching shared *k*-mers between reference sequences (*47*, *48*). Pseudoalignment is faster than conventional graph alignment, which uses a seed-and-extend approach between a query and reference sequence, and has been used previously in metagenomic read classification (*49*, *50*). We implemented graph psuedoalignment using Bifrost (*51*), which builds coloured compacted DBGs, whereby *k*-mers are ‘coloured’ by their source genomes, with non-branching paths of k-mers ‘compacted’ into sequences known as ‘unitigs’, reducing graph size. We named this method ‘GNASTy’ (Graph-based Nanopore Adaptive Sampling Typing, pronounced ‘nasty’). A detailed description of the GNASTy method is available in the Supplementary Material (Section A.3).

**Figure 6:**
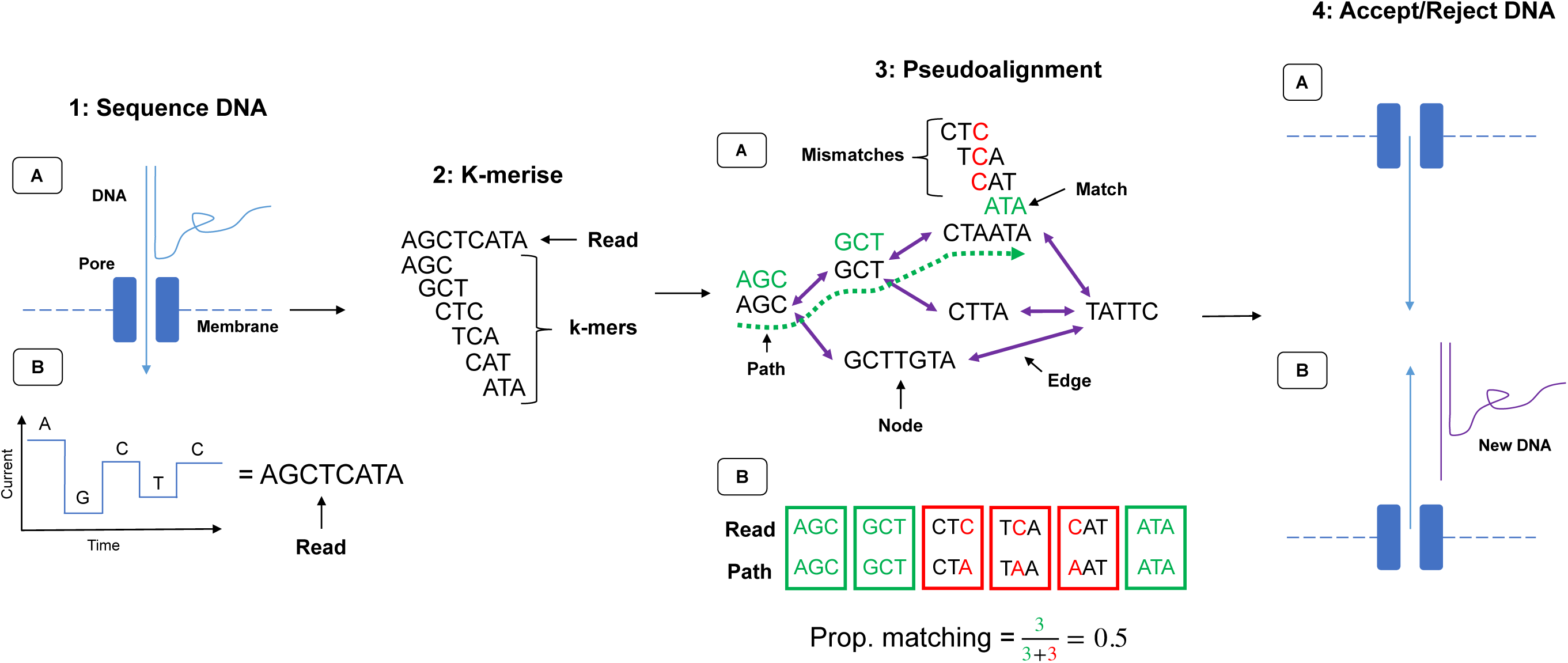
NAS using graph pseudoalignment in GNASTy. (**1a**) The start of a DNA fragment passes through a nanopore, disrupting movement of ions and causing a change in current determined by the base passing through the pore. (**1b**) This current change is used to basecall the read. (**2**) The read is *k*-merised, depending on the *k*-mer size used to build the DBG. (**3a**) The *k*-mers are matched to those in the graph via pseudoalignment, analogous to traversing a hypothetical path (dotted green line). (**3b**) The number of matches (green) and mismatches (red) are used to calculate the proportional number of k-mer matches between the read and the hypothetical path in the graph. (**4a**) If the read surpasses the pre-defined identity threshold, the remainder of the DNA is sequenced. (**4b**) If not, the voltage is reversed across the membrane, pushing the read in the revere direction and freeing the pore to sequence a new DNA fragment.

To benchmark the accuracy of GNASTy against the current linear alignment used during NAS, we generated a simulated dataset of nanopore reads from the Spn23F and *E. coli* DH5-*α* reference genomes. Reads originating from the 23F CBL were classified as target reads, and all others were classified as non-target. We compared classification accuracy of GNASTy against Minimap2 (*52*), the linear aligner used in Readfish (*26*). The alignment index for both methods was generated from the 106 CBL sequences used previously. Graph pseudoalignment using GNASTy was carried out with a *k*-mer size of 19 bp based on simulation sequencing run performance (Supplementary Material, Sections A.4 and A.5), with a percentage identity, defined by minimum read-graph identity or ‘*S*’, of 75% between a read and a path within the DBG for a read to be classified as a target. Additionally, minimum read length was set to 50 bp, with unaligned reads below this length being rejected.

We found that alignment sensitivity was higher for GNASTy than Minimap2, whilst specificity was similar between the two methods (Table 2). Therefore, Minimap2 had a greater tendency to incorrectly reject target reads than GNASTy, whilst correct rejection of non-target reads by GNASTy was only slightly lower than Minimap2.

**Table 2:** Alignment accuracy comparison between graph pseudoalignment using GNASTy (denoted by ‘Graph’) and Minimap2. Sensitivity is defined as *TP/*(*TP* + *FN*), specificity is defined as *TN/*(*TN* + *FP*).

| Tool | No. TP | No. FN | No. TN | No. FP | Sensitivity | Specificity |
| --- | --- | --- | --- | --- | --- | --- |
| Graph | 1149 | 495 | 494790 | 3566 | 0.699 | 0.993 |
| Minimap2 | 713 | 931 | 497514 | 842 | 0.434 | 0.998 |

When comparing computation speed between the two methods, Minimap2 outperformed Bifrost/GNASTy during index generation and read alignment. Minimap2 was 30-fold faster at index generation than Bifrost and used 4.5-fold less memory, although Bifrost generated an index 2-fold smaller than Minimap2 (Supplementary Table 5). This is an upfront cost not relevant during sequencing. Per-read alignment times for GNASTy were notably higher than those for Minimap2 (Supplementary Figure 27). For GNASTy, all reads were individually aligned in less than 1*/*8*^th^* of a millisecond, equivalent to sequencing 0.056 bases assuming a rate of 450 bases sequenced per second (*26*). If 512 reads were aligned in a single chunk (the maximum number of reads that could be generated at once on a MinION), this would be equivalent to an additional 29 bases being sequenced per pore before a decision is made to accept or reject each read. Therefore we tested whether GNASTy’s greater sensitivity for detecting target reads, at the cost of slower rejection of non-target reads, would increase the enrichment of CBL sequences.

### 2.5 Graph-based alignment facilitates the discovery of novel CBL

We investigated whether GNASTy’s graph representation of CBL variation would enable it to discover and enrich novel CBL variants more accurately than conventional NAS. To evaluate this, we tested whether GNASTy could outperform linear alignment when the target sequence was not present in a reference database. We used the 106 CBL sequences used in Section 2.2 as a reference database, removing the 23F CBL, along with all closely related CBL (cluster two from Mavroidi *et al.* (*53*)). We then sequenced the same samples used previously (Figure 1a and b), and calculated enrichment of the 23F CBL. These experiments used V14 sequencing chemistry, which generates reads faster than now-discontinued V12 chemistry, but has similar read quality (*54*). Minimap2 was compared with graph psuedoalignment in GNASTy with *k* = 19 and minimum read length of 50 bp as before. We tested two percentage identity thresholds, *S* = 75% and *S* = 90%, to understand the effect of increasing graph pseudoalignment stringency on enrichment.

Results showed that enrichment could be achieved by all NAS methods and parameter values, although graph pseudoalignment (*S* = 75%) performed best, with equivalent or higher enrichment than Minimap2 across all samples (Figure 7). Therefore, the slower read alignment speed of graph pseudoalignment compared to Minimap2 did not have a large enough effect to negatively impact enrichment. In addition to enrichment, absolute yield of 23F bases was significantly increased using graph pseudoalignment (*S* = 75%) relative to control channels. These achieved a mean yield increase of 2.75 fold (p=9.8 *×* 10*^−^*^4^), which was greater than for Minimap2, which achieved a mean yield increase of 2.0 fold (p=2.4 *×* 10*^−^*^3^) (Supplementary Figure 28). Furthermore, graph pseudoalignment with *S* = 75% performed similarly to Minimap2 when the 23F CBL was included in the reference database, meaning that GNASTy can be used as a direct replacement for Minimap2 for NAS (Supplementary Material, Section A.6). Graph pseudoalignment with *S* = 90% performed worst of the three methods, resulting in lower enrichment and reduced absolute yield which was not significantly different from control channels.

**Figure 7:**
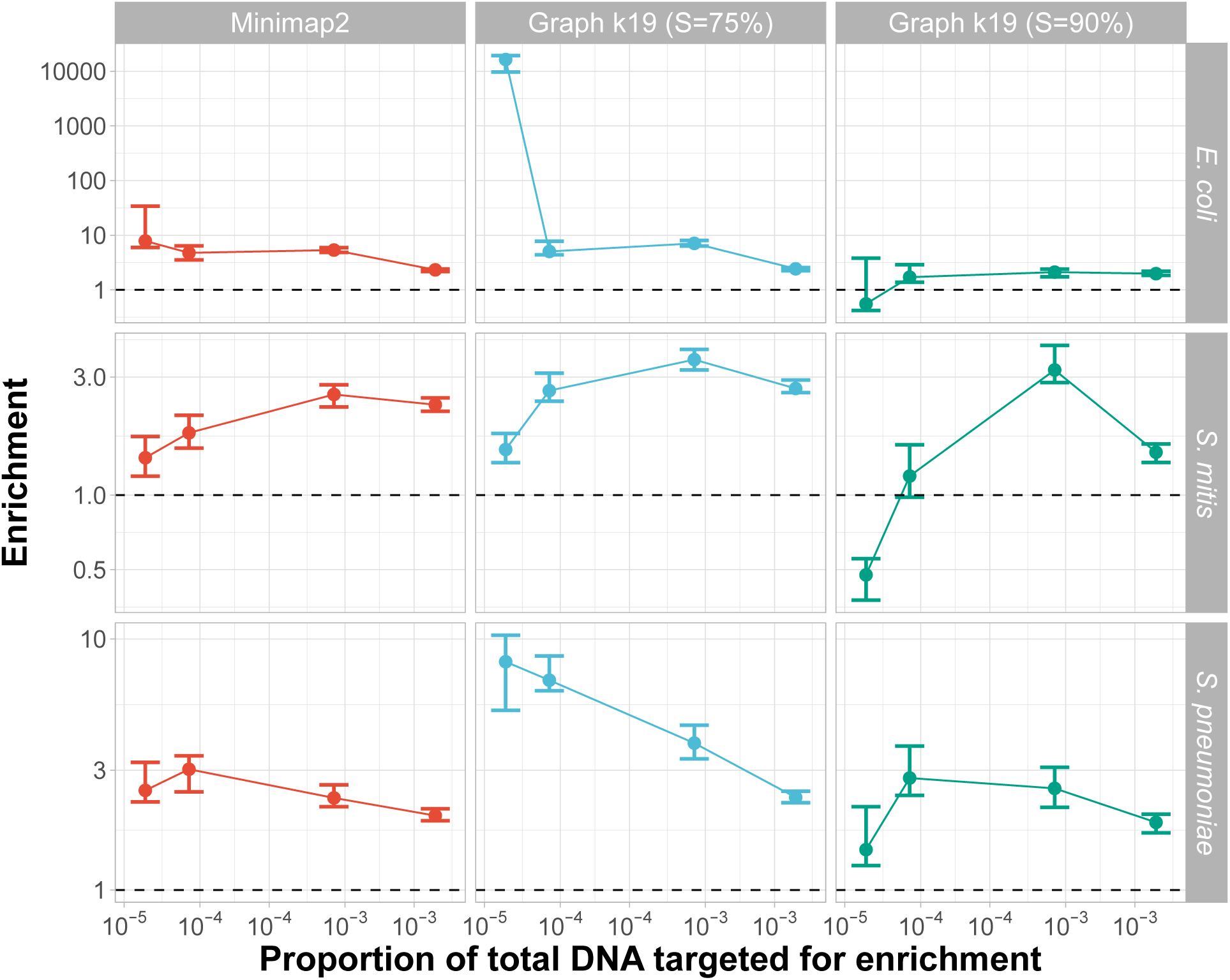
Enrichment comparison of 23F CBL at different concentrations of target between Minimap2 and graph pseudoalignment in GNASTy when aligning to a partial CBL reference database. Bar ranges are inter-quartile range of enrichment from 100 bootstrap samples of reads. Data points connected by lines are observed enrichment values for each library, with solid lines connecting the same genome diluted at different concentrations. Rows describe the non-target species in the mixture, columns describe the alignment method used. To plot on a log scale, all enrichment values had 0.01 added to them. Horizontal dashed line describes enrichment = 1 i.e. no enrichment has occurred.

The increased enrichment we observed for graph pseudoalignment in GNASTy over Minimap2 may be due to biased over-sequencing of a specific position in the 23F CBL, rather than even coverage of the entire 23F CBL. To enable accurate assembly of the full 23F CBL, coverage should ideally be increased evenly across the target sequence, rather than over-represented in specific regions. Comparison of normalised coverage across the full 23F CBL for Minimap2 and graph pseudoalignment showed similar read coverage variability across the 23F CBL all three methods (Supplementary Figure 29). For example, at the highest target proportion (4 *×* 10*^−^*^3^), there were coverage spikes at both ends of the CBL locus for all three methods. Therefore, enrichment achieved by linear alignment can be explained by mapping of the start of read to shared regions at the ends of CBL (Supplementary Figures 18). Coverage for both Minimap2 and graph pseudoalignment fell in the center of the CBL, which is particularly notable at 4 *×* 10*^−^*^3^ target dilutions. At the lowest target dilutions (4 *×* 10*^−^*^5^), a spike in coverage can be observed at the 18 kb position in the 23F CBL for mixtures containing *S. mitis* and *S. pneumoniae*. As the non-target isolates *S. mitis* SK142 and *S. pneumoniae* R6 both contain *aliA* homologues (*55*, *56*), which is also found at the end boundary of all *S. pneumoniae* CBL (*21*), this peak can be attributed to non-specific enrichment of a gene common to streptococci. However, coverage was equivalent or higher for graph pseudoalignment with *S* = 75% over Minimap2 across all target concentrations and non-target species. In summary, although NAS can enrich for novel serotypes using linear alignment, using GNASTy increases NAS sensitivity.

Next, we compared the ability for Minimap2 and GNASTy to correctly identify the 23F serotype in the mixtures using PneumoKITy serotype calls (Supplementary Figure 30). We compared the proportion of the 23F CBL reference sequence covered by the reads, which is used by PneumoKITy as a proxy for serotype prediction confidence (*18*). Minimap2 and graph pseudoalignment (*S* = 75%) performed similarly, with reads from NAS channels providing more support for the 23F CBL call than for controls in all cases. Even at low target concentrations (*≤* 8 *×* 10*^−^*^5^), these alignment methods were still able to identify 23F as the most likely serotype, with the exception of the mixture with *S. pneumoniae* R6, where serotype 2 was predicted to be the most likely serotype. *S. pneumoniae* R6 is derived from a serotype 2 strain via deletion of its respective CBL (*57*); however, presence of CBL flanking sequences in *S. pneumoniae* R6, as described above, likely lead to false detection of serotype 2 CBL.

We then compared assemblies of the 23F CBL across the three alignment methods. We chose samples containing 0.1 Spn23F dilution with *S. mitis* to mimic carriage of a single isolate (Supplementary Figure 31). For all alignment methods, read coverage was higher for NAS channels than for control channels, although graph pseudoalignment (*S* = 90%) had the lowest absolute coverage for both channel types. Despite variation in coverage, all assemblies covered a majority of the CBL with minimal errors of any kind. Assembly completeness was similar between control and NAS channels, except at the right end of the CBL, where Minimap2 and graph pseudoalignment (*S* = 90%) were unable to generate an aligning contig. This effect was also observed in Supplementary Figure 24, and may be due to uneven local read coverage affecting assembly contiguity, as Metaflye expects uniform coverage for individual strain genomes (*40*). Additionally, two central regions (*∼* 7.5 kb and *∼* 12 kb), and a small region in the 18 kb end of the CBL, were missing in the control assembly for graph pseudoalignment (*S* = 75%). However, these were correctly identified when reads were enriched with GNASTy. When GNASTy was run with the suboptimally high alignment specificity parameter (*S* = 90%), the NAS assembly was missing a single region (*∼* 7.5 kb) present in the control assembly. Therefore, whilst assemblies were largely similar between NAS and control channels, these small differences indicate higher graph pseudoalignment stringency slightly lowered assembly quality compared to the control, whilst greater sensitivity for CBL reads improved assembly quality.

### 2.6 Graph-based alignment enriches CBL in complex samples mimicking the nasopharynx microbiome

Previous experiments demonstrated GNASTy was capable of enriching for CBL from simple mixtures. Therefore, we tested whether the method was also effective with more realistic microbial compositions that would be observed in the nasopharynx or oral cavity. We used samples containing a mixed culture generated from nasopharyngeal swabs, spiking in Spn23F as before. As there was no ground truth for these samples, it was unknown whether *S. pneumoniae* strains were already present prior to spiking. Spn23F DNA was added to give a final proportion of 0.1 of total DNA in each sample, reflecting typically observed *S. pneumoniae* prevalences in the nasopharynx (*58*), resulting in a final 23F CBL DNA proportion of 8 *×* 10*^−^*^4^. Libraries were run without size selection, as we observed a detrimental affect on extracted DNA yield with mixed culture samples which did not affect single isolate samples (Supplementary Figure 32). NAS was conducted using graph pseudoalignment (*k* = 19%, *S* = 75%) with GNASTy using a database containing all 106 CBL sequences, including the 23F CBL. As a control, a sample containing Spn23F mixed with *S. pneumoniae* R6 without size selection at 0.1 and 0.5 proportions was also run, and compared with equivalent samples with size selection.

Enrichment of the 23F CBL was achieved for all mixed culture samples (Figure 8), with all samples outperforming the unselected *S. pneumoniae* R6 sample at the equivalent concentration (8 *×* 10*^−^*^4^). Size selection had a notable positive impact on enrichment in the *S. pneumoniae* R6 mixtures, increasing enrichment from 1.4-2.0-fold to 3.9-4.4-fold for 8 *×* 10*^−^*^4^ and 4 *×* 10*^−^*^3^ target proportions, respectively. This was consistent with the method’s performance with the simpler DNA mixtures (Supplementary Figure 6). Therefore the lowered performance of GNASTy with these complex samples relative to the simpler mixtures was a consequence of the lack of size selection during DNA sample preparation. This also explains the similar target yield between NAS and control channels (Supplementary Figure 33). Therefore, we advise size selection be used where possible to boost NAS efficiency, although it may not be suitable in all cases due to high DNA yield loss.

**Figure 8:**
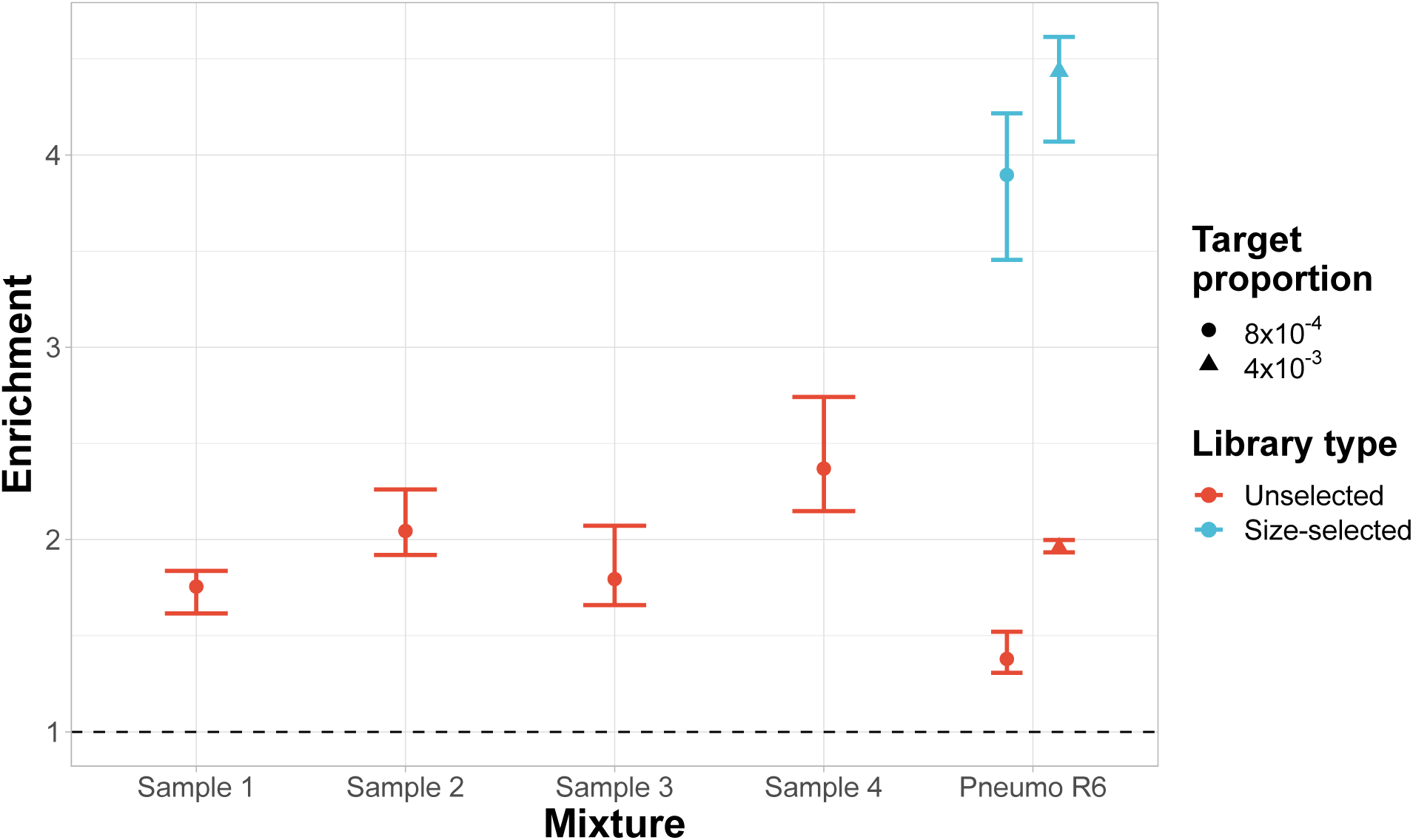
Enrichment of 23F CBL across samples containing mixed cultures from nasopharyngeal swabs. All nasopharyngeal samples (denoted ‘Sample X’) were run without size selection, with control samples containing Spn23F mixed with *S. pneumoniae* R6 (denoted ‘Pneumo R6’) without size selection at 0.1 and 0.5 proportions (8 *×* 10*^−^*^4^ and 4 *×* 10*^−^*^3^ 23F CBL DNA proportions respectively) run alongside. Equivalent control samples from a run with size selection are plotted for comparison. Bar ranges are inter-quartile range of enrichment from 100 bootstrap samples of reads. Data points are observed enrichment values for each library. Dashed line describes enrichment = 1 i.e. no enrichment has occurred.

The 23F CBL was identified as the most likely CBL in Samples 1 and 2, as well as the *S. pneumoniae* R6 mixtures by PneumoKITy, although Sample 1 had evidence of co-carriage (Supplementary Figure 34). In Samples 3 and 4, the 23F CBL was not identified as the most likely serotype, although the proportion of the reference 23F CBL sequence matched was above PneumoKITy’s confidence cutoff (70%) for reads originating from NAS channels, meaning that these samples were identified as containing a mixture of serotypes. Notably, the prediction for the 23F CBL did not meet the confidence cutoff for reads from control channels in sample 4, meaning that NAS enabled the detection of a low level secondary serotype that would have otherwise been missed. Assembly quality was similar between NAS and control channels, although NAS channel read coverage was equivalent or higher for most samples (Supplementary Figure 35). Nevertheless, the read coverage was sufficiently high (>20) to enable assembly in principle, which likely could be achieved by algorithms capable of correcting for the uneven read coverage across the 23F CBL in these datasets. Overall, we have shown that NAS, employing graph pseudoalignment with our tool GNASTy, can be used to enrich for *S. pneumoniae* CBL in mock communities resembling real nasopharyngeal samples.

## 3 Discussion

The complex population dynamics of *S. pneumoniae* reflect the antigenic and genetic diversity of the species, and the serotype replacement that has been driven by the widespread use of serotype-specific vaccines (*8*, *10*, *13*). Current methods for *S. pneumoniae* serotype surveillance are limited either by their requirement for application to individual colonies (e.g. WGS or antiserum agglutination), reducing their sensitivity for detecting co-carriage, or by being restricted to only the serotypes known at the time the assay was designed (e.g. PCR genotyping, microarrays). Deep sequencing has revealed the need for sensitive sequencing-based assays that can detect novel serotypes (*9*, *13*). However, such datasets are resource intensive to produce, and necessitate substantial pre-existing infrastructure. Therefore, simpler, quicker and cheaper surveillance methods are required to provide a comprehensive view of pneumococcal serotype diversity and prevalence to inform public health strategies.

In this work, we developed a pangenome graph-based alignment method, GNASTy, that meets all of these criteria, and can be applied to mixed pneumococcal populations in complex samples, mimicking a culture-free workflow. This represents a substantial improvement over the standard NAS in the direct detection and serotyping of pneumococci. We found that when targeting a whole *S. pneumoniae* genome in the presence of closely-related non-target species, enrichment was reduced, as alignment lacks the specificity to distinguish between target and non-target reads. However, we showed that targeting the *S. pneumoniae* CBL increases both enrichment and yield of NAS due to the specificity of the sequence to *S. pneumoniae*. We show this enables the detection, enrichment and serotyping of multiple CBL simultaneously, and can detect a serotype that only comprises 1% of the sample. Furthermore, we showed that GNASTy enables greater enrichment of novel CBL over linear alignment, used in the current implementation of NAS, and is therefore capable of discovering rare or previously-unknown serotypes.

Direct detection of pneumococcal CBL using GNASTy also promises to be a scalable surveillance method. NAS does not rely on culture, reducing the time required to generate a result compared to WGS or agglutination assays. Library construction takes a few hours, and NAS required one day of sequencing on a portable MinION device, which can be shortened if sufficient read coverage is reached during sequencing. Although this is slower than genotypic methods, such as PCR and microarrays which take a few hours (*22*), GNASTy is capable of identifying the full range of serotypes, with any updates to the method achievable through simply updating the reference database, without alterations to the laboratory protocol. Furthermore, NAS is fully portable and simple enough to be run with limited equipment, requiring only bench-top apparatus such as a centrifuge, thermocycler (*25*) and high performance laptop, as used here. NAS also currently has lower entry costs compared to other sequencing technologies such as Illumina or PacBio, with increased target yield enabling sample multiplexing to reduce per-sample costs. We observed *>* 2-fold increases in target yield compared to standard sequencing using NAS, enabling twice the number of samples to be run on the same flow cell to achieve the same target coverage, therefore halving per-sample costs. Lowered costs, coupled with the portability of Nanopore sequencing, make NAS attractive for applications in low resource settings where pneumoccocal disease burdens are highest (*59*). Overall, GNASTy provides a balance of accuracy, simplicity and cost-effectiveness, making it well-suited for routine pneumococcal surveillance in both high and low resource settings.

A key improvement of targeted sequencing over shotgun metagenomic sequencing is the increased limit of detection, meaning more rare sequences can be identified. We showed that NAS can increase the proportion of target DNA more than 10-fold over that of the control channels, based on enrichment by composition, when enriching for CBL sequences at concentrations *<* 0.01%, in line with previous evaluations of NAS efficiency (*27*, *31*). This improved CBL coverage achieved by NAS was further shown to increase the sensitivity of DNA-based serotyping in samples mimicking co-carriage. Finally, we showed that NAS can enrich for CBL DNA in samples mimicking the complexity of the nasopharyngeal microbiome, and improves serotyping accuracy over normal sequencing. However, in the mock communities used throughout this work, our measures of *S. pneumoniae* abundance were relative proportions based on measures from observed nasopharyngeal microbiomes (*58*). We did not convert target concentrations into absolute concentration values (e.g. in ng/µL of target DNA), as sequencing sensitivity will be dependent on the number of bases generated per sample, which itself is contingent on multiplexing and variability in DNA loading onto the flowcell.

Sample throughput is a key factor when determining the suitability of methods for routine pneumococcal serotype surveillance. For a method to be viable for routine surveillance, it must offer the ability to batch process, or multiplex, samples with sufficient sensitivity to detect true positive cases. Based on our experience, we recommend sequencing between 12-24 samples on a single flowcell to provide sufficient coverage to detect *S. pneumoniae* DNA, whilst reducing the cost per sample through multiplexing. Such relatively small batches are practical in routine surveillance applications, contrasting with the hundreds of samples that need be multiplexed for higher-throughput sequencing methods to be maximally cost effective.

The current limitations to GNASTy primarily represent the challenges of optimising DNA sample preparation. The mock communities tested did not contain human reads; however, oral and nasopharyngeal samples often contain substantial host DNA, which will ultimately impact target yield. Therefore, GNASTy will require further optimisation to include host DNA depletion, for which suitable laboratory methods are available (*60*, *61*). One potential solution is to use NAS to deplete human DNA as off-target reads, which is possible if the human genome were supplied as a database of unwanted sequence. Such depletion sequencing is more effective than targeted sequencing when sampling the full bacterial diversity of a sample (*29*). Hence GNASTy may have additional utility when applied to culture-free nasopharyngeal samples using depletion sequencing.

NAS has the potential to enable accurate, direct and relatively inexpensive *S. pneumoniae* surveillance. However, this work highlights the current limitations of enriching for low-abundance species with NAS in mixtures containing closely-related taxa, and the suboptimal sensitivity for identifying loci that are not present in the target database. We have developed and tested NAS for the detection and serotyping of *S. pneumoniae* in complex samples, providing methodological recommendations and a novel pangenome graph-based method for use by public health researchers, which we hope will improve access to accurate *S. pneumoniae* surveillance in low resource settings. GNASTy promises to be a powerful method both for routine epidemiology, and novel serotype discovery.

## 4 Materials and Methods

### 4.1 Isolate and sample acquisition

All isolate bacterial strains used in this work included: *E. coli* DH5-*α*, *Moraxella catarrhalis* 0193-3, *Haemophilus influenzae* 0456-2, *S. mitis* SK142, *Streptococcus oralis* SK23, *S. pneumoniae* ATCC 706669 (referred to as ‘Spn23F’, GPSC16, serotype 23F), *S. pneumoniae* R6 (GPSC622, unencapsulated), *S. pneumoniae* 110.58 (GPSC81, unencapsulated), *S. pneumoniae* MalM6 (GPSC16, serotype 19F), *S. pneumoniae* 8140 (GPSC16, serotype 19A), *S. pneumoniae* Tw01-0057 (GPSC1, serotype 19F), *S. pneumoniae* K13-0810 (GPSC23, serotype 6B), and *S. pneumoniae* 99-4038 (GPSC3, serotype 3).

Nasopharyngeal swab samples were chosen from a collection originating from a study of mother-infant pairs in the Maela camp for refugees in Thailand (*62*, *63*). This research complied with all relevant ethical regulations, and was approved by the Ethics Committee of The Faculty of Tropical Medicine, Mahidol University, Thailand (MUTM-2009-306), and by the Oxford Tropical Research Ethics Committee, Oxford University, UK (OXTREC-031-06). All women gave written informed consent to participate in the study. Individuals did not receive monetary compensation for their participation.

### 4.2 Bacterial culture and DNA extraction

For culture, glycerol stocks containing bacterial isolates and nasopharyngeal swab (referred to as ‘mixed culture’) samples were inoculated in 10 mL of Todd-Hewitt broth (Oxoid, UK) and 2% yeast extract (Sigma-Aldrich, UK) and cultured overnight overnight at 35°C in 5% CO_2_ atmosphere. For culture of *M. catarrhalis* and *H. influenzae*, 3 mM hemin (X factor) and 22.5 mM nicotinamide-adenine-dinucleotide (NAD, V factor) were also added to respective inocula. Liquid cultures of *M. catarrhalis*, *H. influenzae* and *E. coli* and mixed cultures were incubated with shaking at 150 rpm. Following incubation, the inocula were centrifuged at 16000 *g* for 10 min, with supernatent being discarded to obtain cell pellets.

DNA was extracted from cell pellets using the Wizard Genomic DNA Extraction Kit (Promega UK, Catalogue number: A1120). For *S. pneumoniae* isolates and mixed culture samples, cell pellets were re-suspended in 480 µL 50 mM EDTA, before addition of 120 µL freshly prepared lysozyme (30 mg/mL). The solution was incubated at 37°C for 60 min, before centrifugation at 16000 *g* for 2 min, with the supernatent being discarded. For all isolates, 600 µL nuclei lysis solution was added to pellets and incubated at 80°C for 5 min. Three µL RNase solution was then added and incubated at 37 °C for 15 min, before cooling to room temperature. 50 µL of 20 mg/mL recombinant Proteinase K solution (Life Technologies, Catalogue number: AM2548) was then added, with the sample being incubated at 55°C for one hour. Two hundred µL protein precipitation solution was then added and incubated on ice for 5 min, before solutions were centrifuged at 16000 *g*, and the supernatent transferred to a clean tube. Six hundred µL of room temperature 100% isopropanol was then added to the supernatent and centrifuged at 16000 *g*, with the supernatent being discarded. Six hundred µL of room temperature 70% ethanol was then added to the pellet and mixed to resuspend the pellet. The solutions were centrifuged at 16000 *g*, with the supernatent being discarded, and the pellets were allowed to air-dry for 15 min. DNA pellets were then resuspended in 150 µL DNA rehydration solution.

Extracted DNA was size selected using the SRE XS kit (Pacbio US, Catalogue number: SKU 102-208-200) following manufacturer’s instructions to remove fragments *<* 10 kb in length.

### 4.3 DNA Quality control

Extracted DNA was quantified using a dsDNA broad-range assay kit (Catalogue number: Q32850) on the Qubit 3 fluorimeter (ThermoFisher Scientific UK) following manufacturer’s instructions. DNA was also sized using a Genomic DNA ScreenTape Assay (Catalogue numbers: 5067-5366 [Reagents], 5067-5365 [Screentape]) on the TapeStation 2200 system (Agilent UK) following manufacturer’s instructions. DNA samples with modal peaks *>* 45 kb were carried forward for library construction and sequencing.

### 4.4 Library construction

Library construction was conducted using the native barcoding kits (ONT UK, Catalogue numbers: SQK-NBD112.24 [V12 chemistry], SQK-NBD114.24 [V14 chemistry]) following manufacturer’s instructions. Briefly, 400 ng DNA was aliquoted per barcoded sample for end and single-strand nick repair using NEBNext Ultra II End repair/dA-tailing Module and NEBNext FFPE Repair Mix (New England Biolabs UK, Catalogue numbers: M6630S, E7546S), with samples then being cleaned using AMPure XP Beads (Beckman Coulter UK) and 70% or 80% ethanol for V12 and V14 chemistry respectively. Barcode ligation followed, using the barcodes provided and the NEB Blunt/TA Ligase Master Mix (New England Biolabs UK, Catalogue number: M0367L), with samples then being pooled together and cleaned as before. Finally, adapter ligation was conducted using the NEBNext Quick Ligation Module (New England Biolabs UK, Catalogue number: E6056S), with the library cleaned using AMPure XP Beads and the long-fragment buffer provided with the ONT library construction kit. Libraries were loaded onto MIN112 or MIN114 flowcells for V12 and V14 chemistries respectively.

### 4.5 Sequencing and adaptive sampling

All analysis scripts and CBL reference sequences used in this work are available on Zenodo (*64*) (see Section 5.4). Genbank reference sequence accession numbers for whole genome assemblies include: *E. coli* DH5-*α* (NZ_JRYM01000009.1), *M. catarrhalis* (NZ_CP018059.1), *H. influenzae* (NZ_CP007470.1), *S. mitis* SK142 (NZ_JYGP01000001.1), *S. oralis* SK23 (NZ_LR134336.1), *S. pneumoniae* Spn23F (FM211187.1), *S. pneumoniae* R6 (NC_003098.1) and *S. pneumoniae* 110.58 (CP007593.1).

Sequencing was conducted using a MinION Mk1B instrument and a Dell Mobile Precision 7560 with an Intel Xeon processor and 128 GB RAM, and a NVIDIA RTX A5000 GPU with 16 GB GPU RAM running MinKNOW v22.12.7 (ONT UK). Local GPU base-calling was conducted using Guppy v6.4.6 (ONT UK) with the fast base-calling model and reads were rejected immediately if they did not align to the reference genome by setting ‘maxchunks’ to 0 in the Readfish ‘.yaml’ file. For each new library, a control sequencing run was conducted for 1 hour with no adaptive sampling with bulk capture, providing a ‘recording’ for simulation playback.

Adaptive sampling was carried out using Readfish v0.0.10dev2 (*26*). Graph pseudoalignment was carried out using a custom fork from the Readfish GitHub repository (*65*) (see Section 5.4). Readfish was installed using the ‘readfish.yml’ file present in the GitHub repository by running the command ‘conda create -f readfish.yml’. During sequencing, Readfish was run using the command ‘sudo runuser minknow -c ‘/path/to/readfish targets --device [device] --experiment-name [name] --channels 1-256 --toml /path/to/toml --logfile [logfile] --port 9502 --graph [True/False] --align_threshold [threshold] --len_cutoff [cutoff]”.

Adaptive sampling was used on channels 1-256 of the flowcell, with the remaining 256 channels run as controls without adaptive sampling. Linear alignment for adaptive sampling was carried out using Mappy v2.24 (https://pypi.org/project/mappy/). Metadata for all sequencing runs and samples, including the number of bases generated and aligned, the number of reads generated and aligned, and calculated enrichment, are available in Supplementary File 1. This file also links each sequencing run archived on the European Nucleotide Archive (*66*) to individual barcoded samples.

### 4.6 Enrichment analysis

Enrichment analysis was based on analysis performed by Martin *et al.* (*31*). Enrichment by composition of the target *x*, denoted by *E_x_*, was calculated as described in Equation 1. Each flowcell was bioinformatically split into two halves; one half contained channels (a segment of a flowcell containing a nanopore) which were ‘adaptive’ (using NAS), the other half contained channels which were ‘controls’ (not using NAS). *E_x_*was calculated as the fold increase in the proportion of read bases aligning to *x* in adaptive channels, *a*, versus control channels, *c*:

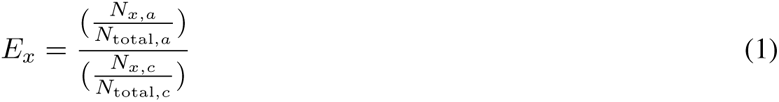

where *N_x_*is the number of bases aligning to *x*, and *N*_total_ is the total bases in either adaptive or control channels. Using enrichment by composition enables results to be compared across sequencing runs, which may vary in the amount of data generated.

To calculate enrichment post-sequencing, all reads, including those passing and failing the phred-score filter (Q-score *≥* 8), were aligned to a reference sequences using Mappy v2.24 using the custom script ‘analyse_RU.py’. Reads were aligned to specific reference sequences based on known isolates present within each sample (‘-t <target>’). All reads were used to avoid any potential biases introduced by read filtering, such as flow cell spatial effects, in the calculation of enrichment. Reads were split by channel (‘-c 1-256’) to identify which reads were sequenced under NAS (channels 1-256) or control (channels 257-512) conditions. Reads aligning above a specified minimum identity threshold (84% identity within the aligned block, ‘-p 0.84’) were assigned as target reads, with the highest-identity alignment for multi-mapping reads being taken as the only alignment. Only regions of reads aligning to a reference sequence were included in enrichment calculations. Quartiles were generated from 100 bootstrapped samples of aligned reads (‘-bs 100’).

### 4.7 Serotype prediction

Serotype prediction was conducted using a customised version of PneumoKITy which can be run using a single fastq file, as opposed to paired fastq files as in the original version, available on Zenodo (https://doi.org/10.5281/zenodo.10590659) (*67*). Reads were split using a custom script (split_by_channel.py) to generate files for reads sequenced under adaptive (channels 1-256) and control (channels 257-512) conditions (‘--channels 1-256’). PneumoKITy was run in ‘mix’ mode using a minimum median-multiplicity value of 4 (‘-n 4’) and a minimum kmer percentage of 85% (‘-p 85’) for reference CBL sequence matching.

### 4.8 Assembly and quality control

All reads were first re-basecalled using Guppy v6.4.6 with the super-high accuracy model using the following command: ‘guppy_basecaller --compress_fastq --input_path [input_path] --save_path [output_path] --config dna_r10.4.1_e8.2_400bps_sup.cfg --device cuda:0 --recursive --barcode_kits SQK-NBD114-24 --enable_trim_barcodes --trim_adapters --trim_primers’. Reads were then assembled using MetaFlye v2.9.2 (*40*) in ‘--nano-raw’ mode. We did not use the high accuracy ‘--nano-hq’ mode, as testing showed this was too stringent and resulted in no assembly being generated for some samples. Assembly quality was then analysed using Inspector v1.2 (*41*), with reads mapped to respective assemblies, and assembly contigs mapped to respective reference sequences to identify errors. Errors were identified in contigs *≥* 50 bp in length (‘--min_contig_length_assemblyerror 50’, ‘--min_contig_length 50’). BED files generated by Inspector, containing contig alignment and error positions on respective reference sequences, were visualised using a custom script (‘plotting_scripts /generate_linear_assembly_plot.R’). Read alignment for coverage analysis was conducted using the custom script, ‘analyse_coverage.py’, using the original reads basecalled using Guppy’s fast basecalling model. Alignment and read parsing settings were the same as ‘analyse_RU.py’ described above. All alignment was carried out using Mappy v2.24. Assembly statistics are available in Supplementary File 2.

### 4.9 Nanopore sequencing simulation and analysis

Simulations of nanopore sequencing runs were conducted using bulk capture recordings from previous sequencing runs, as described on the Readfish GitHub repository (https://github.com/LooseLab/readfish). Results were analysed using a custom script (analyse_unblocks.py). This script aligns reads to a specified target sequence using Mappy v2.24 and classifies them as either accepted or rejected by the adaptive sampling process. Reads that align to a target sequence and were accepted or rejected are classified as true positives and false negatives respectively. Reads that did not align to a target sequence and were accepted or rejected are classified as false positives and true negatives respectively. For all experiments described here, the reference sequence was the Spn23F chromosome (‘--ref data/cps/sequences/SP_ATCC700669.fasta’) and the target sequence was the 23F CBL sequence (‘--loci data/cps/split_cps/23F.fa’).

For benchmarking of alignment speed, a bespoke simulation model was generated using Nanosim-H v1.1.0.2 (*68*, *69*). Model training used FASTQ files from a V14 chemistry nanopore sequencing run containing 50%-50% dilutions of *S. pneumoniae* Spn23F and *E. coli* DH5-*α*, and their respective reference sequences (‘nanosim-h-train -i training_reads.fasta reference/genome.fasta output’). Using this model, 500,000 simulated nanopore reads were generated (‘nanosim-h -n 500000 -p output reference_genome.fasta’). Simulated reads were then split into true positive and true negative reads based on whether they originated from the 23F CBL using the custom script split_simulated.py, which parses reads simulated by Nanosim-H based on their original locus. Reads overlapping by at least 50 bp with the 23F CBL (position 303558-322212 bp within the Spn23F chromosome) were classified as true positives (‘--pos 303558-322212 --min-overlap 50’). CBL sequences (updated_cps.fasta, N=106) were indexed using Minimap2 v2.26 (*52*) and Bifrost v1.2.0 (*51*) with *k* = 19. The time taken to align all 500,000 simulated reads for Mappy v2.24 and graph pseudoalignment was measured using a custom script (simulate_readuntil.py) which parses the start of each read, with length defined by Poisson sampling (‘--avg-poi 180’, based on Payne *et al.* (*26*)). This fragment is then aligned using both Mappy and graph pseudoalignment, with alignment timed using the Python ‘timeit’ module. Graph pseudoalignment was run using minimum read identity 75% (--id 0.75) and minimum read length 50 bp (‘--min-len 50’). Mappy was run with default parameters. Alignment accuracy was measured based on whether a read was accepted or rejected, depending on whether it originated from the 23F CBL or not. Comparisons were carried out on a server cluster with dual processor x86-64 nodes, running CentOS v8.2.

### 4.10 Pseudoalignment simulation

Pseudoalignment simulations proceeds as follows. A specified number of target and non-target sequences of given lengths are generated by random sampling of DNA bases. Constituent k-mers of these sequences are then generated, and reads with specified mutation rates are simulated from target sequences. Read k-mers are then matched back to the respective target and non-target k-mers sets, enabling calculation of recall and precision respectively. The code for this process can be found in the ‘kmer_simulation.R’ script.

## Abbreviations

AMR: Antimicrobial Resistance
CBL: Capsular Biosynthetic Locus/Loci
DBG: de Bruijn Graph
Gb: Gigabase(s)
GNASTy: Graph-based Nanopore Adaptive Sampling Typing
GPSC: Global Pneumococcal Sequence Cluster
IPD: Invasive pneumococcal disease
Mb: Megabase(s)
NAS: Nanopore adaptive sampling
ONT: Oxford Nanopore Technologies
PCR: Polymerase Chain Reaction
PCV: Polysaccharide conjugate vaccine
Spn23F: *Streptococcus pneumoniae* isolate ATCC 700669 serotype 23F
WGS: Whole Genome Sequencing

## 5 Acknowledgments

## 5.1 Funding

S.T.H. was funded by the MRC Centre for Global Infectious Disease Analysis (studentship grant ref.: MR/S502388/1), jointly funded by the UK Medical Research Council (MRC) and the UK Foreign, Commonwealth and Development Office (FCDO), under the MRC/FCDO Concordat agreement and is also part of the EDCTP2 program supported by the European Union. S.T.H. acknowledges funding from the MRC Centre for Global Infectious Disease Analysis (reference MR/X020258/1), funded by the UK Medical Research Council (MRC). This UK funded award is carried out in the frame of the Global Health EDCTP3 Joint Undertaking. N.J.C. and J.A.L. were funded by the UK Medical Research Council and Department for International Development (grants MR/R015600/1 and MR/T016434/1). N.J.C. was also supported by a Sir Henry Dale fellowship jointly funded by Wellcome and the Royal Society (grant 104169/Z/14/A). J.A.L. and S.T.H. were also supported by the European Molecular Biology Laboratory. P.T. was funded by the Wellcome Trust (grants 083735 and 220211). For the purpose of open access, the authors have applied a Creative Commons Attribution (CC BY) license to any Author Accepted Manuscript version arising from this submission.

## 5.2 Author contributions

Conceptualization: S.T.H., N.J.C., and J.A.L. Methodology: S.T.H., N.J.C., and J.A.L. Software: S.T.H. Validation: S.T.H., B.F., Y.F. Formal analysis: S.T.H. Investigation: S.T.H. Resources: S.T.H., P.T, N.J.C. and J.A.L. Data curation: S.T.H. Writing - original draft: S.T.H. Writing - review and editing: All authors. Visualization: S.T.H. Supervision: N.J.C. and J.A.L. Project Administration: S.T.H., N.J.C. and J.A.L. Funding acquisition: S.T.H., P.T., N.J.C., and J.A.L.

## 5.3 Competing interests

The authors declare no competing interests.

## 5.4 Data and materials availability

All analysis scripts used in this work are available on Zenodo (https://doi.org/10.5281/zenodo.10581200) (*64*). This repository also contains 106 *S. pneumoniae* CBL sequences and associated sources (updated_cps.fasta, updated December 19*^th^* 2022) used as reference sequences for NAS. Code for GNASTy is also available on Zenodo (https://doi.org/10.5281/zenodo.10581132) (*65*).

All sequence datasets generated and analysed in this study are available on the European Nucleotide Archive (http://www.ebi.ac.uk/ena) under accession number PRJEB72455 (*66*).

## 6 Supplementary Materials

### A Supplementary Text

#### A.1 Optimising alignment identity threshold for calculation of enrichment

To calculate enrichment, reads are aligned to the Spn23F reference twice: once during NAS, and once in post-sequencing analysis using a custom script (see Section 4.6). Therefore, calculation of enrichment is dependent on the accuracy of read alignment both during NAS and in post-sequencing analysis. Alignment accuracy during NAS is dependent on the base-caller and aligner which are set *a priori*. However, in post-sequencing analysis, alignment accuracy is dictated by the minimum alignment identity between reads and the reference sequence. Minimum alignment identity controls whether a read aligning to the reference sequence is accepted as a true target read. A cutoff that is too low will incorrectly identify non-target reads as target, whilst one that is too high may remove target reads that have high numbers of errors, with both scenarios resulting in lower estimates of enrichment.

We tested the effect of varying the minimum alignment identity on the calculated Spn23F enrichment across all mixed samples, in order to set a cutoff to be used throughout this work. We generated mock communities containing mixtures of genomic DNA of *S. pneumoniae* strain Spn23F (*39*) with that of closely and distantly related non-target non-target species. Spn23F DNA was mixed with DNA from species from different phyla (*E. coli* and *M. catarrhalis* + *H. influenzae*), the same genus but different species (*S. mitis* and *S. oralis*) and the same species (*S. pneumoniae* R6 and *S. pneumoniae* 110.58) (Supplementary Table 1). Mixture of Spn23F with *M. catarrhalis* + *H. influenzae* was used to simulate the composition of the nasopharynx, where co-carriage with these three species is common (*70–72*). Titrations ranged from 0.001-0.5 proportions (0.1% *−* 50%) of Spn23F DNA, to test the effect of lower target inputs on enrichment. Sequencing was carried out using the V12 sequencing chemistry using the Guppy v5.0.15 base-caller.

We found that enrichment was insensitive to minimum alignment identity in the range 80-90% (Supplementary Figure 1). At *>* 90% minimum alignment identity, enrichment was more variable (Supplementary Figure 2) and resulted in depletion of Spn23F DNA in many samples between 92-100%, where enrichment falls below one which should be avoided. This effect was independent of concentration or contaminating species, and is indicative of rejection of correct Spn23F reads containing sequencing errors. Based on these results, a cutoff of 84% minimum alignment identity was used for all further analysis, as this is equivalent to the minimum phred score (Q8) used by the Guppy base-caller in fast mode to identify failed reads.

**Supplementary Table 1:** Experimental set-up of Spn23F whole genome enrichment. Spn23F DNA was diluted in proportion with non-target DNA from members of different genera (Non-*Streptococcus*), the same genera (*Streptococcus*) and the same species (*S. pneumoniae*).

| Non-target type | Non-target species | Spn23F proportion |
| --- | --- | --- |
| Non-*Streptococcus* | *E. coli* | 0.001 (0.1%) |
|  |  | 0.01 (1%) |
|  |  | 0.1 (10%) |
|  |  | 0.5 (50%) |
|  | *M. catarrhalis* + *H. influenzae* | 0.01 (1%) |
|  |  | 0.1 (10%) |
|  |  | 0.5 (50%) |
| *Streptococcus* | *S. mitis* | 0.01 (1%) |
|  |  | 0.1 (10%) |
|  |  | 0.5 (50%) |
|  | *S. oralis* | 0.01 (1%) |
|  |  | 0.1 (10%) |
|  |  | 0.5 (50%) |
| *Streptococcus pneumoniae* | *S. pneumoniae* R6 | 0.001 (0.1%) |
|  |  | 0.01 (1%) |
|  |  | 0.1 (10%) |
|  |  | 0.5 (50%) |
|  | *S. pneumoniae* 110.58 | 0.01 (1%) |
|  |  | 0.1 (10%) |
|  |  | 0.5 (50%) |

**Supplementary Figure 1:**
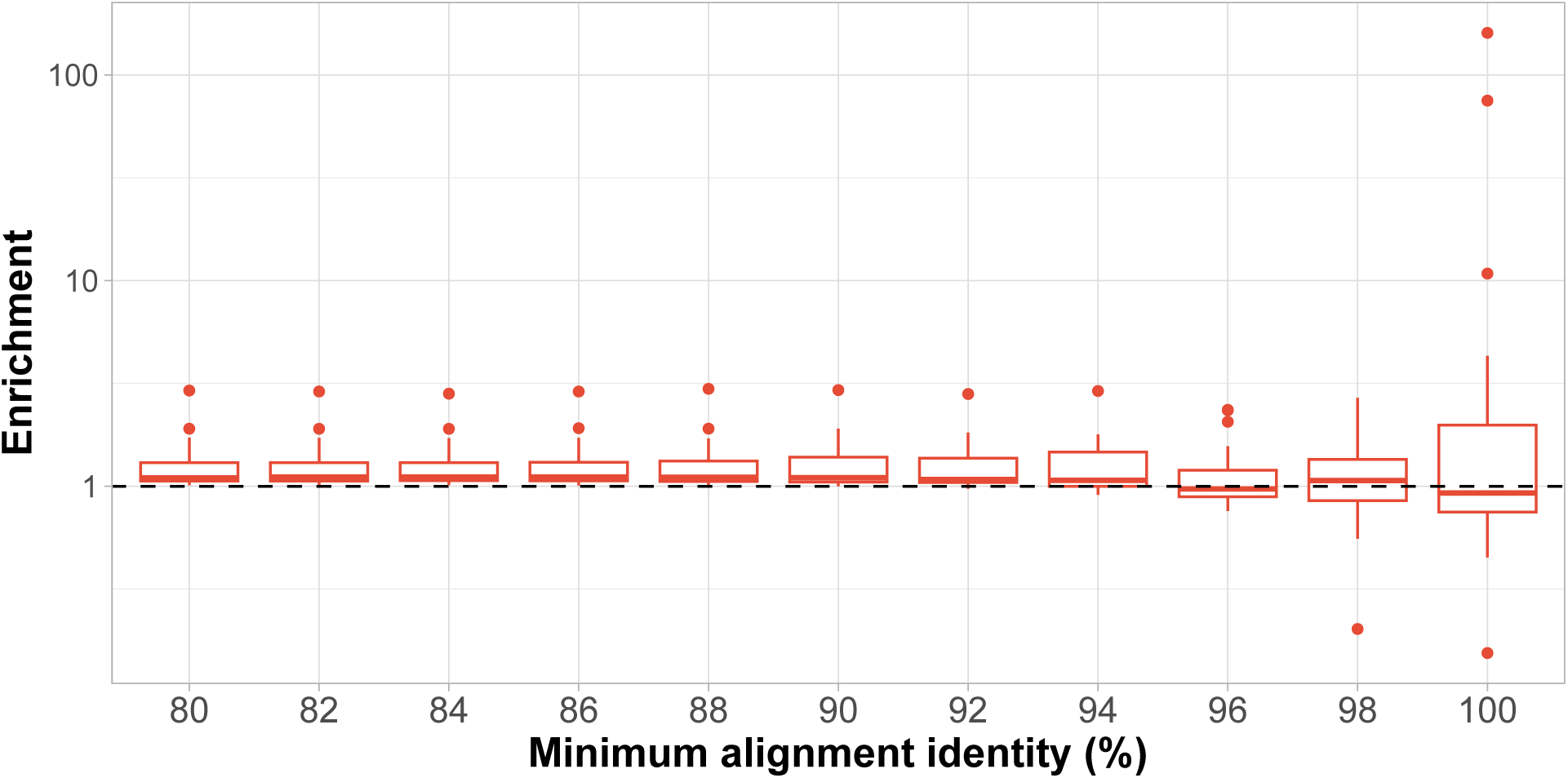
Effect of minimum alignment identity on Spn23F whole genome enrichment. Boxplots represent distributions of enrichment values of Spn23F whole genome in all samples. Dashed line describes enrichment = 1 i.e. no enrichment has occurred.

**Supplementary Figure 2:**
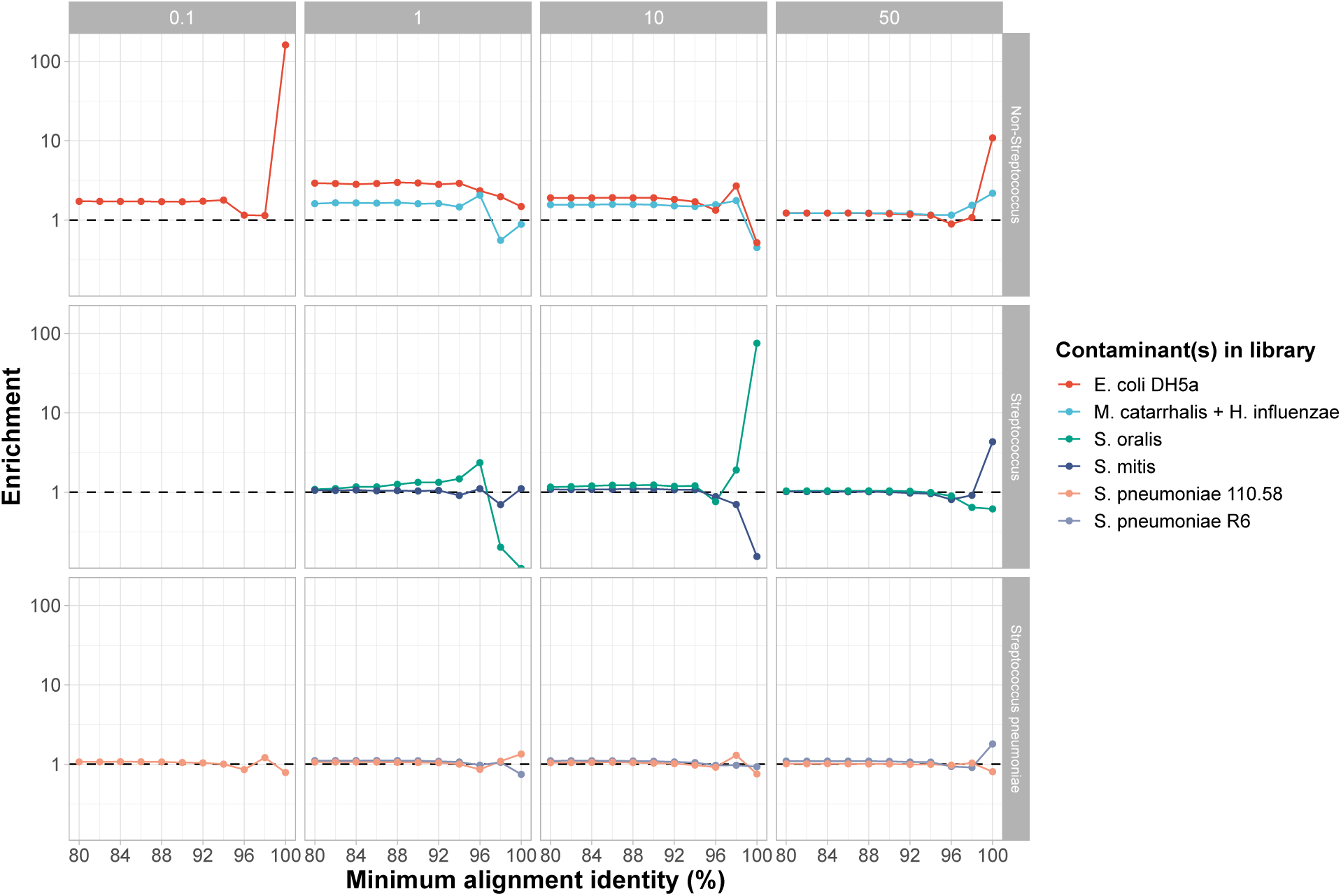
Effect of minimum alignment identity on Spn23F whole genome enrichment across sample concentrations and contaminating species. Lines connect the same sample with different minimum alignment identity thresholds. Columns indicate the % Spn23F DNA in the sample, rows indicate the non-target type. Colours describe the species of the non-target species. No library was sequenced at 0.1% Spn23F DNA with a *Streptococcus* non-target species. Dashed line describes enrichment = 1 i.e. no enrichment has occurred.

#### A.2 Removal of short DNA fragments from libraries using size selection increases target enrichment

Short DNA fragments are sequenced too quickly to be rejected by NAS, resulting in reduced target enrichment if many short non-target sequences are present in a sample. Martin *et al.* (*31*) showed that removal of short fragments improves both enrichment and absolute yield of target DNA. To determine whether this effect could be reproduced, size selection was used on all libraries in Supplementary Table 1, in which DNA fragments *<* 10 kb were removed from each sample.

Comparison of read length distributions before and after size selection highlighted read length proportions were shifted to higher values for accepted reads in adaptive channels and all reads in control channels (Supplementary Figure 3). Mean and median read lengths were still below 10 kb (Supplementary Table 2), indicating size selection was effective but imperfect. The size of rejected reads was similar between both experimental conditions, indicating that size selection had no impact on rejection speed, with faster read rejection expected to result in shorter rejected reads (*26*). size selection had a small but significant effect on enrichment (Supplementary Figure 4), increasing average enrichment from 1.31 to 1.5. Overall, size selection increased the size of accepted reads, although the observed effect on enrichment was small.

**Supplementary Table 2:**
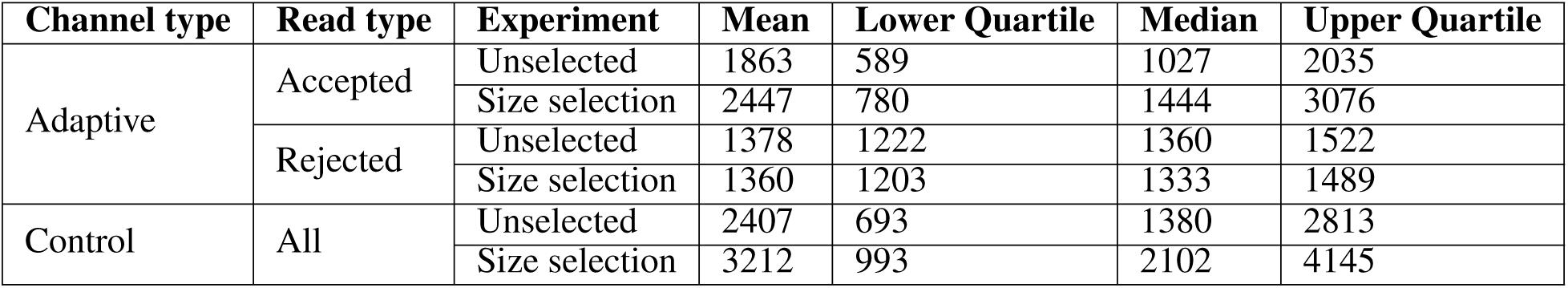
Read length statistics for Spn23F whole genome enrichment. All statistics are in basepairs.

**Supplementary Figure 3:**
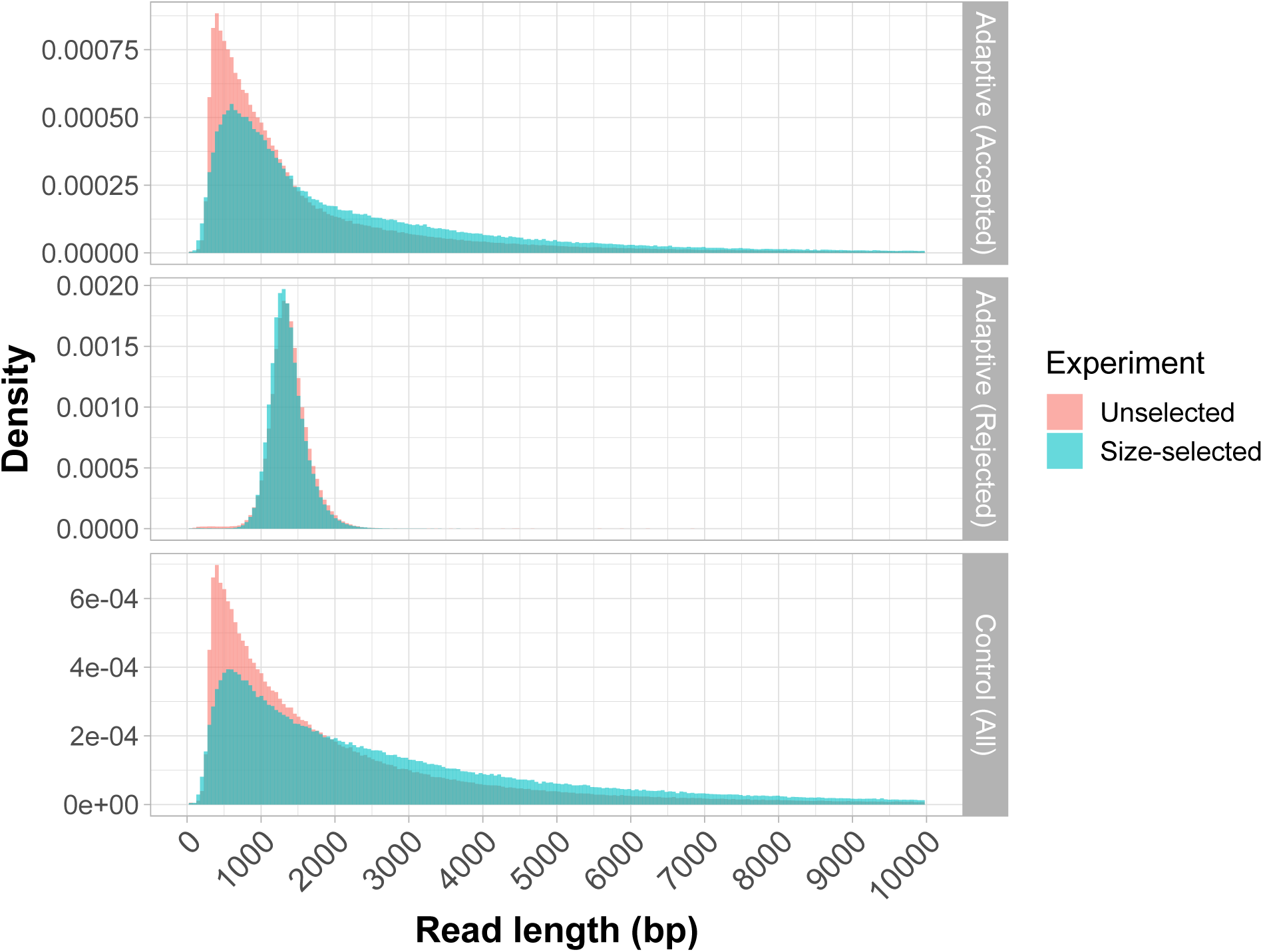
Effect of size selection on read length distributions for Spn23F whole genome enrichment. Histograms for reads accepted (**top**) and rejected (**middle**) by NAS in adaptive channels, and all reads in control channels (**bottom**).

**Supplementary Figure 4:**
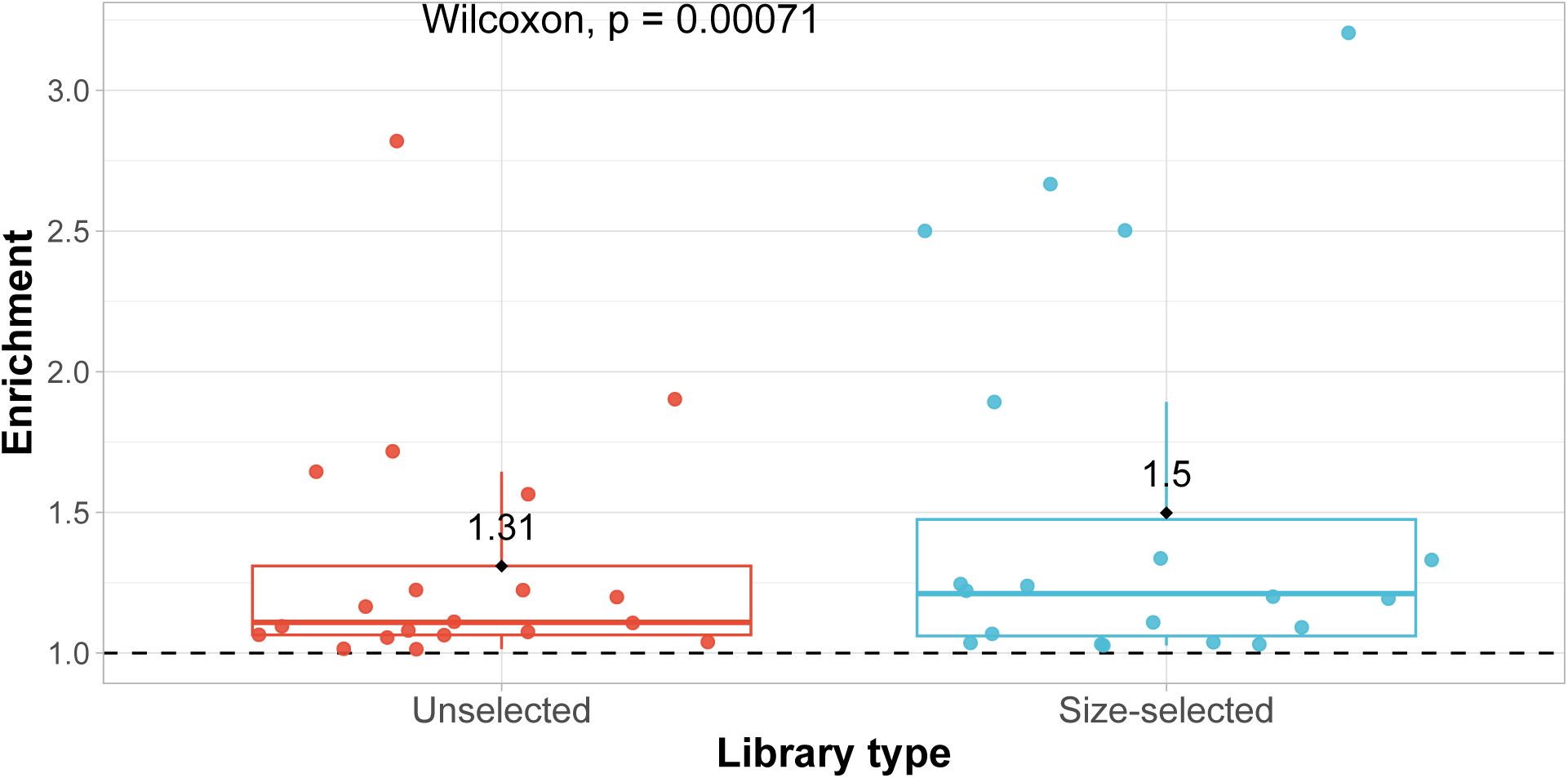
Enrichment comparison of Spn23F genome with and without size selection, which removed DNA fragments *<* 10 kb in length. Each data point represents the enrichment of Spn23F in a single barcoded library. Dashed line describes enrichment = 1 i.e. no enrichment has occurred. Black points highlight distributions means. Distributions were compared using a paired Wilcoxon test.

We additionally tested the effectiveness of size selection on enrichment of CBL. Previously, Viehweger *et al.* (*33*) showed that enrichment of loci, in this case AMR genes, was possible using NAS, although performance was dependent on the library fragment length. Paradoxically, the authors suggested that when enriching for genes, libraries with a greater proportion of short fragments provide better enrichment, in contrast to our observation that size selection improves enrichment. Viehweger *et al.* (*33*) postulated that this was due to the higher probability of a short target locus being found in the middle of a longer read, rather than at the start, which is the region aligned during NAS. In longer DNA fragments, it is therefore more likely that the start region of a read will align outside of the target locus, and will therefore be incorrectly rejected, despite the target sequence being present later in the read. CBL contain *∼* 20 genes, and so are substantially longer than individual genes. Therefore it is unknown whether read length has the same effect on CBL enrichment.

To determine the effectiveness of CBL enrichment with size selection, libraries containing single isolate DNA from *S. pneumoniae* strains were generated, as well as mixtures with Spn23F DNA with a variety of serotypes and genotypes (Supplementary Table 3). Strains with the same serotype but different genotypes, and strains with the same strain background but different serotypes, were included to determine whether CBL are sufficiently different from the rest of the genome and each other to be effectively enriched. Sequencing was carried out using the Guppy v6.3.8 base-caller, which has higher base-calling speed over Guppy v5.0.15 used in the previous section according to the ONT release notes, with V12 sequencing chemistry.

**Supplementary Table 3:** Experimental set-up of CBL enrichment. All strains are members of the *S. pneumoniae* species. Samples with 50% proportion were diluted with Spn23F DNA.

| Strain name | Strain genotype | Serotype | Proportion |
| --- | --- | --- | --- |
| Spn23F | GPSC16 | 23F | 100% |
| 8140 | GPSC16 | 19A | 100% |
|  |  |  | 50% |
| Mal M6 | GPSC16 | 19F | 100% |
|  |  |  | 50% |
| Tw01_0057 | GPSC1 | 19F | 100% |
|  |  |  | 50% |
| 99_4038 | GPSC3 | 03 | 100% |
|  |  |  | 50% |
| K13_0810 | GPSC23 | 06B | 100% |
|  |  |  | 50% |
| R6 | GPSC622 | NT | 50% |

We observed longer reads in size-selected than unselected libraries for accepted reads in adaptive channels, and all reads in control channels (Supplementary Figure 5, Supplementary Table 4), as seen before with whole genome enrichment. However, accepted and rejected reads in adaptive channels for CBL enrichment were notably shorter than those in whole genome enrichment, whilst control channel read lengths were similar (Supplementary Table 2). This effect may be due to accepting of shorter reads when aligning to shorter targets, as observed in Viehweger *et al.* (*33*). Size selection also had a positive significant effect on enrichment when targeting CBL, increasing the average enrichment from 3.4 to 4.8 (Supplementary Figure 6).

**Supplementary Figure 5:**
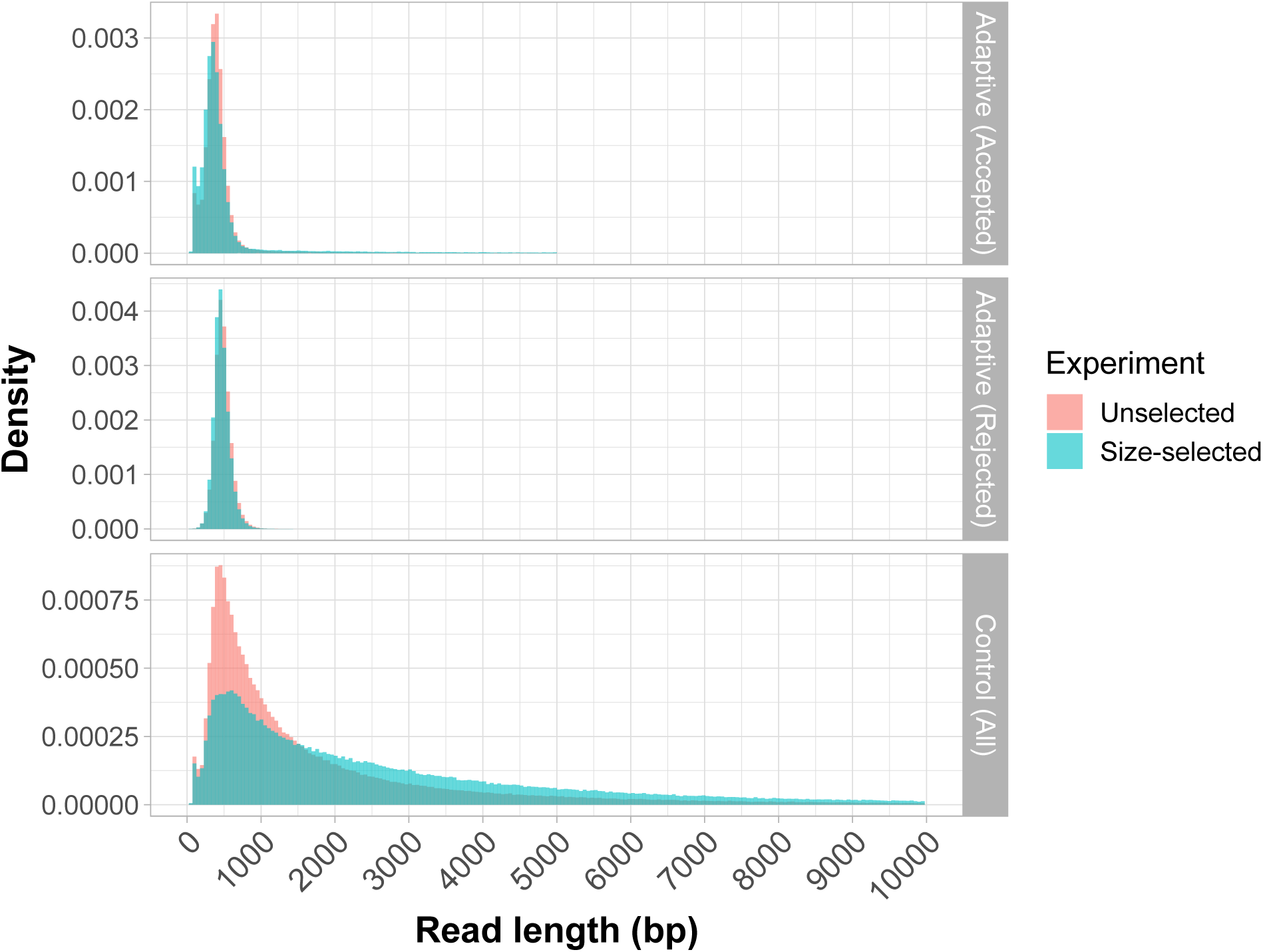
Effect of size selection on read length distributions for CBL enrichment. Histograms for accepted (**top**) and rejected reads (**middle**) in adaptive channels, and all reads in control channels (**bottom**).

Read alignment to the Spn23F genome for the Spn23F-only sample revealed that adaptive channels had lower coverage across genome compared to control channels, with the exception of the CBL locus (located at *∼* 0.3 Mbp) where coverage reached *∼* 4 *−* 5 fold higher than the rest of the genome (Supplementary Figure 7). Moreover, size selection increased CBL normalised coverage by *∼* 1.5 fold over that of the unselected library, whilst coverage of the remainder of the genome was similar between the two libraries. The positive effect of size selection on 23F CBL enrichment was also clear from normalised read coverage across the Spn23F genome, generating a larger spike at the 0.3 Mb position where the CBL is located, relative to the unselected library (Supplementary Figure 7). Therefore, size selection notably increased enrichment when targeting the 23F CBL, in contrast to results from Viehweger *et al.* (*33*).

**Supplementary Table 4:** Read length statistics for CBL enrichment.

| Channel type | Read type | Experiment | Mean | Lower Quartile | Median | Upper Quartile |
| --- | --- | --- | --- | --- | --- | --- |
| Adaptive | Accepted | Unselected | 506 | 305 | 386 | 472 |
|  |  | Size selection | 703 | 271 | 362 | 475 |
|  | Rejected | Unselected | 485 | 414 | 475 | 546 |
|  |  | Size selection | 468 | 400 | 457 | 526 |
| Control | All | Unselected | 2026 | 560 | 1051 | 2281 |
|  |  | Size selection | 3092 | 874 | 1963 | 4045 |

**Supplementary Figure 6:**
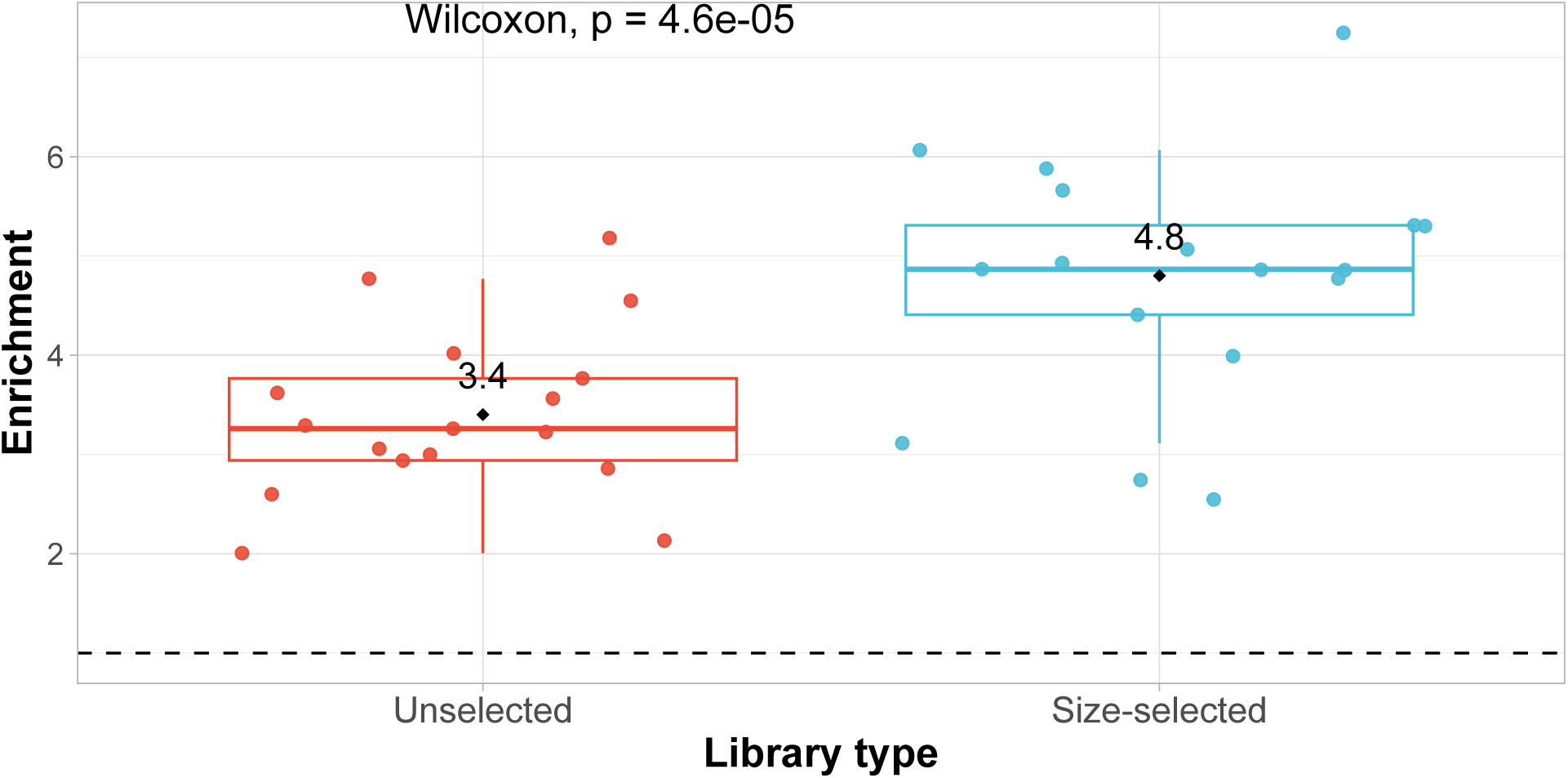
Enrichment comparison of pneumococcal CBL with and without size selection, which removed DNA fragments *<* 10 kb in length. Each data point represents the enrichment of a single CBL found within each library. In libraries containing one strain, only one CBL is present, in 50:50 mixtures, one or more CBL are present. Dashed line describes enrichment = 1 i.e. no enrichment has occurred. Black points highlight distributions means. Distributions were compared using a paired Wilcoxon test.

**Supplementary Figure 7:**
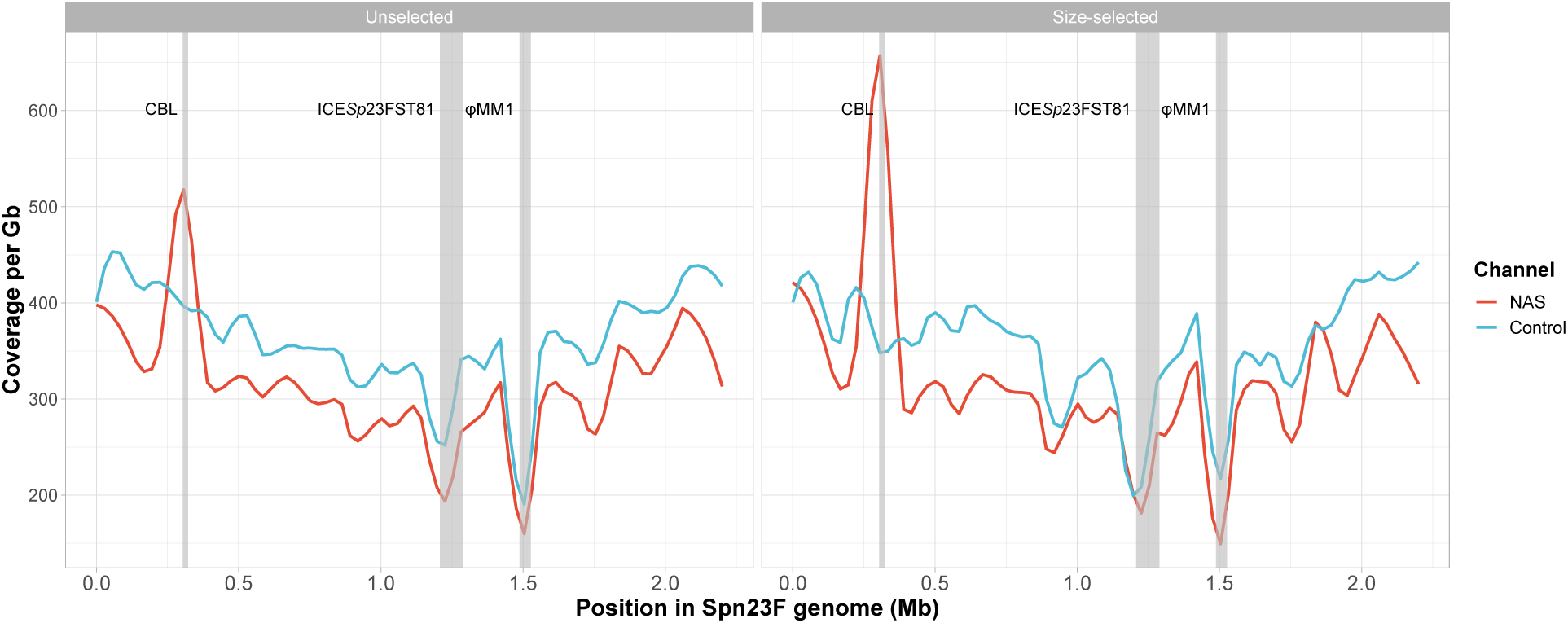
Normalised coverage across Spn23F genome for enrichment of 23F CBL for the Spn23F-only sample. Coverage was normalised by dividing the number of read bases aligned at each position in the Spn23F genome by the total number of bases generated for a sample by adaptive or control channels in gigabases (Gb). Columns describe library type; unselected or size-selected. Loci of interest are annotated by grey bars; CBL, as well as ICE*Sp*23FST81 and *ψ*MM1 prophage which are missing in Spn23F (*42*).

#### A.3 Implementing graph pseudoalignment in NAS to capture previously unobserved structural variants

As NAS effectiveness is impacted by how closely related reads are to the reference sequences (*31*, *33*), using references that are not closely related to the query can result in reference bias, whereby reads will either map in the wrong location, or not map at all (*44*, *73*), ultimately reducing enrichment. Reference bias can be overcome by including more sequences within the database, capturing a greater amount of diversity and increasing the chance of a match between the read and a reference sequence. However, alignment is still linearly restricted, meaning structural variation, such as reshuffling of genes seen in CBL (*21*), will not be captured and therefore previously unobserved structural variants cannot be enriched for (Supplementary Figure 8a).

**Supplementary Figure 8:**
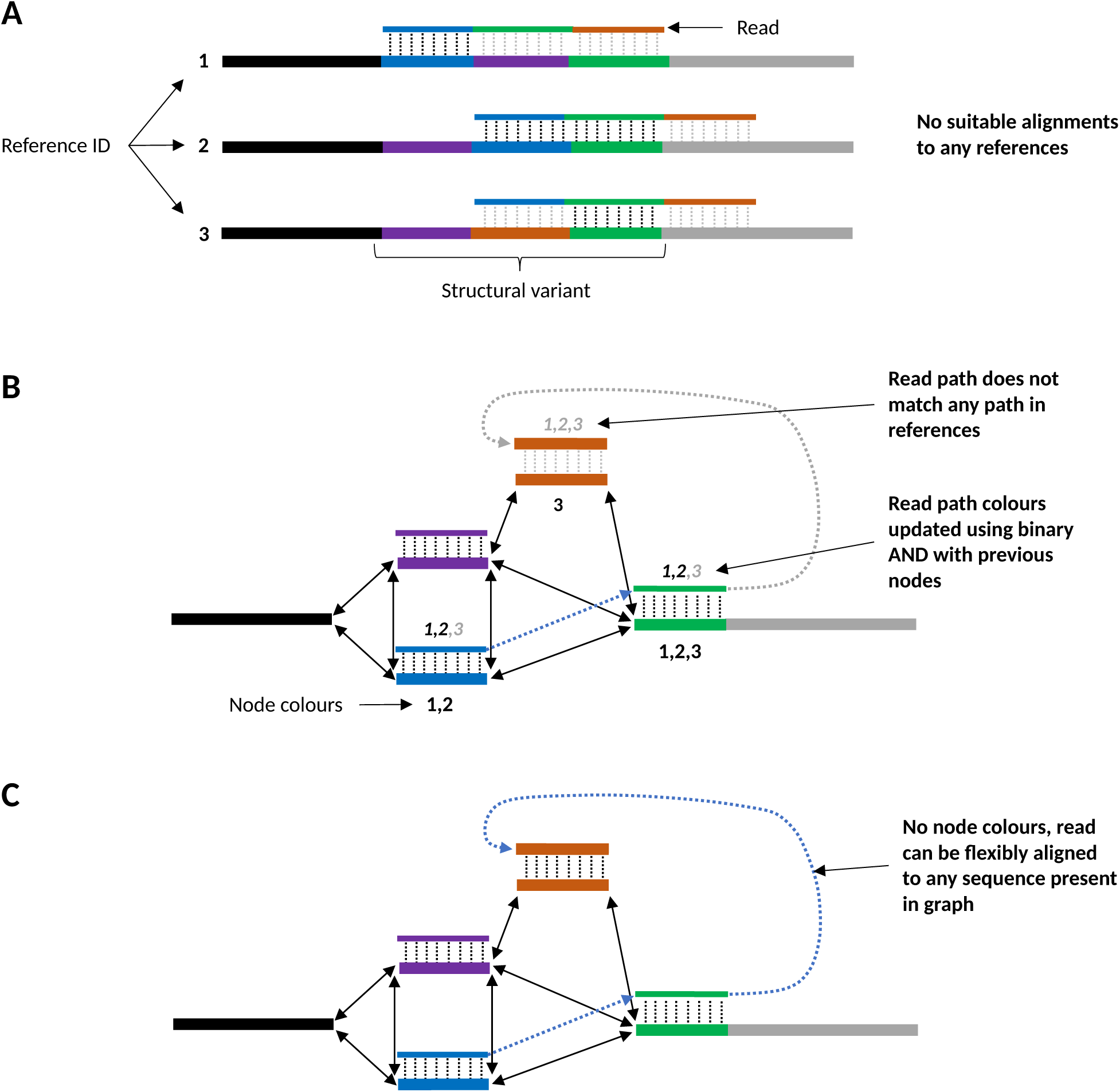
Example of alignment of a read containing a previously unobserved structural variant. (**a**) When aligning to a collection of linear reference sequences, the read does not align, despite the references containing the correct sequence blocks. (**b**) Using alignment with a coloured de Bruijn graph, the read is able to align more flexibly, however, as no colours contain all the same sequence blocks as the read, it cannot align fully. (**c**) Using alignment with an uncoloured de Bruijn graph, there are no restrictions on the path the read can take to align to the graph, therefore the read is able to align fully.

Alignment to pangenome graphs has been shown to improve recall of structural variants which are not present in reference sequences (*45*, *74*, *75*). Therefore, graph-based alignment can be integrated into NAS to aid in enrichment of new haplotypes. To achieve this, we developed a read-to-graph *k*-mer-matching method, referred to as graph pseudoalignment, using Bifrost (*51*) due to its scalability, fast *k*-mer matching functionality and intuitive C++ API. Pseudoalignment between reads and coloured DBGs has been used previously in transcriptomic and metagenomic studies to identify the likely source of reads (*48*, *49*). However, pseudoalignment to coloured DBGs is still restrictive, for example, if a read contains a set of sequences that are not found collectively in any single reference (Supplementary Figure 8b). Therefore, we implemented a method using pseudoalignment in uncoloured DBGs called GNASTy, which provides the greatest flexibility for identifying novel structural variants (Figure 8c).

As pseudoalignment operates in nucleotide-space, this method requires reads to be base-called from raw current signals generated by the sequencer before pseudoalignment. Therefore, we modified Readfish (*26*) to use graph pseudoalignment. Readfish takes raw current signals from a sequencer in ‘chunks’ (defined as a set period to allow sufficient time for reads to pass through a pore, default = 0.4 seconds), passes them to a base-caller, and then uses Minimap2 (*52*) to align the reads to linear references. Readfish then sends an accept or reject signal back to the sequencer for each read depending on the result of alignment. We replaced Minimap2 with the Bifrost-based DBG pseudoaligner in GNASTy, meaning Readfish can be run in either linear-alignment or graph pseudoalignment modes.

An overview of the graph pseudoalignment in GNASTy is shown in Figure 6. Before a sequencing run, an uncoloured DBG is built using Bifrost, with a user-specified value of *k* (default = 19 bp). At the start of a sequencing run, Readfish is initialised in graph-alignment mode. As each fragment is sequenced, Readfish passes the raw current to a base-caller, generating a nucleotide sequence (**Step 1**), which is then passed to the graph-pseudoaligner. The read must be above a length cutoff (default = 50 bp) to be aligned, otherwise a ‘proceed’ signal is sent to allow the read to be sequenced for another chunk. This step is designed to ensure that reads are sufficiently long to avoid false positive or negative mappings. If above the length cutoff, the read is split into its constituent k-mers (**Step 2**), with each k-mer being queried in the graph (**Step 3**). Once all k-mers in the read have been queried, a measure of similarity between the read and graph is calculated. This is either the proportion of matching k-mers, *P* (Equation 2), or a Jaccard index, *J* (Equation 3), based on the number of matching k-mers, *N*_match_, and mismatching k-mers, *N*_mismatch_, between the read and the hypothetical path in the graph. The Jaccard index assumes that for each mismatching k-mer in the read, there is also a respective mismatching k-mer in the path, meaning that for each mismatch the denominator increases by 2. The Jaccard index can be used to calculate a mash-like distance, *D*, as used in Mash and SKA (*76*, *77*) (Equation 4). Alternatively, taking 1 *− D* gives the mash-like index, *S* (Equation 5), an estimate of the sequence identity between the read and the graph, assuming a Poisson distribution of mutations within k-mers (*78*). Equation 4 can be simplified to Equation 5 to give *S* in terms of *P*.

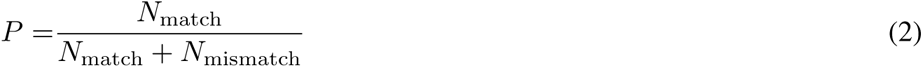

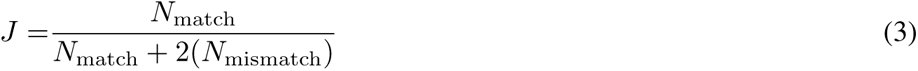

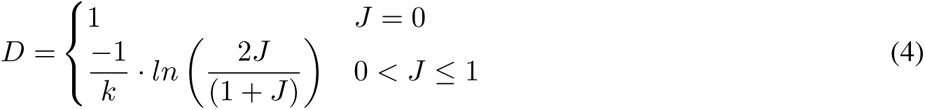

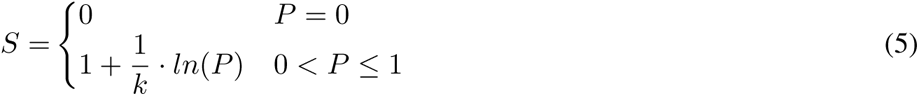

*S* is a more intuitive measure of similarity than *P*, which is difficult to interpret due to sensitivity to both *k* and identity between two sequences (*78*). We use *S* in simulations and empirical experiments below. If the similarity between the read and graph is greater than the identity cutoff, a ‘stop receiving’ signal is sent to allow the rest of the read to be sequenced, otherwise an ‘unblock’ signal is sent to reject the read (**Step 4**).

#### A.4 Evaluating mash-like index, *S*, as a similarity measure for graph pseudoalignment

We evaluated use of the mash-like index, *S* (Equation 5), which provides an estimate of the real identity between a read and a corresponding path in the DBG. Using simulations, we investigated the effect of varying an *S*-cutoff on accuracy of target and non-target assignment at varying *k* and mutation rate, *µ*. Reads were generated from a *k*-mer database generated from 100 sequences of 20 kb with 39.49% GC content, matching *S. pneumoniae* (*39*), and mimicking the database used for pneumococcal CBL enrichment (See Methods). 100 reads were generated from the reference database and randomly mutated at different rates before being decomposed into respective *k*-mers with varying size. The *k*-mers were then matched back the reference database to determine the effect of mutation rate, *µ*, and *k*-mer size on the sensitivity of pseudoalignment. Read length for all simulations was set to 200 bp, which approximately corresponds to a single chunk of sequence generated by Readfish (*26*).

Recall of target reads at varying cutoffs of *S* is shown in Supplementary Figure 9. At *µ ≤* 0.01, recall was 100% at all values of *k*, with the exception of high cutoffs of 98% or greater. At higher *µ*, *k* size had a large effect on recall of target reads, with large drops between cutoffs of 0 *< S ≤* 0.02, followed by ‘plateaus’ where no further reads were rejected, and finally by a sharp decline at cutoffs of 0.6 *≤ S ≤* 1.0. However, the *S* cutoff at which the second decline in target read assignment occurred was dependent on *k*, but not *µ*. These distributions can be explained by the proportion of matching *k*-mers in each scenario. Higher *µ* values cause a greater number of reads to have a no matching *k*-mers (*S* = 0), which is more likely at larger values of *k*, resulting in the initial sharp decline at 0 *< S ≤* 0.02. The plateau corresponds to the lower bound of *S* sensitivity at a given *µ* and *k*, when only a single *k*-mer matches between the read and the reference. At 0.6 *≤ S ≤* 1.0, the cutoff is sensitive to reads that have more than one matching *k*-mer. This sensitive range varies with *k* but not *µ*: for *k* = 19 the sensitive range begins at cutoff=0.75 for *µ* = 0.1 and *µ* = 0.16. In comparison, for *k* = 31 the sensitive range begins at a *S* cutoff of 0.85 for *µ* = 0.1 and *µ* = 0.16. Therefore, for read classification during pseudoalignment to be sensitive to sequence identity, choice of *S* must sit within the *µ* sensitive range for a given value of *k*.

**Supplementary Figure 9:**
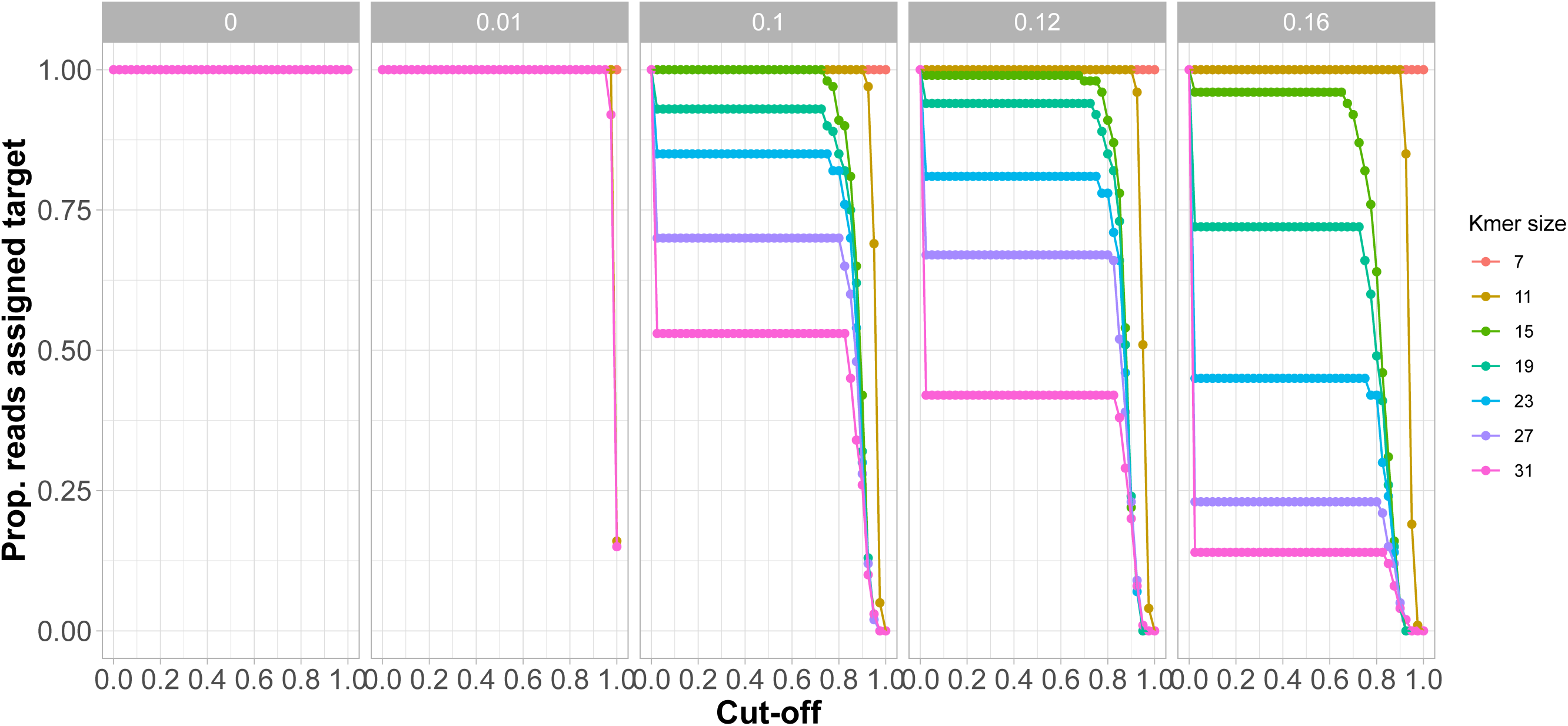
Effect of cutoff of mash-like similarity, *S*, and *µ* on recall of target reads. Plots show the proportion of reads identified as a target based on a set minimum cutoff of *S* between the read and reference database. Columns indicate the mutation rate in the reads per base, *µ*. Read length set at 200 bp.

We then investigated the concordance of *S* with sequence identity when varying *k* (Supplementary Figure 10) using LOESS regression, a form of linear regression with fits to data locally, resulting in a smooth regression line. Results highlighted that the agreement between *S* and pairwise identity is dependent on *k*. When *k <* 15, sequence identity is always overestimated due to random *k*-mer matching. For *k ≥* 15, when the read and reference match well (identity *≥* 0.9), *S* estimates pairwise identity correctly. At *≤* 0.9 identity, *S* underestimates the pairwise identity, with a larger underestimation at higher *k* due to increased *k*-mer mismatching.

**Supplementary Figure 10:**
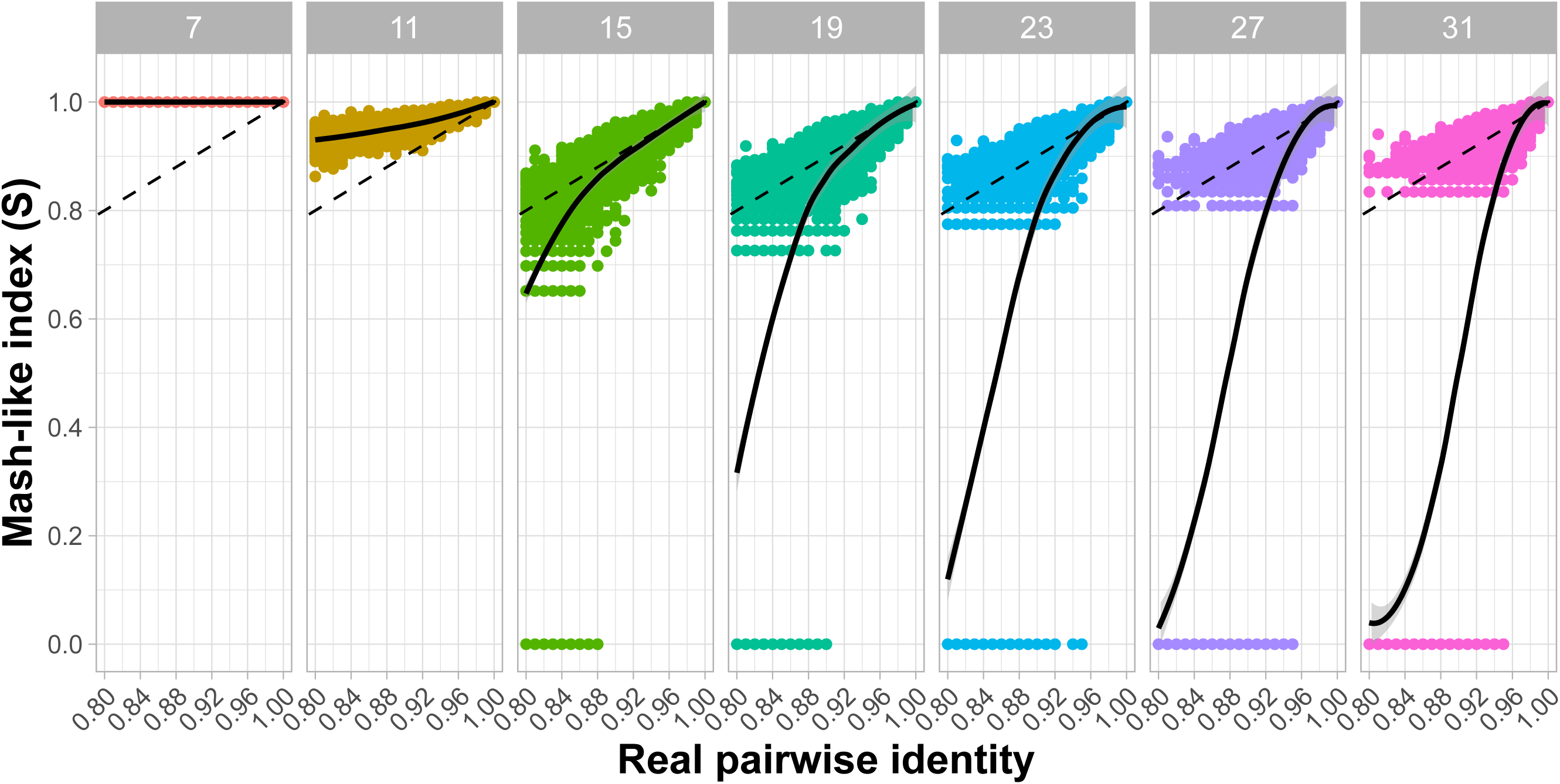
Concordance of mash-like distance, *S*, and identity between reads and reference sequence with varying *k* size. Each point describes a single read with a given value of *µ*. Columns describe *k* size. Dashed line represents identity line, y=x. Solid line represents LOESS regression between *S* and real identity, with grey shading for 95% confidence intervals.

We then calculated the root mean square error (RMSE) between *S* and the identity line, y=x, which is the ideal relationship between *S* and real pairwise identity, for real pairwise identity in range 0.9 *−* 1.0 at varying read lengths (Supplementary Figure 11). Results show that RMSE is reduced as read-length increases at all *k ≥* 15, due to the increased probability of *k*-mer matching in longer reads, and is lowest at *k* = 15. At *k <* 15, random *k*-mer matching means read-length has no impact on RMSE. Overall, agreement between *S* and the real pairwise identity of a read and reference sequence is dependent on *k*, read length and the underlying rate of mutation, *µ*. When *µ* is too large, *S* will tend to underestimate the real pairwise identity at *k ≥* 15, which can be remedied by increasing alignment length.

**Supplementary Figure 11:**
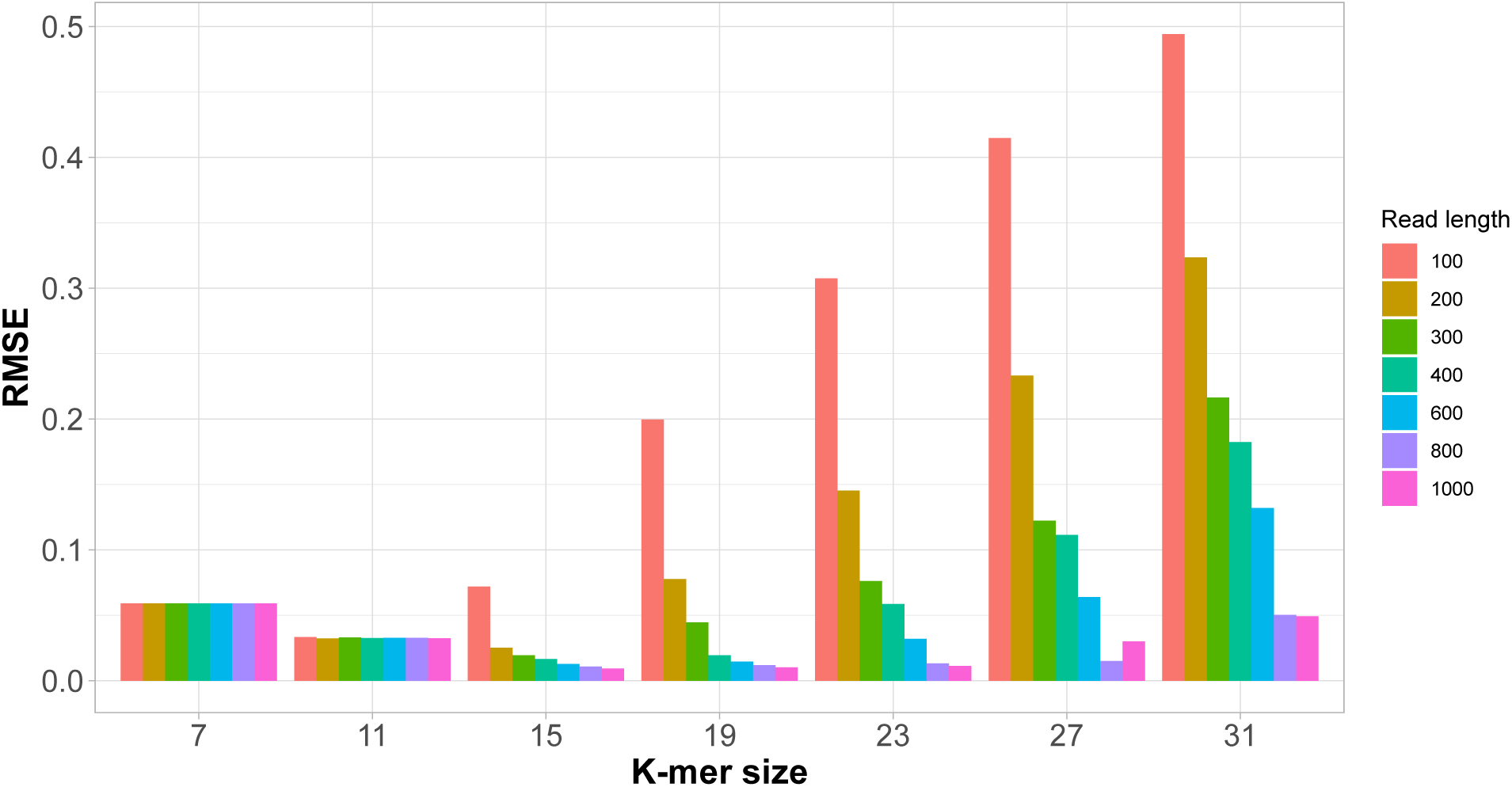
RMSE between identity line, y=x, and pairwise identity vs. *S* relationship for pairwise identity in range 0.9 *−* 1.0 for varying *k* size and read length.

We then considered the effect of the *S* cutoff and *k* on the specificity of pseudoalignment. 100 non-target reads were randomly generated from a different 2.2 Mbp sequence with the same GC content as before, and matched to the same reference k-mer database as used above. These reads represent non-target sequences, and so ideally should not match the original reference with high identity. At *k >* 15, almost no reads had *S >* 0, indicating that no *k*-mers matched between the non-target reads and the graph (Supplementary Figure 12). Contrastingly, for *k ≤* 15 there were a large proportion of non-target reads that matched with *S ≥* 0.5, indicating lower specificity of graph pseudoalignment with lower values of *k*.

**Supplementary Figure 12:**
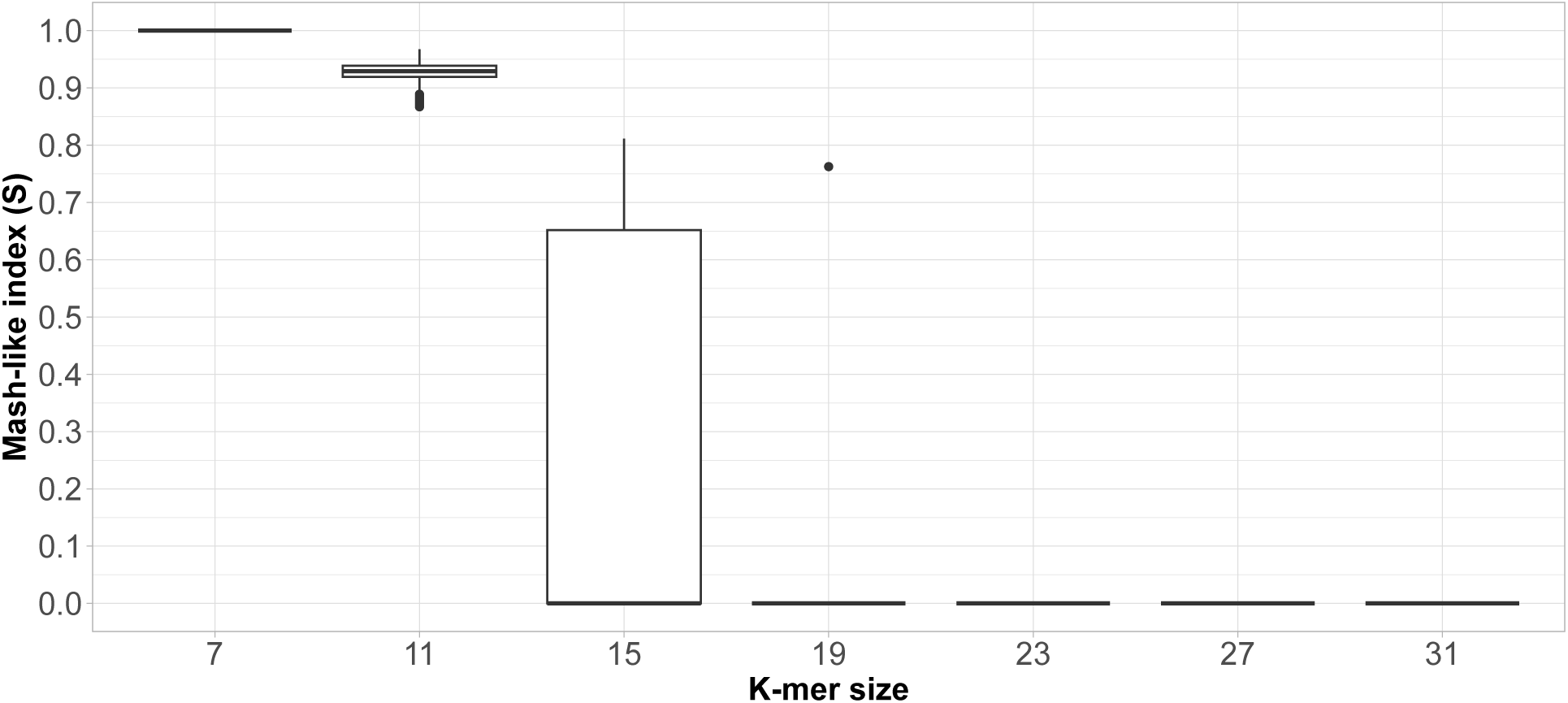
Effect of *k*-mer size on mash-like index, *S*, between non-target reads and reference. Box-plots show the proportion of random matching *k*-mers per read. Read length set at 200 bp.

To directly compare sensitivity and specificity of varying *S* cutoffs, we generated receiver operating characteristic (ROC) curves and conducted area-under-curve (AUC) analysis (Supplementary Figures 13 and 14). These visualisations compare the sensitivity and specificity of a parameter set as a cutoff is increased. An ideally performing parameter set should follow the top left corner of the ROC curve plot with an AUC of 1, indicating a high true positive rate and low false positive rate. A poorly performing parameter set will follow the line *y* = *x* and have an AUC of 0, which is equivalent to a random assignment of target or non-target. At *µ ≤* 0.01, all parameter sets with *k ≥* 7 had the ideal relationship between true positives and false positives. As *µ* increased, *k* = 15 had the best performance, with *k <* 15 having reduced specificity, and *k >* 15 having reduced sensitivity.

**Supplementary Figure 13:**
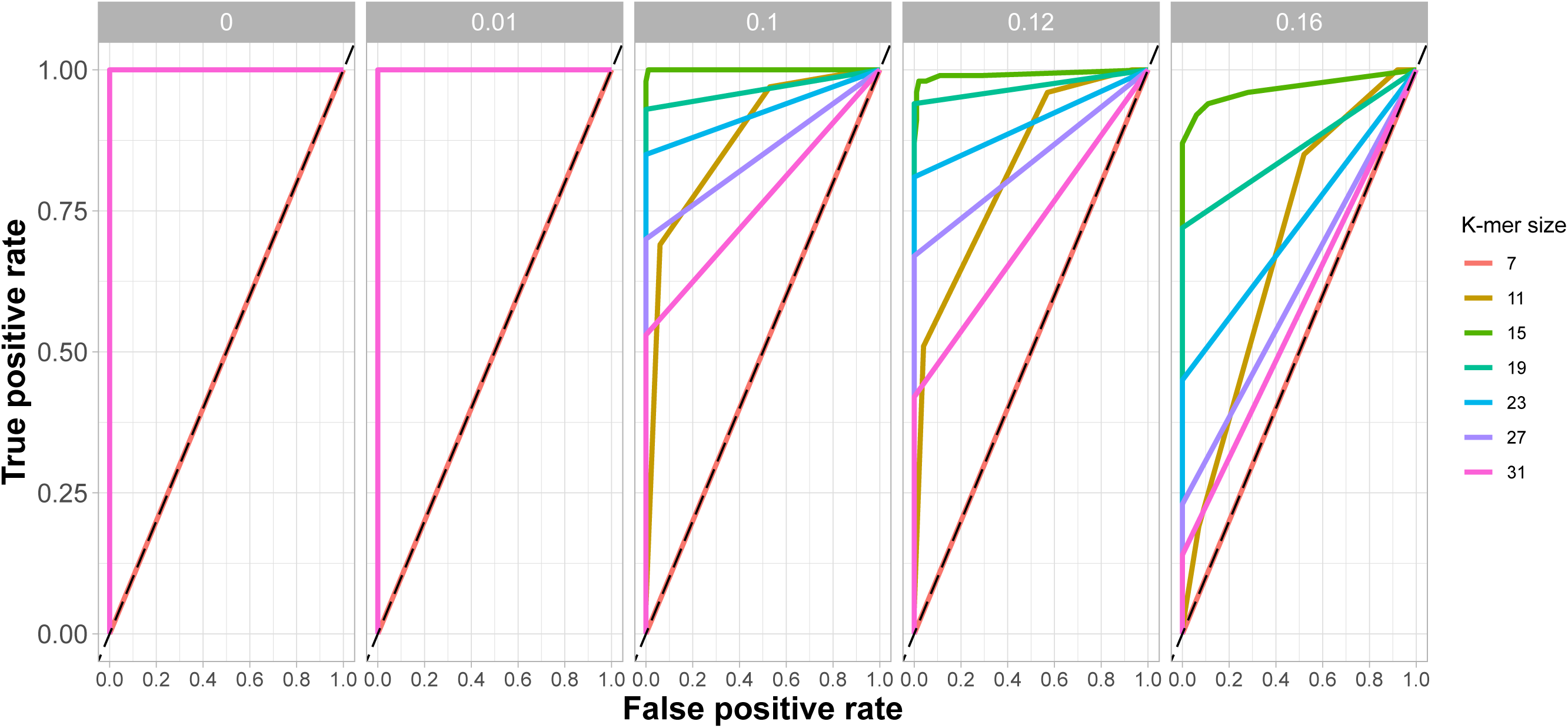
ROC curves for varying *S* cutoff and *µ*. Columns indicate the mutation rate in the reads per base, *µ*. Dashed lines represent line *y* = *x*. Read length set at 200 bp.

**Supplementary Figure 14:**
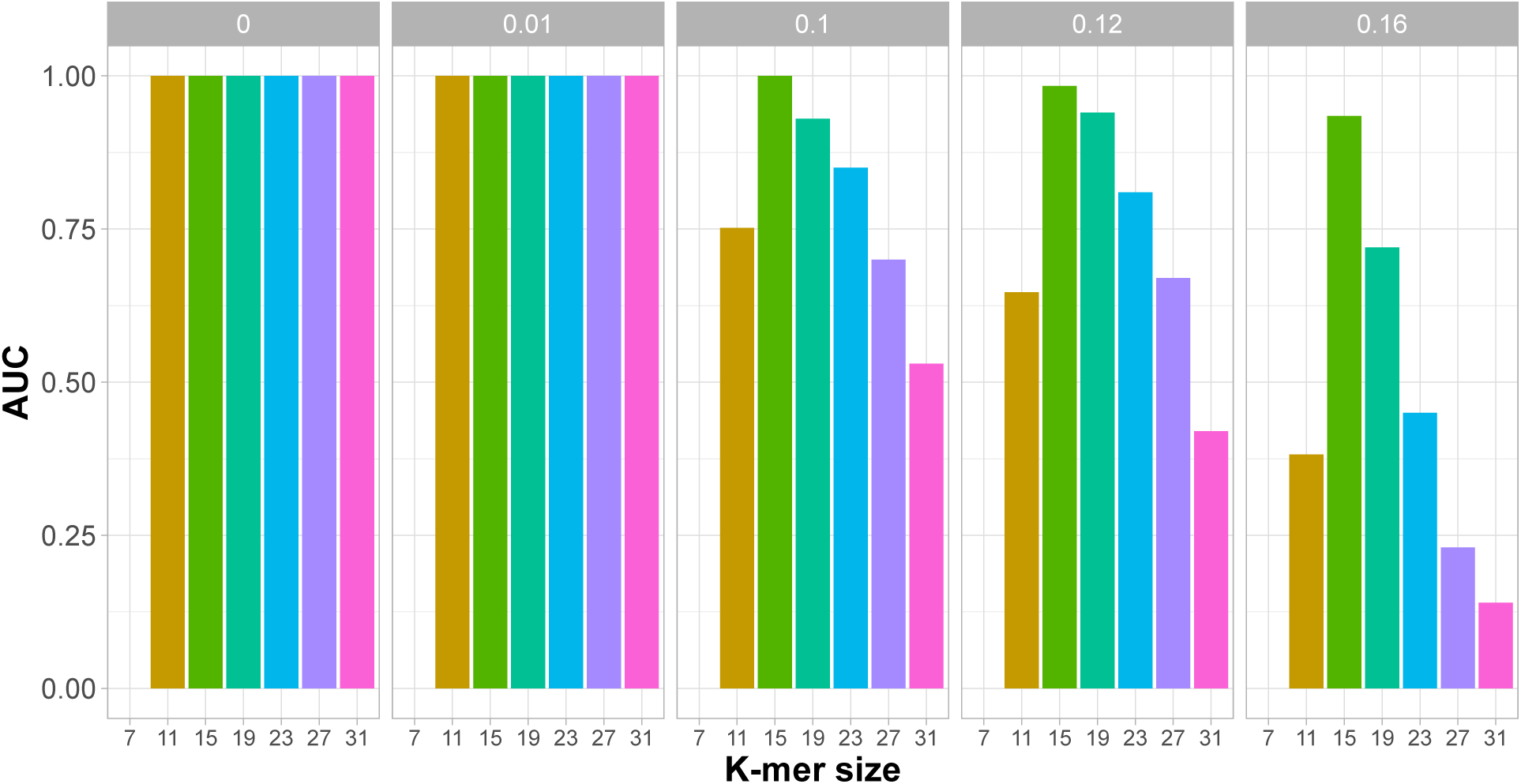
AUC from ROC curve for varying *k*-mer size and *µ* based on mash-index, *S*. Columns indicate the mutation rate in the reads per base, *µ*. Read length set at 200 bp.

Overall, concordance of *S* with *µ* was shown to be dependent on *k*, with *k ∼* 15 shown to have a good trade off of sensitivity and specificity even at *µ* = 0.16. However, measures of sensitivity and specificity were limited in these simulations, as non-target reads in real metagenomes will likely come from a greater diversity of genomes than the single 2.2 Mbp genome simulated here. Therefore, graph pseudoalignment parameters required further simulation-based optimisation to determine the best combination to use empirically.

#### A.5 Evaluating Graph Pseudoalignment using simulated sequencing runs

The simulations in the Section A.4 are useful for understanding how pseuodalignment parameters impact sensitivity and specificity, and guided down-selection of *k* to *∼* 15. However, there was no clear choice of *k* and *S* cutoff for optimal performance. Therefore, before testing graph pseudoalignment empirically, we ran more realistic simulations where the process of sequencing and read rejection were simulated, whilst pseudoalignment parameters were varied for down-selection.

For simulations, we used a ‘recording’ of the Spn23F whole genome enrichment sequencing experiment using a size-selected library from Section 2.1, containing a 50-50 mixture of Spn23F and *E. coli*. This recording generated simulated target and non-target reads, which were then aligned using graph pseudoalignment or Minimap2 to the CBL database used in Section 2.2, enriching for the 23F CBL. These simulations result in rejected reads being ‘chopped’ into smaller reads, rather than being rejected, as would occur in a normal sequencing run (*79*). Therefore, in order to compare the effect of graph pseudoalignment parameters, we analysed the read lengths of four read categories based on the rejection decision and post-sequencing alignment to the 23F CBL: true positives (TPs), reads that were correctly accepted; false positives (FPs), reads that were incorrectly accepted; true negatives (TNs), reads that were correctly rejected; and false negatives (FNs), reads that were incorrectly rejected. Ideally, TPs should be long relative to the other categories, as these reads have been accepted and therefore sequenced fully. FPs and TNs should be as short as possible; FPs should ideally be reads too short to have been rejected, whilst short TNs indicate fast rejection time. FNs should not be present, as they result from acceptance parameters that are too conservative.

**Supplementary Figure 15:**
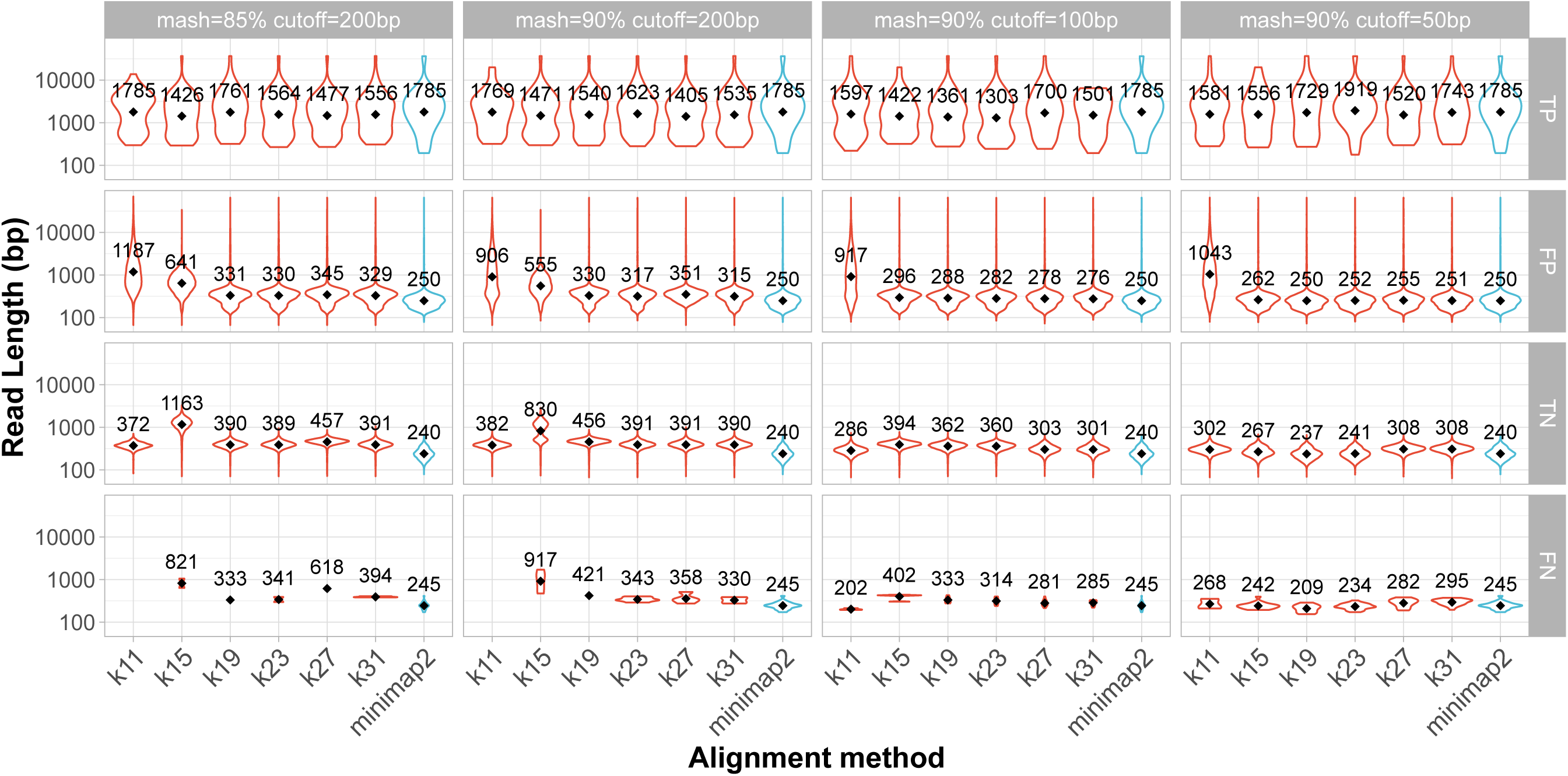
Simulation read length comparison of graph pseudoalignment parameters and Minimap2. X-axis describes either the *k* size used in graph pseudoalignment or Minimap2 alignment. Columns describe pseudoalignment parameters (mash: minimum *S*, cutoff: minimum read length). Rows describe categories of reads based on accept/reject decision and post-alignment to 23F CBL; TPs, reads that were correctly accepted, FPs, reads that were incorrectly accepted, TNs, reads that were correctly rejected, and FNs, reads that were incorrectly rejected.

We initially tested *S* = 85%, as simulations in the previous section highlighted this was within the sensitive range of all *k* values (Supplementary Figure 9), as well as a minimum length cutoff of 200 bp (approximate size of one chunk from Payne *et al.* (*26*)), which sets the minimum length of a read that can be aligned. An appropriate minimum length cutoff ensures that reads are not rejected prematurely if the initial sequence generated in the first chunk aligns poorly. TP read lengths were similar across all *k* values, and comparable to Minimap2 (Supplementary Figure 15, left). However, FP and TN reads were longer on average than Minimap2 for all values of *k*, indicating either slower rejection speed or lower specificity. Only *k* = 11 resulted in no FN reads due to low specificity but high sensitivity, resulting in no reads being rejected incorrectly.

As longer FP reads indicated graph pseudoalignment may have lower specificity than Minimap2, we increased *S* to 90% (Supplementary Figure 15, center-left). This change reduced the average size of FP reads without affecting TPs for all values of *k*, however, TNs were still longer than Minimap2. We therefore decreased the minimum length cutoff to 100 bp and 50 bp with the aim of increasing rejection speed (Supplementary Figure 15, center-right and right respectively). Reducing this parameter to 50 bp had no impact on TP read lengths, but reduced the FP and TN lengths to similar ranges observed with Minimap2.

To determine the best pseudoalignment parameter set, we compared the proportion of bases assigned to TP and TN categories, rather than read lengths. We ignored FNs and FPs, as these are equivalent to 1*−*TP and 1*−*TN respectively. Graph pseudoalignment performed similarly to, or outperformed, Minimap2 at *k >* 15 (Supplementary Figure 16). Proportions of TP bases were always higher for graph pseudoalignment than Minimap2, but were reduced as parameters became more stringent (increasing *S*, decreasing length cutoff). Proportions of TN bases were also the same or greater for *k >* 15 for all parameters, and did not change substantially at *k ≥* 19.

Additionally, we compared parameters based on their *F*_1_ score based on base proportions:

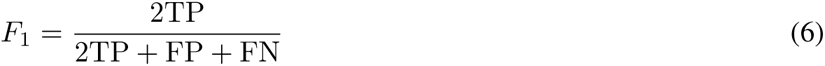

which describes the trade off between sensitivity and specificity, with a value of 1 indicating recall of all TPs with no FPs. Based on *F*_1_, the most stringent parameter set (*k* = 31, *S* = 90%, minimum length cutoff 50 bp) performed the best, also outperforming Minimap2 (Supplementary Figure 17).

**Supplementary Figure 16:**
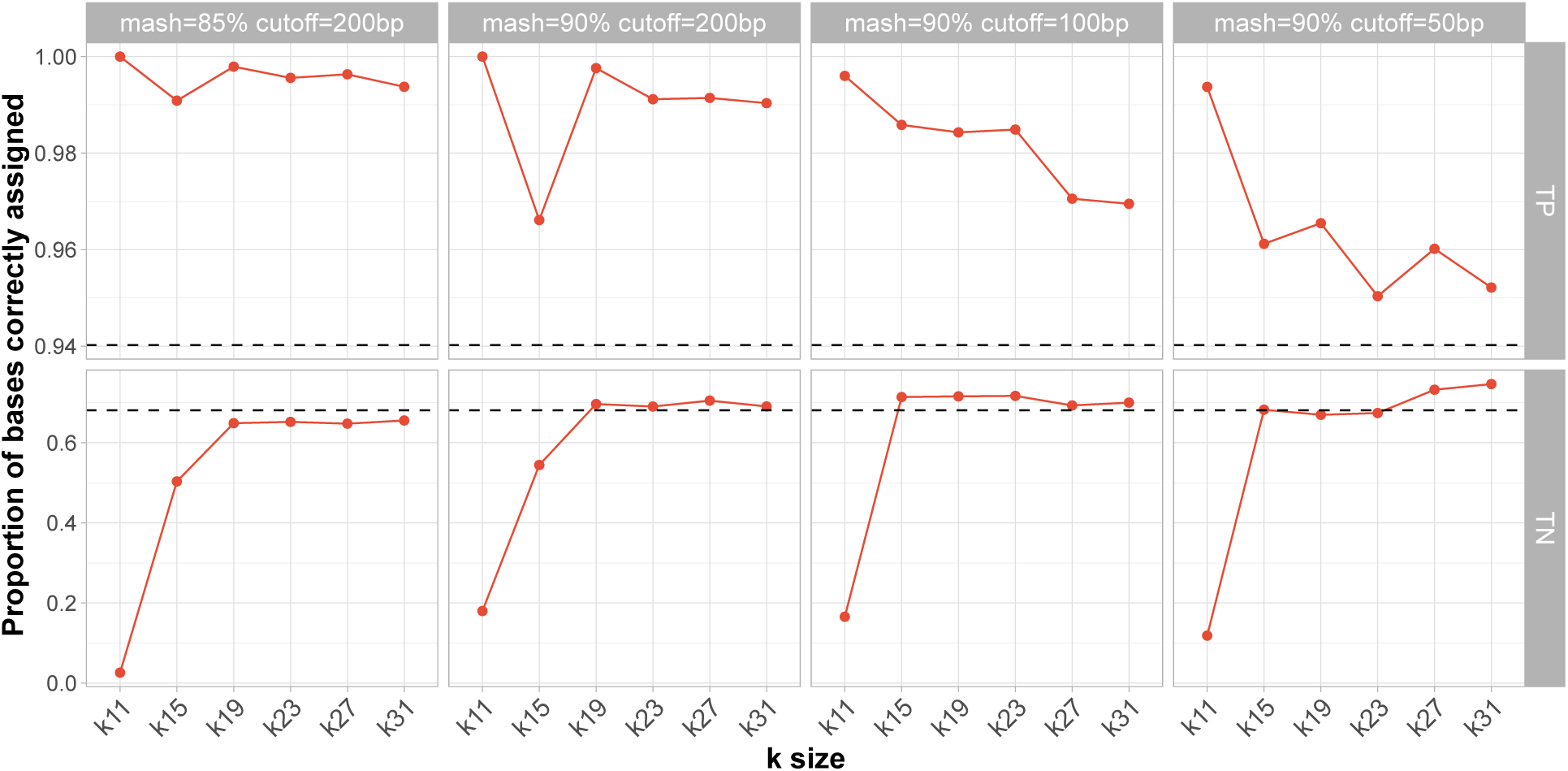
Comparison of graph pseudoalignment parameters and Minimap2 based on proportions of bases correctly assigned to positive and negative categories. Minimap2 results are indicated by the black dashed line on each facet. Columns describe pseudoalignment parameters (mash: minimum *S*, cutoff: minimum read length). Rows describe categories of reads based on accept/reject decision and post-alignment to 23F CBL: TPs, reads that were correctly accepted; TNs, reads that were correctly rejected.

**Supplementary Figure 17:**
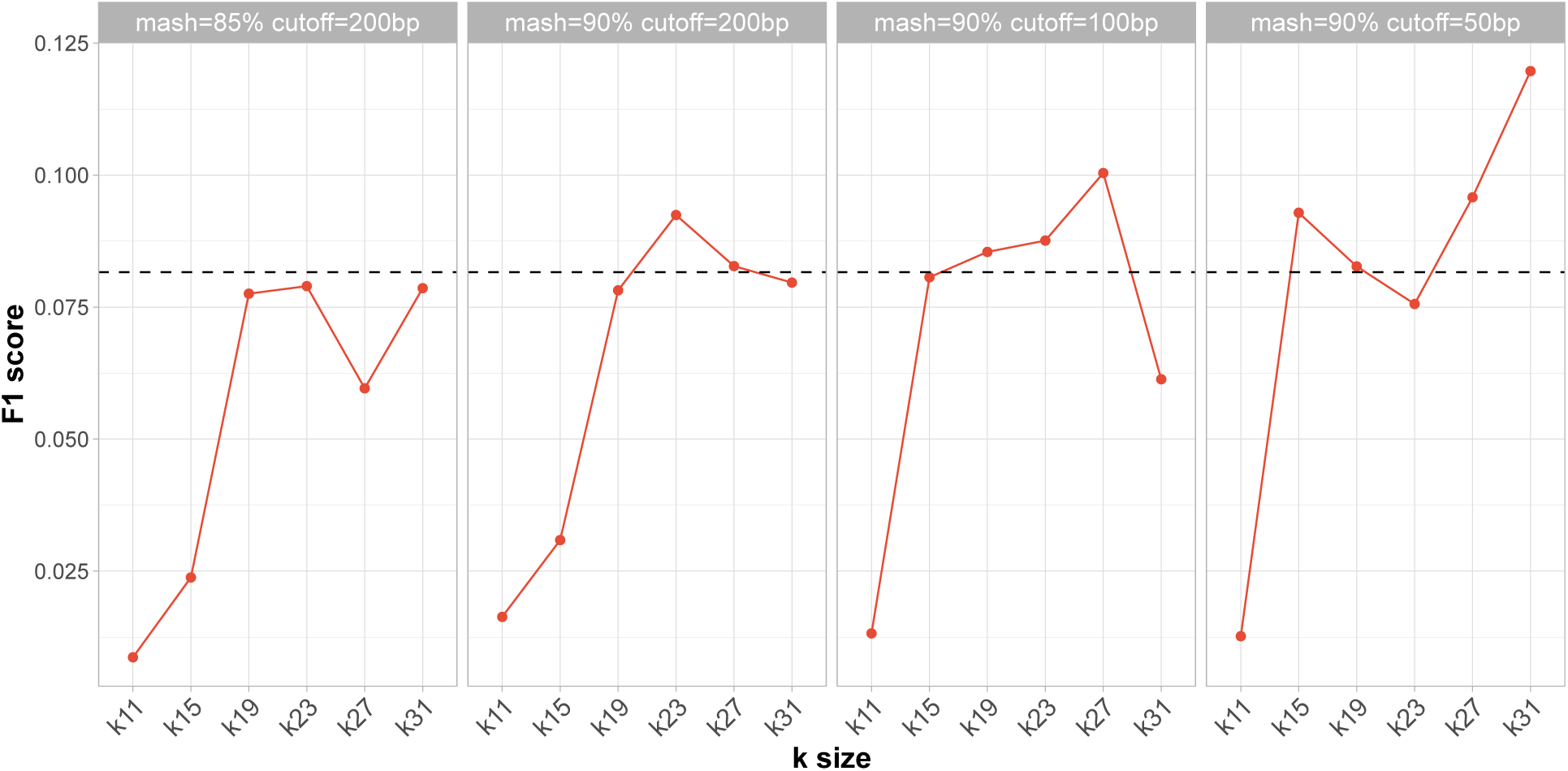
Comparison of *F*_1_ scores of graph pseudoalignment parameters and Minimap2 based on base proportions. X-axis describes either the *k* size used in graph pseudoalignment. Minimap2 results are indicated by the black dashed line on each facet. Columns describe pseudoalignment parameters (mash: minimum *S*, cutoff: minimum read length).

We hypothesised that graph pseudoalignment will outperform Minimap2 when enriching for an unobserved sequence due to more flexible alignment (Supplementary Figure 8). To compare the performance of graph pseudoalignment and linear alignment with a reference database where the target sequence is missing, we generated a partial reference database containing divergent sequences from 23F CBL. Using cluster assignments from Mavroidi *et al.* (*53*), we included sequences from CBL outside cluster two only, which contains the 23F CBL. As shown in Supplementary Figure 18, the presence of homologous blocks between 23F and CBL in different clusters varies between clusters. For example, the 23A CBL, in the same cluster as 23F, has 12 blocks in common, covering *∼* 17.5 kb, whilst 25F, belonging to cluster 3, has only two blocks in common, covering *∼* 3.5 kb. By combining CBL sequences from different clusters apart from cluster two, we were able to test the ability of graph pseudoalignment to enrich for haplotypes missing from the reference database.

**Supplementary Figure 18:**
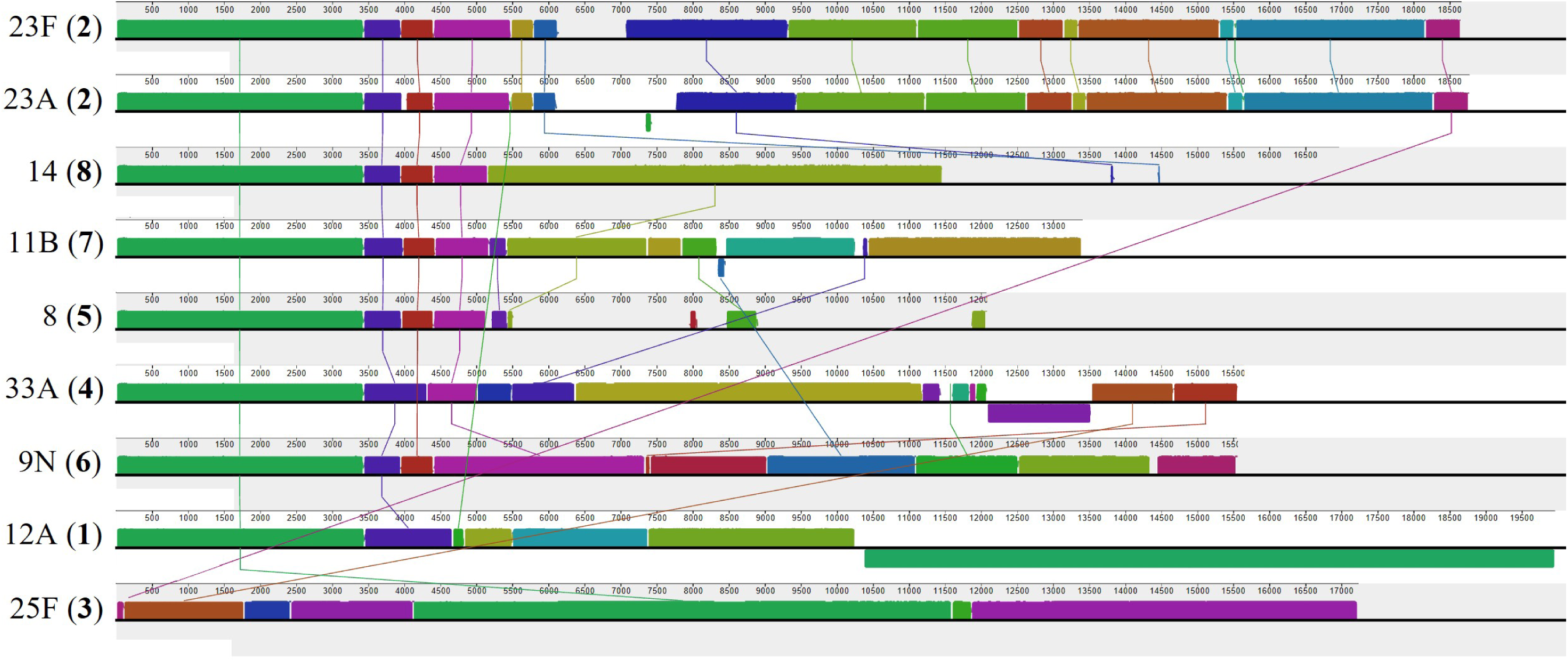
Structural alignment of representative pneumococcal CBL. Labels describe the serotype, followed by the cluster assignment from Mavroidi *et al.* (*53*) in brackets. Alignments were generated using ProgressiveMauve and visualised by MauveViewer (*80*). Same coloured blocks were identified as homologous by ProgressiveMauve between CBL and are connected by coloured lines. CBL coordinates are given for each CBL sequence.

We re-ran simulations using the same run recording used above, aligning to the partial CBL reference database using Minimap2 and graph pseudoalignment, aiming to enrich for the 23F CBL. We varied *k*, keeping *S* and read-length cutoff at the best performing parameters from previous simulations (*S* = 90%, read length cutoff 50 bp). Results showed that with a partial database, *k* = 19 performed best of all parameters, with a combination of high TP and TN, leading to the only *F*_1_ score above that of Minimap2 (Supplementary Figures 19 and 20). These results indicate that reduced stringency should be used when a target is not present in the reference database, as larger values of *k* lead to incorrect rejections due to *k*-mer mismatching.

Overall, we have shown that graph pseudoalignment can theoretically increase the sensitivity of target read identification. The best performing parameters from these simulations did not agree with results from Section A.4, with *k* = 19 and *k* = 31 shown to perform best, rather than *k* = 15. Therefore, alignment identity parameters, *k* and *S*, can have a large effect on NAS accuracy, which cannot be entirely modelled using simple simulations.

**Supplementary Figure 19:**
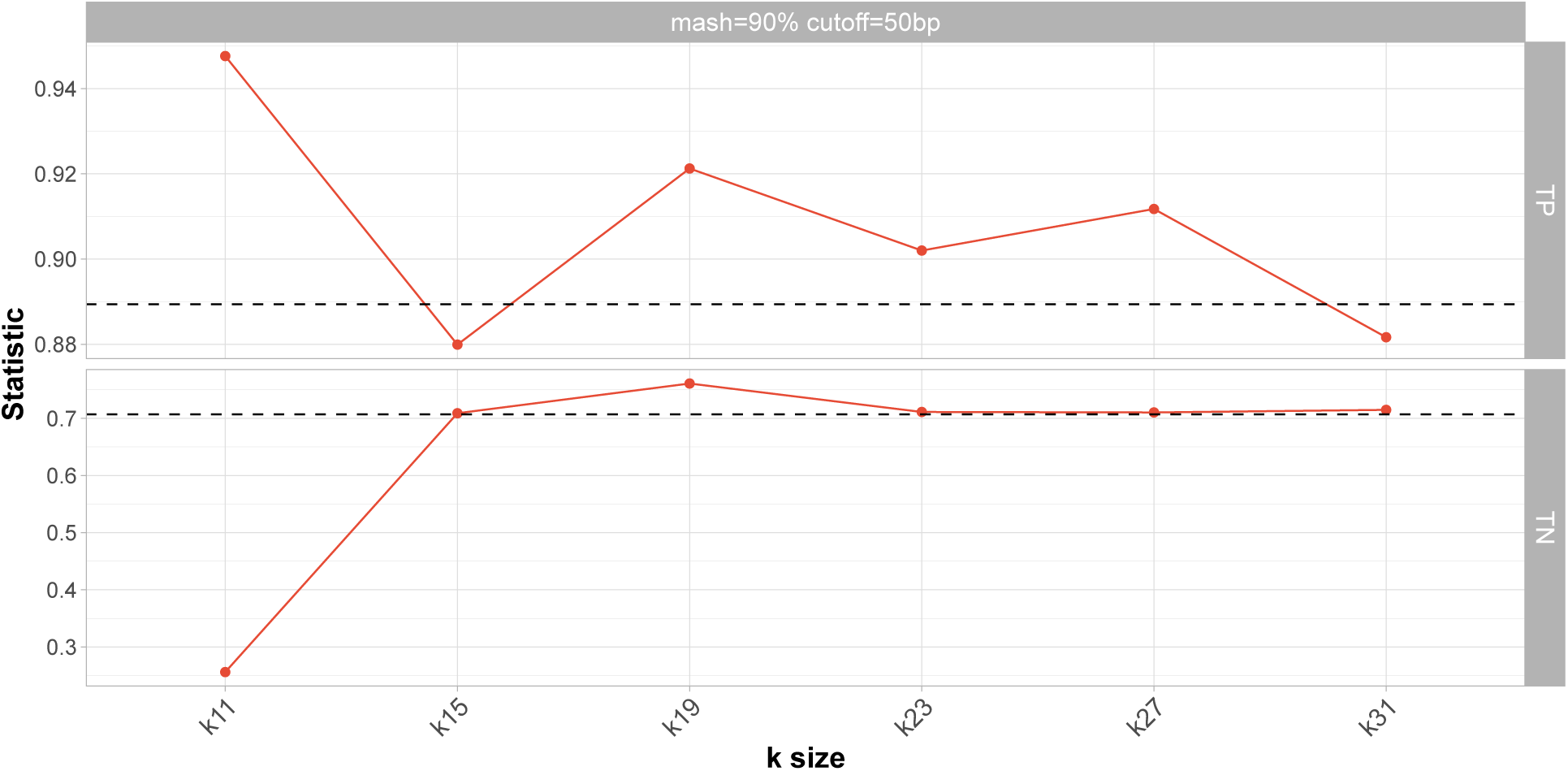
Comparison of graph pseudoalignment parameters and Minimap2 based on base proportions. X-axis describes either the *k* size used in graph pseudoalignment. Minimap2 results are indicated by the black dashed line on each facet. Facet heading describes pseudoalignment parameters. Rows describe categories of reads based on accept/reject decision and post-alignment to 23F CBL; TPs, reads that were correctly accepted, TNs, reads that were correctly rejected.

**Supplementary Figure 20:**
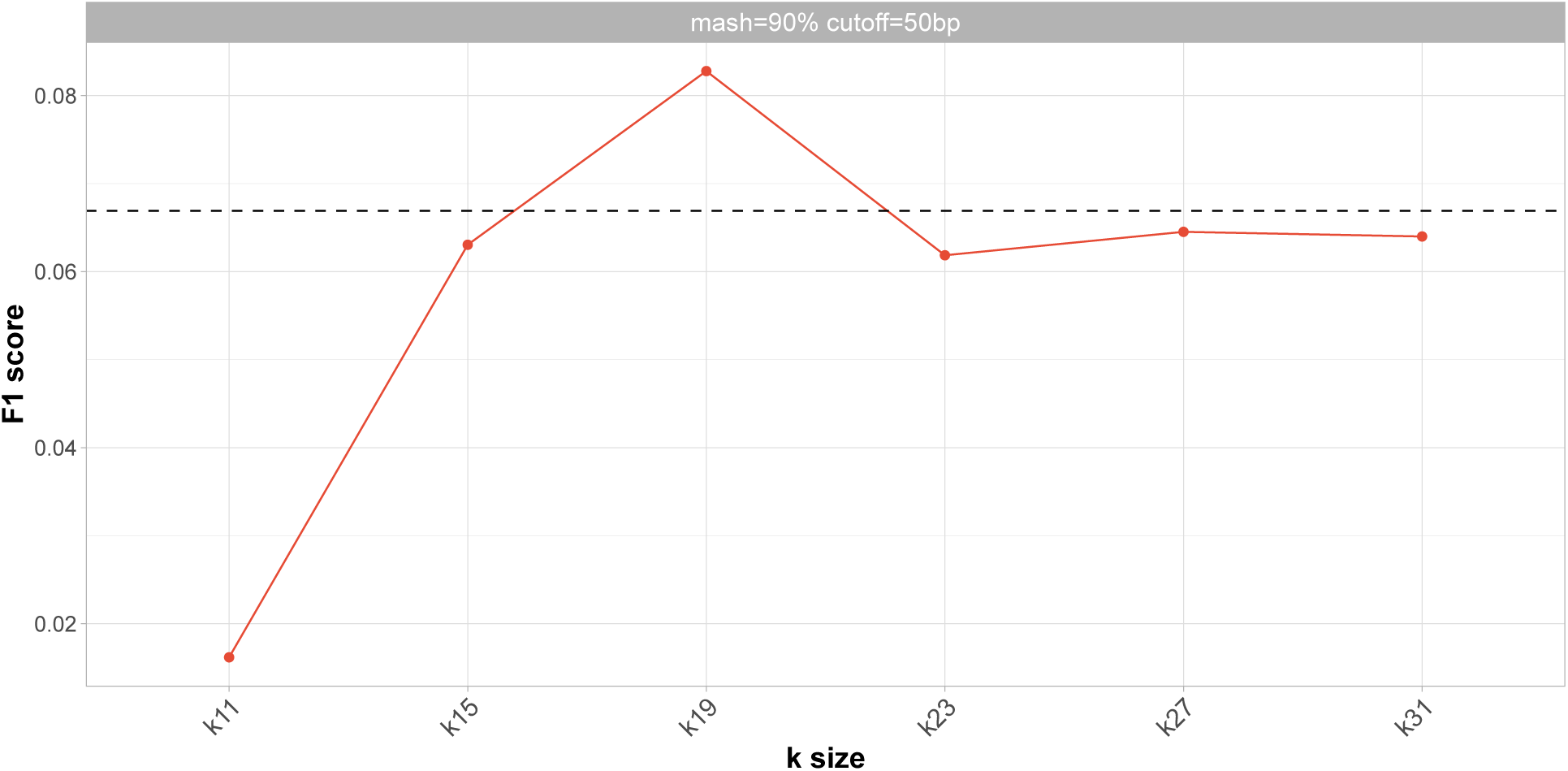
Comparison of *F*_1_ scores of graph pseudoalignment parameters and Minimap2 using a partial CBL database. X-axis describes either the *k* size used in graph pseudoalignment. Minimap2 results are indicated by the black dashed line on each facet. Facet heading describes pseudoalignment parameters.

#### A.6 Graph pseudoalignment performs similarly to linear alignment when enriching for an observed locus

To determine whether graph pseudoalignment in GNASTy could perform equally well using a full CBL database, we enriched for the 23F CBL using a reference database containing all 106 CBL sequences, including 23F. The experimental setup was the same as described in Figure 1a and b, with GNASTy using parameters *k* = 19*, S* = 75%.

GNASTy performed similarly to Minimap2, with either method performing better at specific target concentrations and non-target species (Supplementary Figure 21). Both methods significantly increased the absolute yield of 23F bases compared to control channels by 2.8- and 3.4-fold on average for Minimap2 and GNASTy respectively (Supplementary Figure 22). Normalised coverage was similar between the two methods (Supplementary Figure 23). Moreover, coverage was more even across the 23F CBL using the full database relative to the partial database (Supplementary Figure 29), particularly when comparing the samples containing CBL proportion at 4 *×* 10*^−^*^3^. This finding suggests that the drop in coverage in the central region of the 23F CBL was caused by absence of similar sequences in the reference database. Therefore, fragments originating from this central region were incorrectly rejected, observed when using Minimap2 or graph pseudoalignment, although graph pseudoalignment maintained greater coverage across the remainder of the CBL.

**Supplementary Figure 21:**
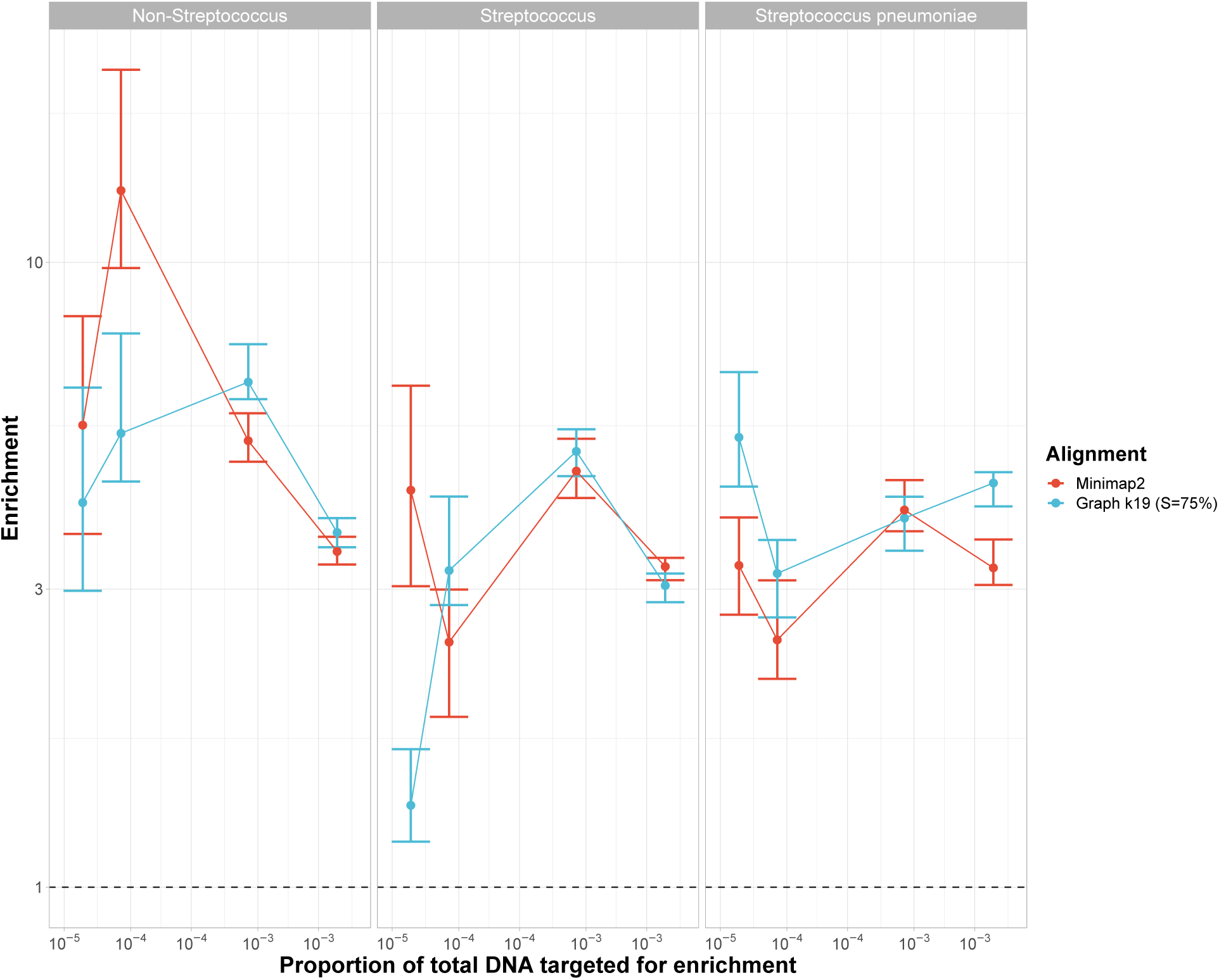
Enrichment comparison of 23F CBL at different concentrations of target between Minimap2 and graph pseudoalignment in GNASTy when aligning to a full CBL reference database using V14 chemistry. Bar ranges are inter-quartile range of enrichment from 100 bootstrap samples of reads. Data points connected by lines are observed enrichment values for each library, with solid lines connecting the same genome diluted at different concentrations. Columns describe the type of non-target species (Non-*Streptococcus* = *E. coli*, *Streptococcus* = *S. mitis* and *S. pneumoniae* = *S. pneumoniae* R6). Dashed line describes enrichment = 1 i.e. no enrichment has occurred.

**Supplementary Figure 22:**
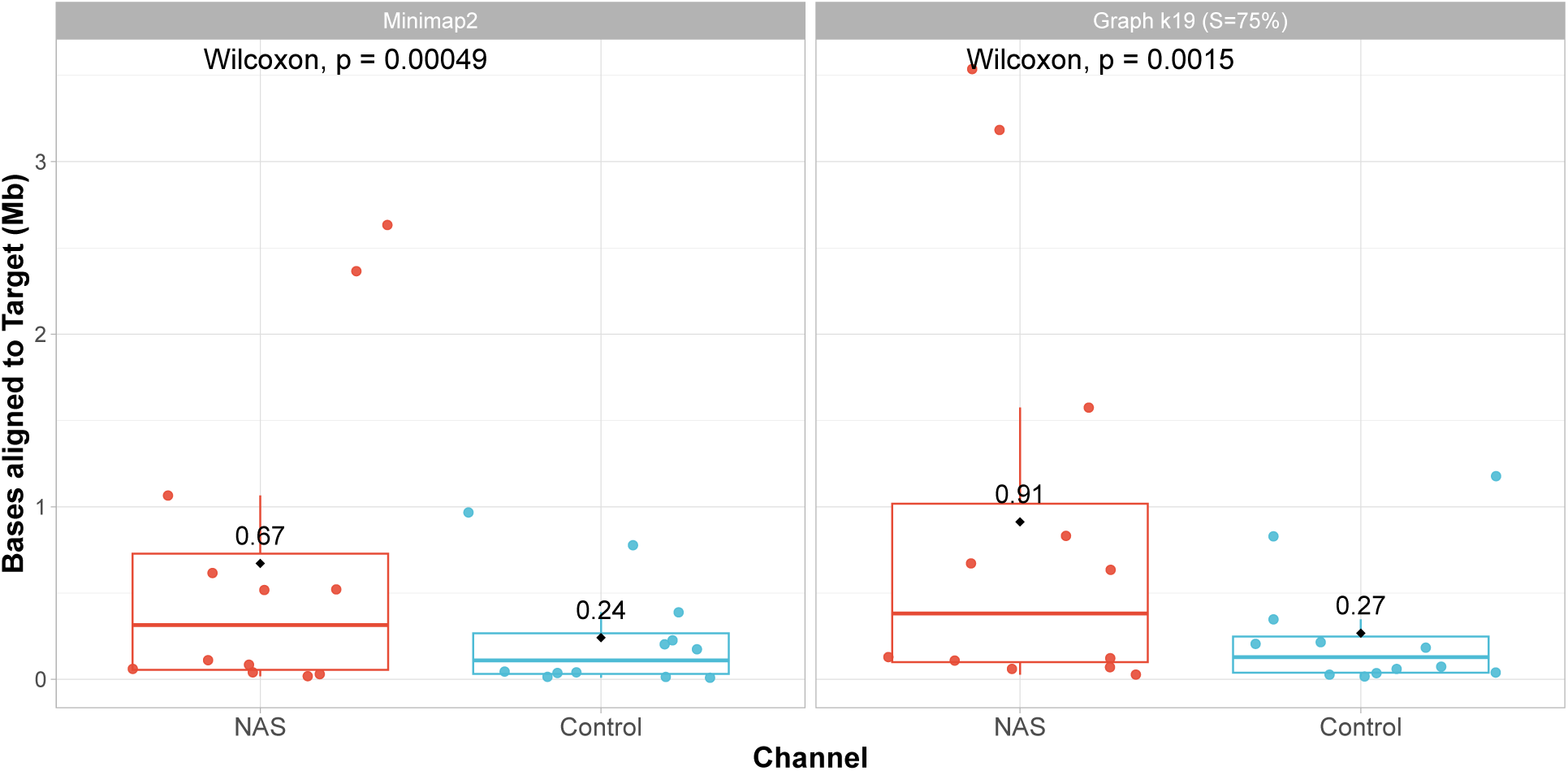
Absolute yield (in megabases) of bases aligning to the 23F CBL when aligning to a full CBL database. Each data point represents the enrichment of the 23F CBL found within each library. Distributions from control and NAS channels were compared using a paired Wilcoxon test.

**Supplementary Figure 23:**
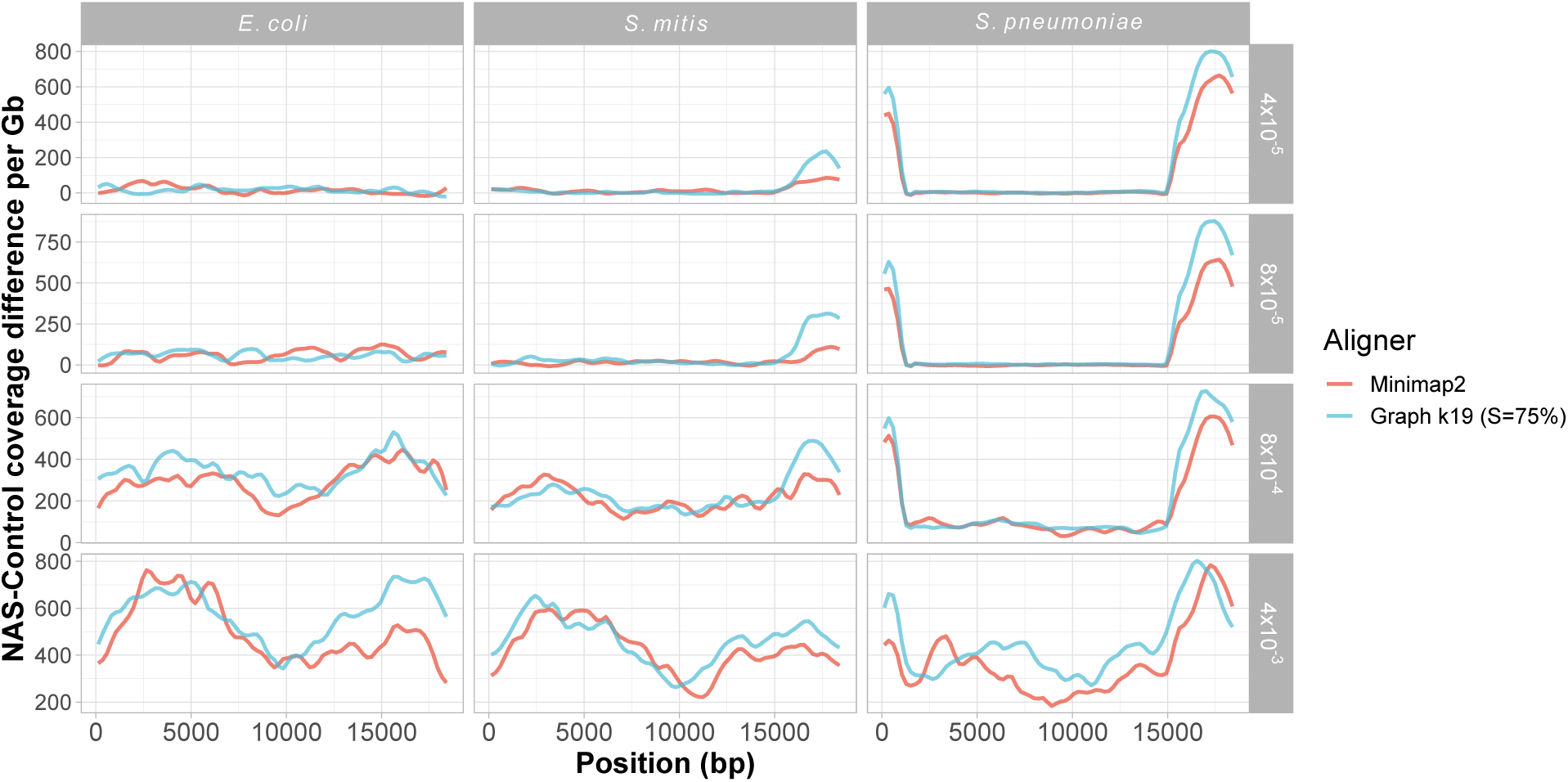
Normalised coverage difference between NAS and control channels across 23F CBL using a full CBL reference database. NAS-control coverage difference per gigabase (Gb) calculated by normalising the read coverage for each locus by the amount of data generated (in Gb) for each respective sample and channel, and then negating the normalised coverage for control channels from NAS channels for each locus. Rows describe the proportion of target DNA present in the sample.

Comparison of assembly quality highlighted similar performance between Minimap2 and graph pseudoalignment in GNASTy (Supplementary Figure 24). Read coverage was higher for adaptive channels than control channels, although assembly quality was similar between control and NAS channels for both tools. For Minimap2, assembly quality was identical between NAS and control channels. For GNASTy, a gap present in the centre of the assembly from control channels was covered in the NAS channel assembly, as observed when using the partial CBL database for alignment (Supplementary Figure 31). However, the NAS channel assembly was missing a section at the 18 kb end of the CBL, which was captured by the control assembly, despite read coverage being similar or higher from adaptive channels. This effect was also observed in Supplementary Figure 31 and discussed in Section 2.5.

**Supplementary Figure 24:**
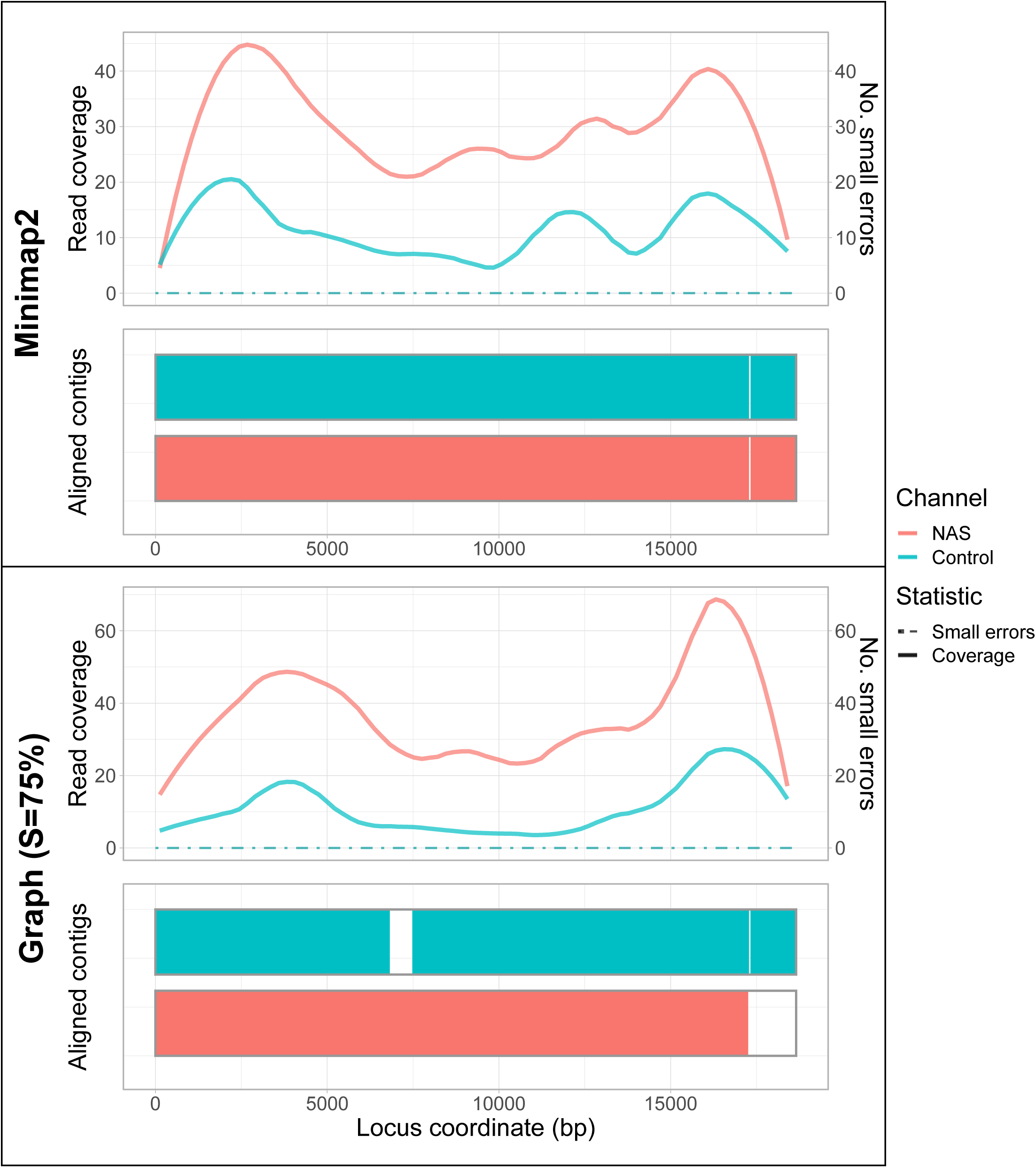
Spn23F CBL assembly comparison across alignment methods using a full CBL database during NAS. Each panel describes a Spn23F assembly generated from 0.1 Spn23F dilutions with *S. mitis*. For each panel, the top plot shows the read coverage (solid), defined as the absolute number of bases aligning to a locus, and number of small errors (*≤* 50 bp, dashed), whilst the bottom plot shows aligned contigs (colours) and large errors (*>* 50 bp) in each assembly.

### B Supplementary Figures

**Supplementary Figure 25:**
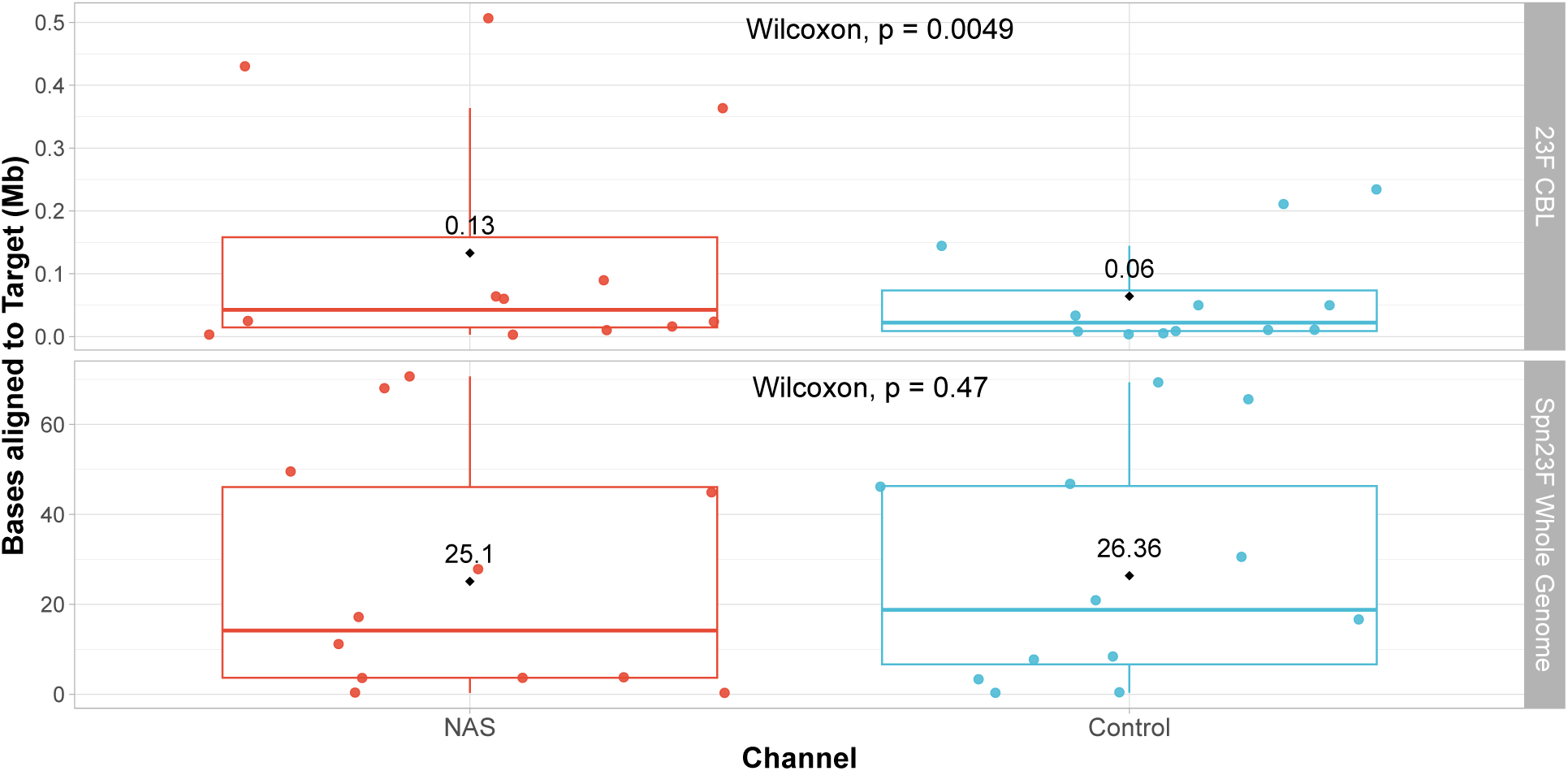
Absolute yield in megabases (Mb) of bases aligning to the whole genome of Spn23F **(top)** and 23F CBL **(bottom)**. Each data point represents the enrichment of the Spn23F whole genome or 23F CBL found within each library. Distributions from control and NAS channels were compared using a paired Wilcoxon test.

**Supplementary Figure 26:**
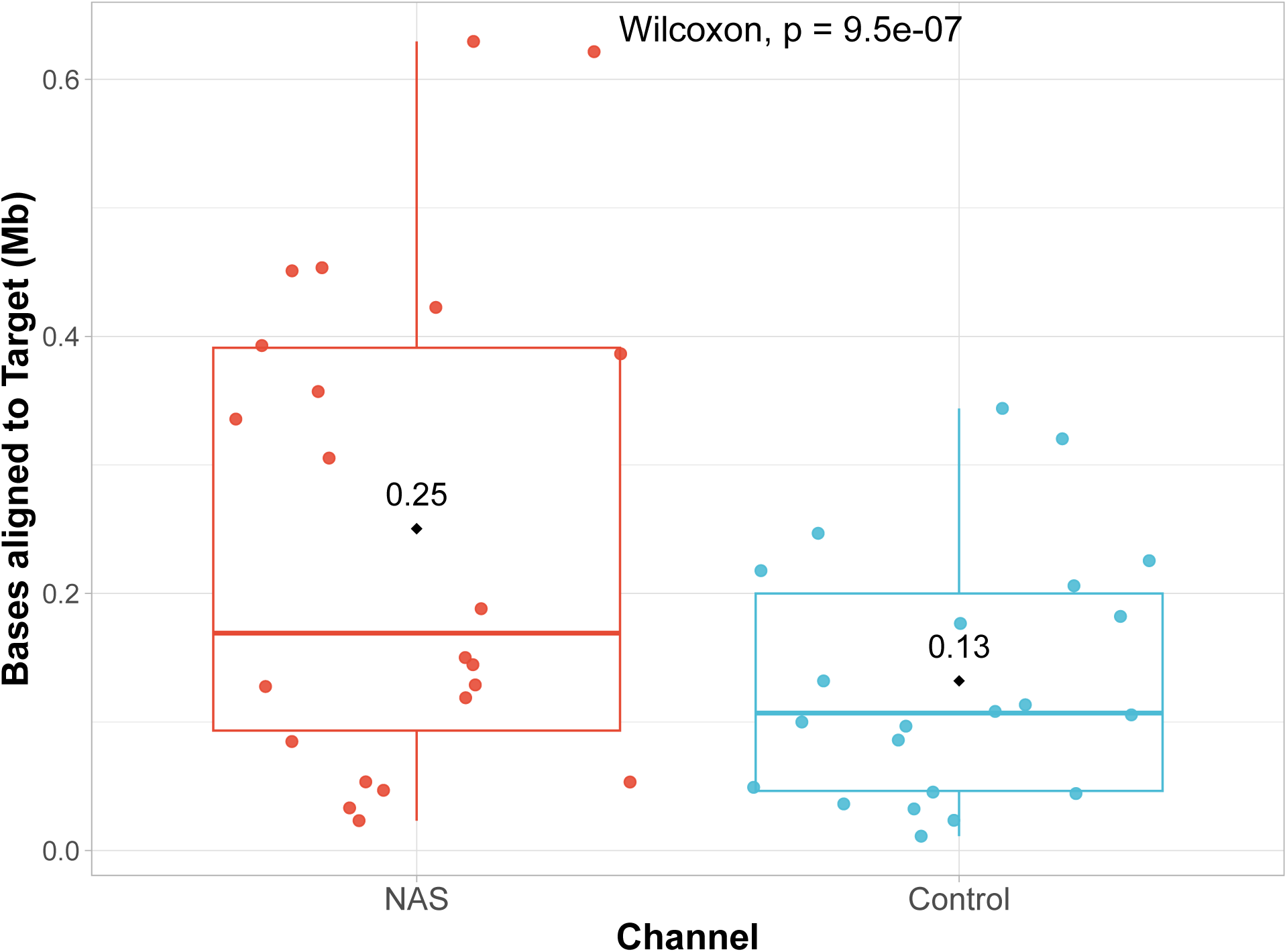
Absolute yield in megabases (Mb) of bases aligning to the individual CBL in multi-serotype samples. Each data point represents the enrichment of a single CBL found within each library. Distributions from control and NAS channels were compared using a paired Wilcoxon test.

**Supplementary Figure 27:**
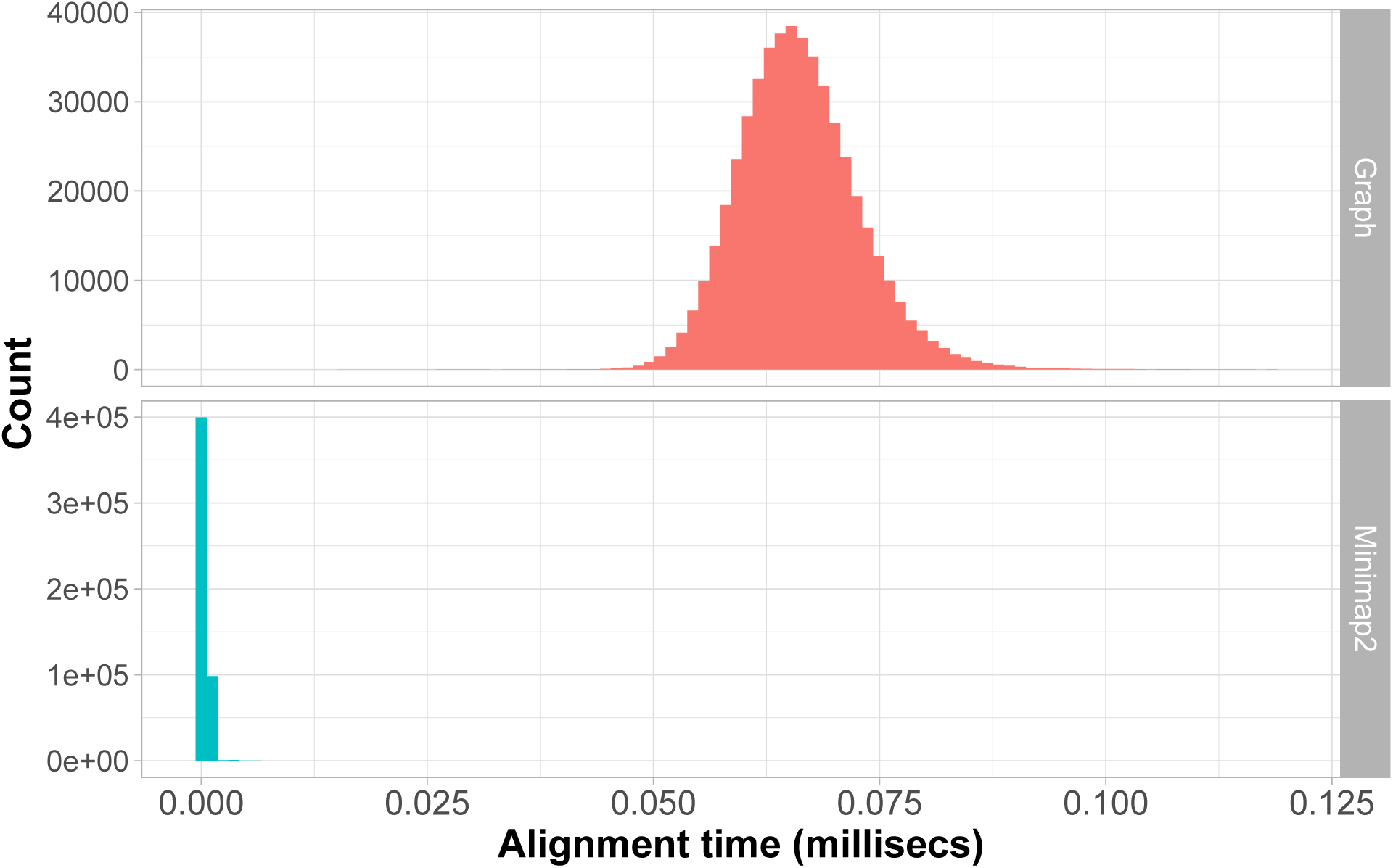
Alignment speed comparison between graph pseudoalignment in GNASTy and Minimap2.

**Supplementary Figure 28:**
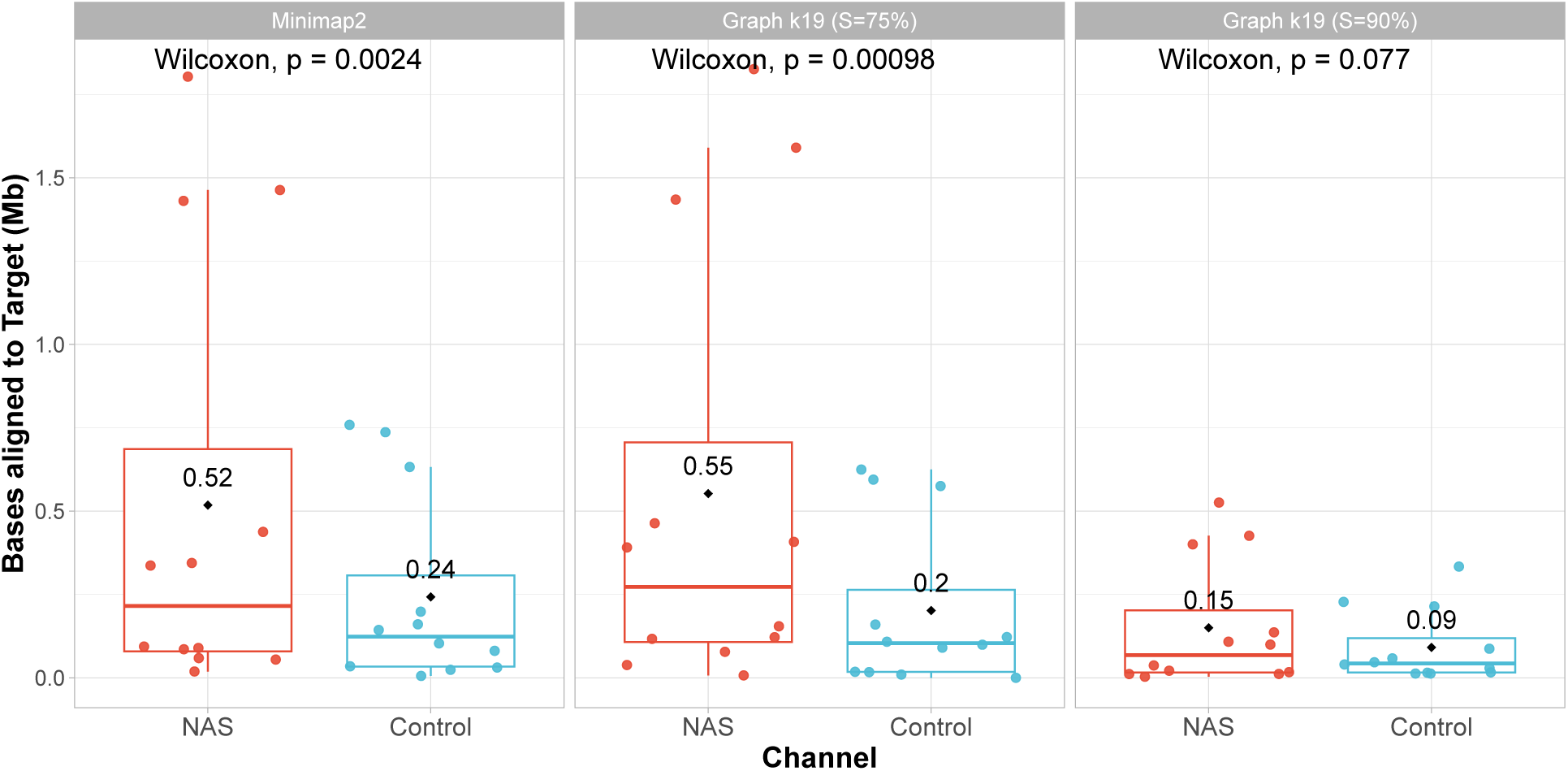
Absolute yield in megabases (Mb) of bases aligning to the 23F CBL when aligning to a partial CBL database using Minimap2 and GNASTy. Each data point represents the enrichment of the 23F CBL found within each library. Distributions from control and NAS channels were compared using a paired Wilcoxon test.

**Supplementary Figure 29:**
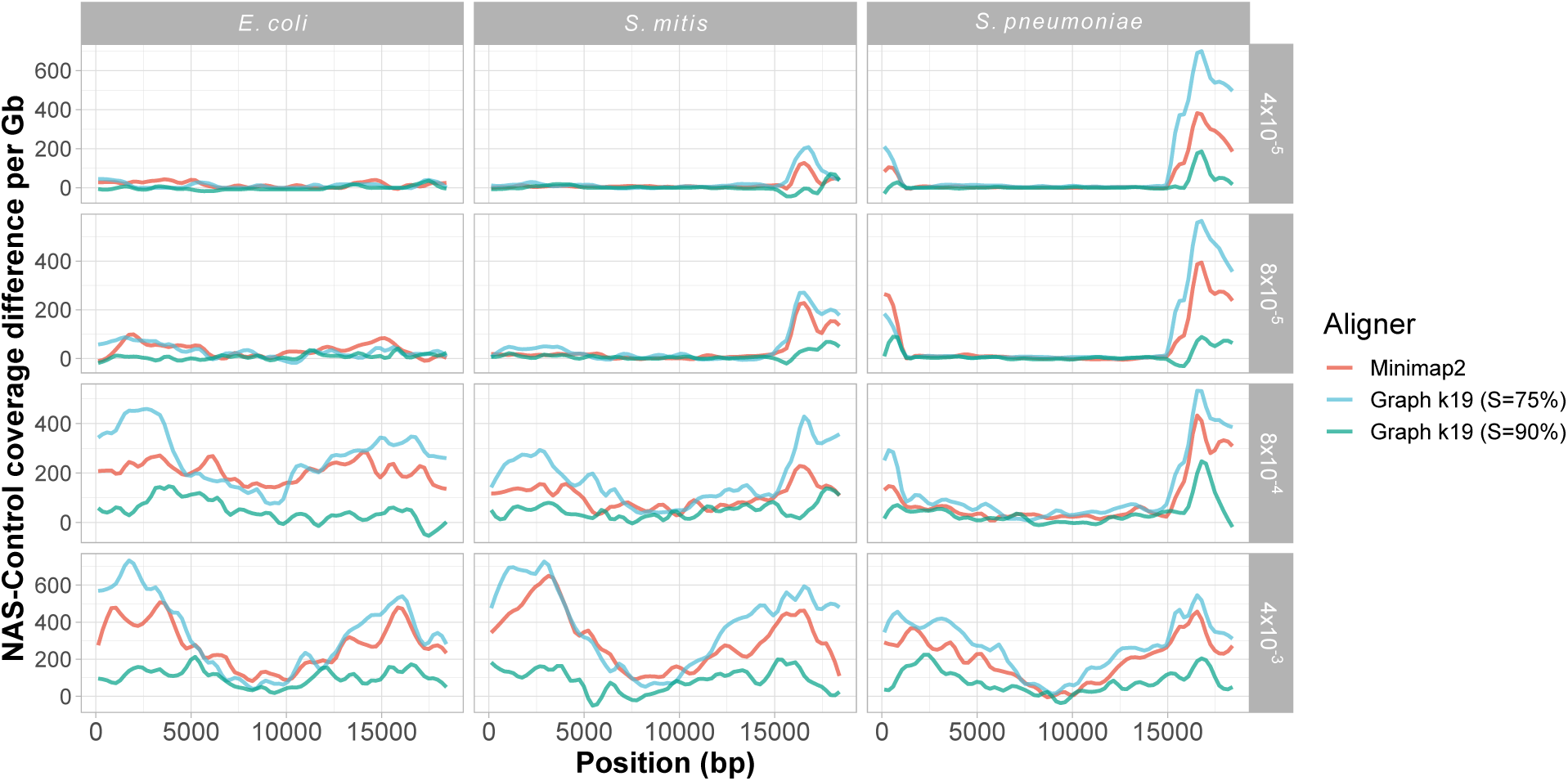
Normalised coverage difference between NAS and control channels across 23F CBL using a partial CBL reference database using Minimap2 and GNASTy. NAS-control coverage difference per gigabase (Gb) calculated by normalising the read coverage for each locus by the amount of data generated (in Gb) for each respective sample and channel, and then negating the normalised coverage for control channels from NAS channels for each locus. Columns describe the non-target species. Rows describe the proportion of target DNA present in the sample.

**Supplementary Figure 30:**
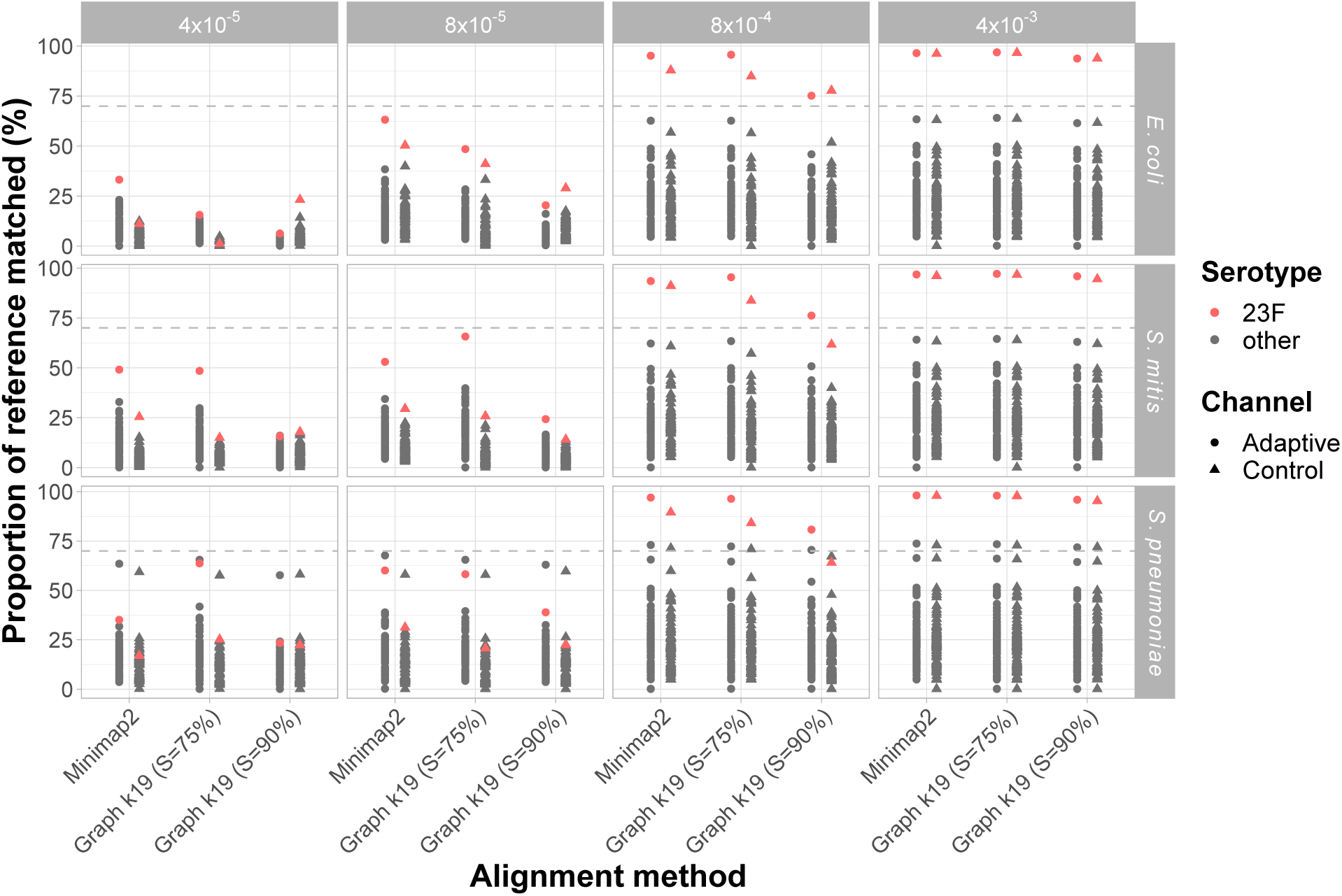
Serotype 23F prediction using reads aligning a to partial CBL reference database using Minimap2 and GNASTy. Y-axis describes the proportion of reference CBL k-mers matched by PneumoKITy (*18*), with each data point describing an individual CBL. The lower limit of matching k-mers used by PneumoKITy (70%) to identify as CBL as present is marked by the grey dotted line. Predictions for the 23F CBL are highlighted in red. Shapes described the channel type (adaptive or control). Columns describe the 23F CBL DNA proportion in the sample. Rows describe the non-target species mixed with Spn23F. Sub-serotypes (e.g. 6A-I, 6A-II etc.) were removed from data to avoid redundancy.

**Supplementary Figure 31:**
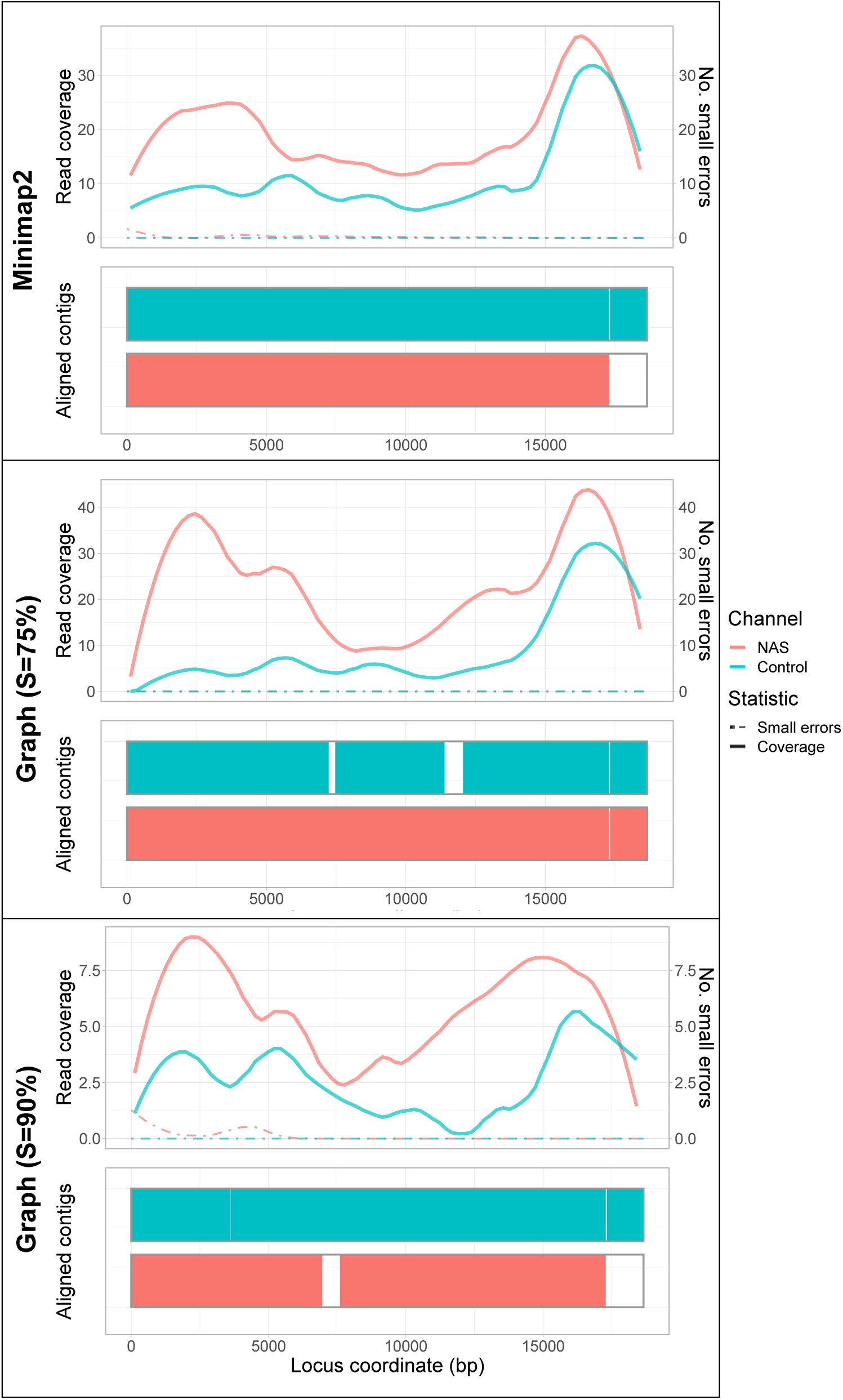
Spn23F CBL assembly comparison across alignment methods using a partial CBL database during NAS using Minimap2 and GNASTy. Each panel describes a 23F CBL assembly generated from 0.1 Spn23F dilutions with *S. mitis*. For each panel, the top plot shows the read coverage (solid), defined as the absolute number of bases aligning to a locus, and number of small errors (*≤* 50 bp, dashed), whilst the bottom plot shows aligned contigs (colours) and large errors (*>* 50 bp) in each assembly.

**Supplementary Figure 32:**
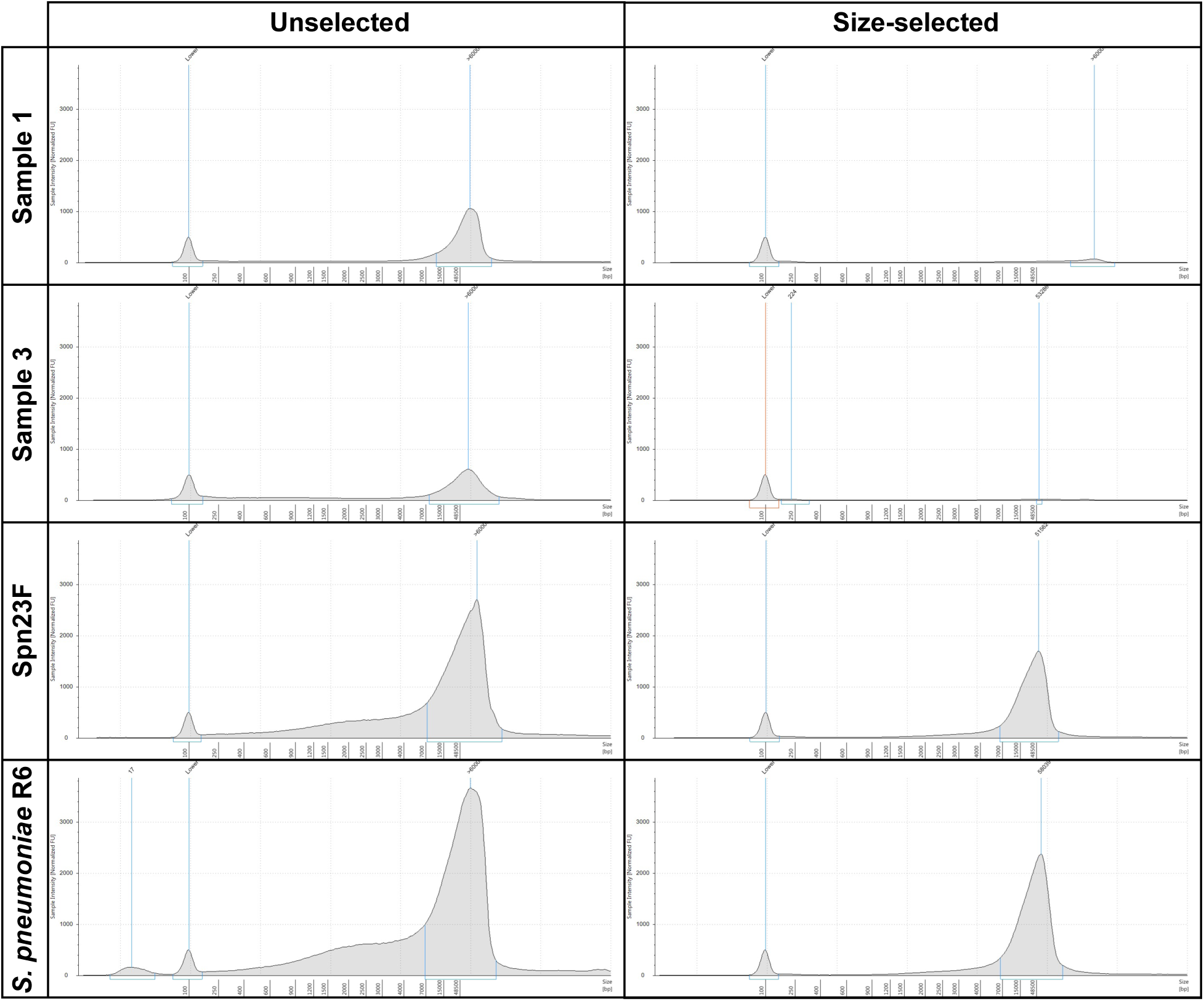
Tapestation images for unselected and size-selected samples. Mixed cultures from nasopharyngeal samples are denoted ‘Sample X’. DNA extractions from single isolate cultures, Spn23F and *S. pneumoniae* R6, were included as positive controls. Y-axis denotes normalised flourescent units (fU). X-axis denotes DNA fragment size (bp). Starting DNA concentrations for size selection: Sample 1: 100 ng/µL, Sample 3: 40 ng/µL, Spn23F: 100 ng/µL, *S. pneumoniae* R6: 100 ng/µL

**Supplementary Figure 33:**
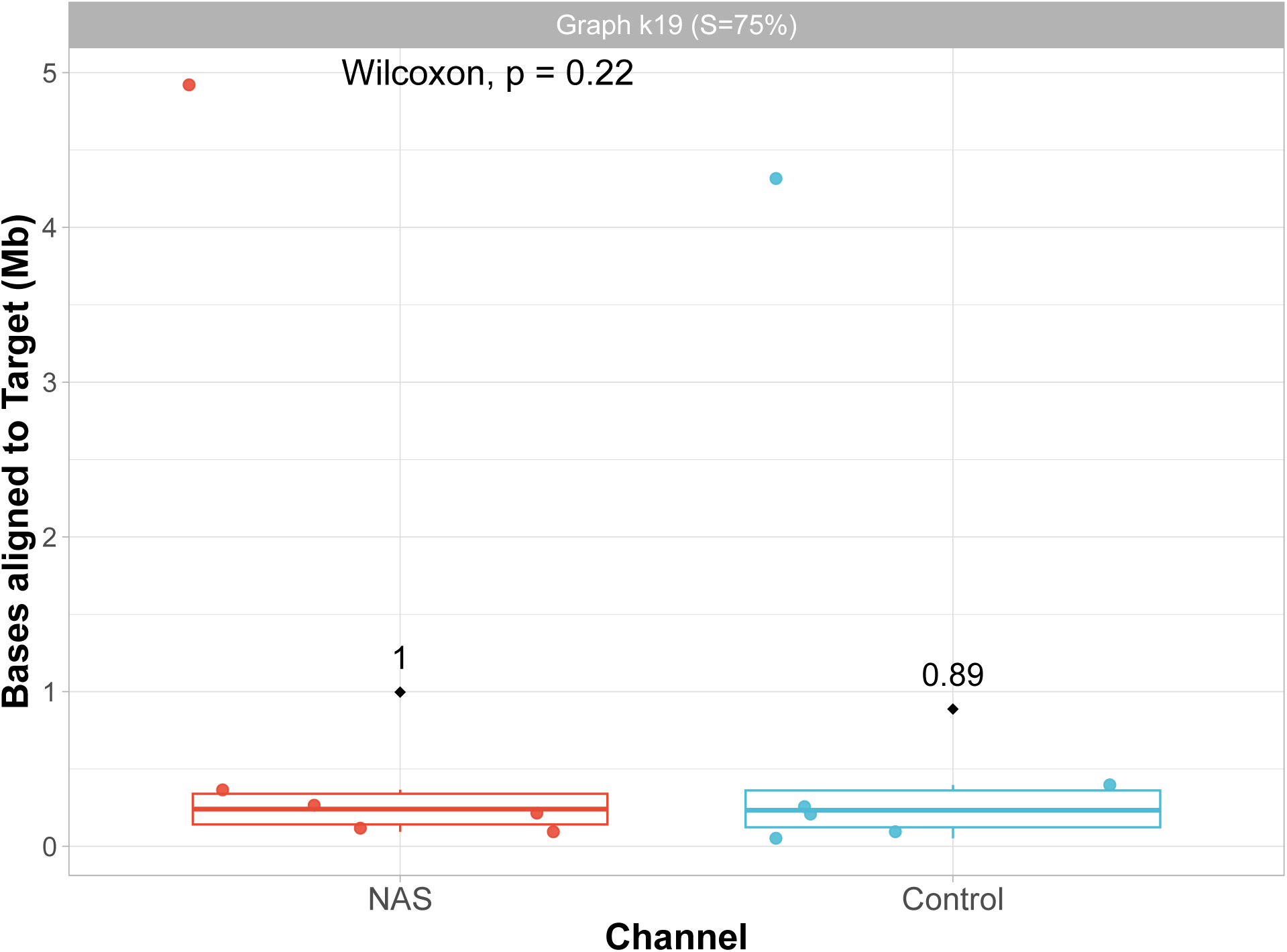
Absolute yield in megabases (Mb) of bases aligning to the 23F CBL when aligning to a full CBL database for mock nasopharyngeal samples using Minimap2 and GNASTy. Each data point represents the enrichment of the 23F CBL found within each library. Distributions from control and NAS channels were compared using a paired Wilcoxon test.

**Supplementary Figure 34:**
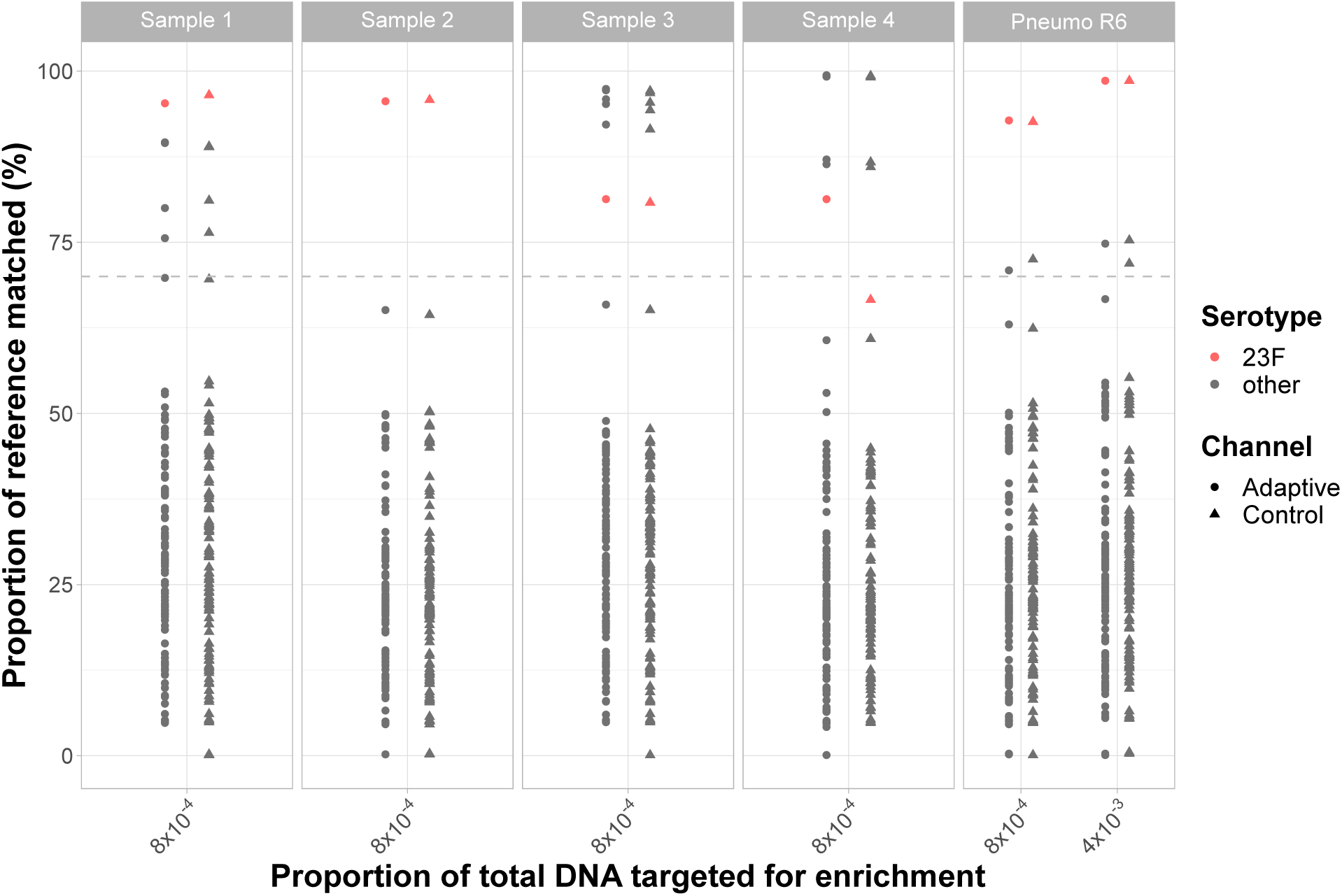
Serotype 23F prediction from mock nasopharyngeal microbiomes using graph pseudoalignment in GNASTy for enrichment. Y-axis describes the proportion of reference CBL k-mers matched by Pneumokity (*18*), with each data point describing an individual CBL. The lower limit of matching k-mers used by PneumoKITy (70%) to identify as CBL as present is marked by the grey dotted line. Predictions for the 23F CBL are highlighted in red. Shapes described the channel type (adaptive or control). X-axis describes the 23F CBL DNA proportion in the sample. Columns describe the nasopharyngeal mixed culture (denoted as ‘Sample X’) or *S. pneumoniae* R6 sample mixed with Spn23F. Subtypes based on *wzy* variation (e.g. 6A-I, 6A-II etc.) were removed from data to avoid redundancy.

**Supplementary Figure 35:**
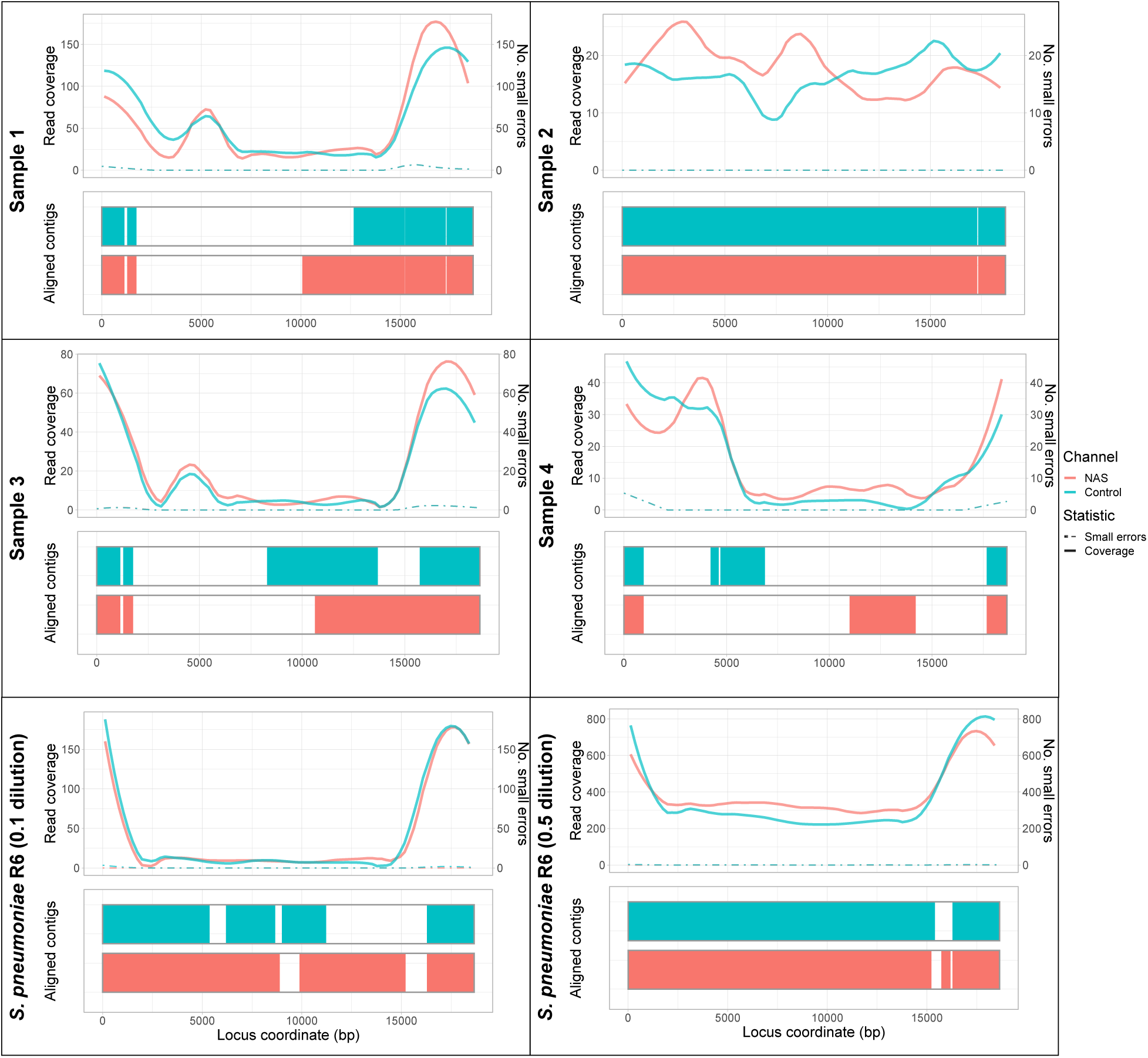
Spn23F CBL assembly from mock nasopharyngeal microbiomes using graph pseudoalignment in GNASTy (*S* = 75%) aligning to a full CBL database during NAS. Each panel describes a 23F CBL assembly generated from 0.1 Spn23F spike into mixed cultures (for ‘Sample X’) or 0.1 and 0.5 Spn23F spikes into *S. pneumoniae* R6. For each panel, the top plot shows the read coverage (solid), defined as the absolute number of bases aligning to a locus, and number of small errors (*≤* 50 bp, dashed), whilst the bottom plot shows the aligned contigs (colours) and large errors (*>* 50 bp) in each assembly.

### C Supplementary Tables

**Supplementary Table 5:**
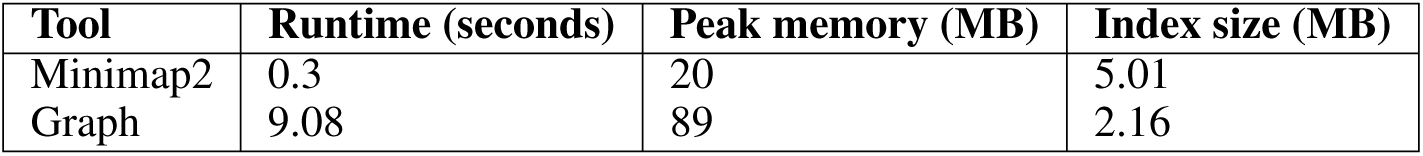
Computational comparison of Minimap2 index and DBG building using *k* = 19 from 106 CBL sequences. Tools were run using 16 threads.

| Tool | Runtime (seconds) | Peak memory (MB) | Index size (MB) |
| --- | --- | --- | --- |
| Minimap2 | 0.3 | 20 | 5.01 |
| Graph | 9.08 | 89 | 2.16 |

**Supplementary Table 6:** Number of reads aligning to 23F CBL when enriching for CBL using NAS.

| Non-target species | Channel | Target proportion | Reads mapped |
| --- | --- | --- | --- |
| *E. coli* | NAS | $4 \times 10^{-5}$ | 1 |
| | | $8 \times 10^{-5}$ | 3 |
| | | $8 \times 10^{-4}$ | 44 |
| | | $4 \times 10^{-3}$ | 203 |
| | Control | $4 \times 10^{-5}$ | 1 |
| | | $8 \times 10^{-5}$ | 2 |
| | | $8 \times 10^{-4}$ | 15 |
| | | $4 \times 10^{-3}$ | 68 |
| *S. mitis* | NAS | $4 \times 10^{-5}$ | 2 |
| | | $8 \times 10^{-5}$ | 6 |
| | | $8 \times 10^{-4}$ | 37 |
| | | $4 \times 10^{-3}$ | 301 |
| | Control | $4 \times 10^{-5}$ | 4 |
| | | $8 \times 10^{-5}$ | 3 |
| | | $8 \times 10^{-4}$ | 22 |
| | | $4 \times 10^{-3}$ | 101 |
| *S. pneumoniae* | NAS | $4 \times 10^{-5}$ | 35 |
| | | $8 \times 10^{-5}$ | 27 |
| | | $8 \times 10^{-4}$ | 42 |
| | | $4 \times 10^{-3}$ | 215 |
| | Control | $4 \times 10^{-5}$ | 10 |
| | | $8 \times 10^{-5}$ | 8 |
| | | $8 \times 10^{-4}$ | 19 |
| | | $4 \times 10^{-3}$ | 78 |

## Notes

https://www.ebi.ac.uk/ena/browser/view/PRJEB72455

https://zenodo.org/records/10581200

https://zenodo.org/records/10581132

https://zenodo.org/records/10590659

### Competing Interest Statement

The authors have declared no competing interest.

## References

1. J. N. Weiser, D. M. Ferreira, J. C. Paton, Streptococcus pneumoniae: transmission, colonization and invasion. Nature Reviews Microbiology, DOI 10.1038/s41579-018-0001-8 (2018).

2. K. S. Ikuta et al., Global mortality associated with 33 bacterial pathogens in 2019: a systematic analysis for the Global Burden of Disease Study 2019. The Lancet 400, 2221–2248, DOI 10.1016/S0140-6736(22)02185-7 (2022).

3. B. Wahl, K. L. O’Brien, A. Greenbaum, A. Majumder, L. Liu, Y. Chu, I. Lukšić, H. Nair, D. A. McAllister, H. Campbell, I. Rudan, R. Black, M. D. Knoll, Burden of Streptococcus pneumoniae and Haemophilus influenzae type b disease in children in the era of conjugate vaccines: global, regional, and national estimates for 2000-15. The Lancet Global Health 6, e744–e757, DOI 10.1016/S2214-109X(18)30247-X (2018).

4. H. Wang et al., Global, regional, and national life expectancy, all-cause mortality, and cause-specific mortality for 249 causes of death, 1980-2015: a systematic analysis for the Global Burden of Disease Study 2015. The Lancet 388, 1459–1544, DOI 10.1016/S0140-6736(16)31012-1 (2016).

5. F. Ganaie, J. S. Saad, L. McGee, A. J. van Tonder, S. D. Bentley, S. W. Lo, R. A. Gladstone, P. Turner, J. D. Keenan, R. F. Breiman, M. H. Nahm, A new pneumococcal capsule type, 10D, is the 100th serotype and has a large cps fragment from an oral streptococcus. mBio 11, DOI 10.1128/mbio.00937-20 (2020).

6. C. Hyams, E. Camberlein, J. M. Cohen, K. Bax, J. S. Brown, The Streptococcus pneumoniae Capsule Inhibits Complement Activity and Neutrophil Phagocytosis by Multiple Mechanisms. Infection and Immunity 78, 704, DOI 10.1128/IAI.00881-09 (2010).

7. N. J. Croucher, A. Løchen, S. D. Bentley, Pneumococcal Vaccines: Host Interactions, Population Dynamics, and Design Principles. Annual Review of Microbiology 72, 521–549, DOI 10.1146/ANNUREV-MICRO-090817-062338 (2018).

8. S. W. Lo, R. A. Gladstone, A. J. van Tonder, J. A. Lees, M. du Plessis, R. Benisty, N. Givon-Lavi, P. A. Hawkins, J. E. Cornick, B. Kwambana-Adams, P. Y. Law, P. L. Ho, M. Antonio, D. B. Everett, R. Dagan, A. von Gottberg, K. P. Klugman, L. McGee, R. F. Breiman, S. D. Bentley, A. W. Brooks, A. Corso, A. Davydov, A. Maguire, A. Pollard, A. Kiran, A. Skoczynska, B. Moiane, B. Beall, B. Sigauque, D. Aanensen, D. Lehmann, D. Faccone, E. Foster-Nyarko, E. Bojang, E. Egorova, E. Voropaeva, E. Sampane-Donkor, E. Sadowy, G. Bigogo, H. Mucavele, H. Belabbès, I. Diawara, J. Moïsi, J. Verani, J. Keenan, J. N. Nair Thulasee Bhai, K. M. Ndlangisa, K. Zerouali, K. L. Ravikumar, L. Titov, L. De Gouveia, M. Alaerts, M. Ip, M. C. de Cunto Brandileone, M. Hasanuzzaman, M. Paragi, M. Nurse-Lucas, M. Ali, N. Elmdaghri, N. Croucher, N. Wolter, N. Porat, Ö. Köseoglu Eser, P. E. Akpaka, P. Turner, P. Gagetti, P. E. Tientcheu, P. E. Carter, R. Mostowy, R. Kandasamy, R. Ford, R. Henderson, R. Malaker, S. Shakoor, S. C. Grassi Almeida, S. K. Saha, S. Doiphode, S. A. Madhi, S. Devi Sekaran, S. Srifuengfung, S. Obaro, S. C. Clarke, S. A. Nzenze, T. Kastrin, T. J. Ochoa, V. Balaji, W. Hryniewicz, Y. Urban, Pneumococcal lineages associated with serotype replacement and antibiotic resistance in childhood invasive pneumococcal disease in the post-PCV13 era: an international whole-genome sequencing study. The Lancet Infectious Diseases 19, 759–769, DOI 10.1016/S1473-3099(19)30297-X (2019).

9. A. J. van Tonder, R. A. Gladstone, S. W. Lo, M. H. Nahm, M. du Plessis, J. Cornick, B. Kwambana-Adams, S. A. Madhi, P. A. Hawkins, R. Benisty, R. Dagan, D. Everett, M. Antonio, K. P. Klugman, A. von Gottberg, R. F. Breiman, L. McGee, S. D. Bentley, Putative novel cps loci in a large global collection of pneumococci. Microbial Genomics 5, e000274, DOI 10.1099/mgen.0.000274 (2019).

10. S. N. Ladhani, S. Collins, A. Djennad, C. L. Sheppard, R. Borrow, N. K. Fry, N. J. Andrews, E. Miller, M. E. Ramsay, Rapid increase in non-vaccine serotypes causing invasive pneumococcal disease in England and Wales, 2000-17: a prospective national observational cohort study. The Lancet Infectious Diseases 18, 441–451, DOI 10.1016/S1473-3099(18)30052-5 (2018).

11. S. W. Lo et al., Emergence of a multidrug-resistant and virulent Streptococcus pneumoniae lineage mediates serotype replacement after PCV13: an international whole-genome sequencing study. The Lancet Microbe 3, e735–e743, DOI 10.1016/S2666-5247(22)00158-6 (2022).

12. R. E. Huebner, R. Dagan, N. Porath, A. D. Wasas, M. T, K. P. Klugman, Lack of utility of serotyping multiple colonies for detection of simultaneous nasopharyngeal carriage of different pneumococcal serotypes. The Pediatric Infectious Disease Journal 19, 1017–1020, DOI 10.1097/00006454-200010000-00019 (2000).

13. G. Tonkin-Hill, C. Ling, C. Chaguza, S. J. Salter, P. Hinfonthong, E. Nikolaou, N. Tate, A. Pastusiak, C. Turner, C. Chewapreecha, S. D. Frost, J. Corander, N. J. Croucher, P. Turner, S. D. Bentley, Pneumococcal within-host diversity during colonization, transmission and treatment. Nature Microbiology 7, 1791–1804, DOI 10.1038/s41564-022-01238-1 (2022).

14. C. Colijn, J. Corander, N. J. Croucher, Designing ecologically optimized pneumococcal vaccines using population genomics. Nature Microbiology 5, 473–485, DOI 10.1038/s41564-019-0651-y (2020).

15. P. Turner, J. Hinds, C. Turner, A. Jankhot, K. Gould, S. D. Bentley, F. Nosten, D. Goldblatt, Improved detection of nasopharyngeal cocolonization by multiple pneumococcal serotypes by use of latex agglutination or molecular serotyping by microarray. Journal of Clinical Microbiology 49, 1784–1789, DOI 10.1128/JCM.00157-11 (2011).

16. M. Habib, B. D. Porter, C. Satzke, Capsular serotyping of Streptococcus pneumoniae using the Quellung reaction. Journal of Visualized Experiments, DOI 10.3791/51208 (2014).

17. C. Satzke, E. M. Dunne, B. D. Porter, K. P. Klugman, E. K. Mulholland, J. E. Vidal, F. Sakai, J. E. Strachan, D. C. Hay Burgess, D. Holtzman, K. Boelsen, M. Habib, J. Manning, B. D. Ortika, C. L. Pell, J. A. Smyth, M. Antonio, K. P. Klugman, K. L. O’Brien, R. M. Robins-Browne, J. Anthony Scott, S. K. Saha, F. M. Russell, A. R. Greenhill, D. Lehmann, P. V. Adrian, S. A. Madhi, L. G. Rubin, A. Rizvi, J. Hinds, K. A. Gould, F. Kong, S. Oftadeh, G. L. Gilbert, L. Feng, B. Cao, G. Paranhos-Baccalà, J. N. Telles, M. Messaoudi, R. Borrow, E. Stanford, R. George, H. C. Slotved, S. D. Brugger, K. Mühlemann, M. Hilty, I. A. Rivera-Olivero, J. H. de Waard, B. M. Charalambous, M. H. Leung, C. Azzari, M. Moriondo, F. Nieddu, P. W. Hermans, C. E. van der Gaast-de Jongh, P. Turner, D. J. Ecker, R. Sampath, The PneuCarriage Project: A Multi-Centre Comparative Study to Identify the Best Serotyping Methods for Examining Pneumococcal Carriage in Vaccine Evaluation Studies. PLoS Medicine 12, DOI 10.1371/JOURNAL.PMED.1001903 (2015).

18. C. L. Sheppard, S. Manna, N. Groves, D. J. Litt, Z. Amin-Chowdhury, M. Bertran, S. Ladhani, C. Satzke, N. K. Fry, PneumoKITy: A fast, flexible, specific, and sensitive tool for Streptococcus pneumoniae serotype screening and mixed serotype detection from genome sequence data. Microbial Genomics 8, DOI 10.1099/MGEN.0.000904 (2022).

19. L. Epping, A. J. van Tonder, R. A. Gladstone, S. D. Bentley, A. J. Page, J. A. Keane, SeroBA: rapid high-throughput serotyping of Streptococcus pneumoniae from whole genome sequence data. Microbial Genomics 4, DOI 10.1099/MGEN.0.000186 (2018).

20. G. Kapatai, C. L. Sheppard, A. Al-Shahib, D. J. Litt, A. P. Underwood, T. G. Harrison, N. K. Fry, Whole genome sequencing of Streptococcus pneumoniae: Development, evaluation and verification of targets for serogroup and serotype prediction using an automated pipeline. PeerJ, DOI 10.7717/peerj.2477 (2016).

21. S. D. Bentley, D. M. Aanensen, A. Mavroidi, D. Saunders, E. Rabbinowitsch, M. Collins, K. Donohoe, D. Harris, L. Murphy, M. A. Quail, G. Samuel, I. C. Skovsted, M. S. Kaltoft, B. Barrell, P. R. Reeves, J. Parkhill, B. G. Spratt, Genetic analysis of the capsular biosynthetic locus from all 90 pneumococcal serotypes. PLoS Genetics 2, 0262–0269, DOI 10.1371/journal.pgen.0020031 (2006).

22. E. Jauneikaite, A. S. Tocheva, J. M. Jefferies, R. A. Gladstone, S. N. Faust, M. Christodoulides, M. L. Hibberd, S. C. Clarke, Current methods for capsular typing of Streptococcus pneumoniae. Journal of Microbiological Methods 113, 41–49, DOI 10.1016/J.MIMET.2015.03.006 (2015).

23. L. J. Ricketson, R. Lidder, R. Thorington, I. Martin, O. G. Vanderkooi, M. Sadarangani, J. D. Kellner, PCR and culture analysis of streptococcus pneumoniae nasopharyngeal carriage in healthy children. Microorganisms 9, DOI 10.3390/microorganisms9102116 (2021).

24. C. L. Ip, M. Loose, J. R. Tyson, M. de Cesare, B. L. Brown, M. Jain, R. M. Leggett, D. A. Eccles, V. Zalunin, J. M. Urban, P. Piazza, R. J. Bowden, B. Paten, S. Mwaigwisya, E. M. Batty, J. T. Simpson, T. P. Snutch, E. Birney, D. Buck, S. Goodwin, H. J. Jansen, J. O’Grady, H. E. Olsen, MinION Analysis and Reference Consortium: Phase 1 data release and analysis. F1000Research 4, 1075, DOI 10.12688/f1000research.7201.1 (2015).

25. J. Quick et al., Real-time, portable genome sequencing for Ebola surveillance. eng, Nature 530, 228–232, DOI 10.1038/nature16996 (2016).

26. A. Payne, N. Holmes, T. Clarke, R. Munro, B. J. Debebe, M. Loose, Readfish enables targeted nanopore sequencing of gigabase-sized genomes. Nature Biotechnology 39, 442, DOI 10.1038/S41587-020-00746-X (2021).

27. L. Weilguny, N. De Maio, R. Munro, C. Manser, E. Birney, M. Loose, N. Goldman, Dynamic, adaptive sampling during nanopore sequencing using Bayesian experimental design. Nature Biotechnology 41, 1018–1025, DOI 10.1038/S41587-022-01580-Z (2023).

28. S. H. Ye, K. J. Siddle, D. J. Park, P. C. Sabeti, Benchmarking Metagenomics Tools for Taxonomic Classification. Cell 178, 779–794, DOI 10.1016/j.cell.2019.07.010 (2019).

29. M. Marquet, J. Zöllkau, J. Pastuschek, A. Viehweger, E. Schleußner, O. Makarewicz, M. W. Pletz, R. Ehricht, C. Brandt, Evaluation of microbiome enrichment and host DNA depletion in human vaginal samples using Oxford Nanopore’s adaptive sequencing. Scientific Reports 12, DOI 10.1038/S41598-022-08003-8 (2022).

30. J. Su, W. W. Lui, Y. Lee, Z. Zheng, G. K.-H. Siu, T. T.-L. Ng, T. Zhang, T. T.-Y. Lam, H.-Y. Lao, W.-C. Yam, K. K.-G. Tam, K. S.-S. Leung, T.-W. Lam, A. W.-S. Leung, R. Luo, Evaluation of Mycobacterium tuberculosis enrichment in metagenomic samples using ONT adaptive sequencing and amplicon sequencing for identification and variant calling. Scientific Reports 13, 5237, DOI 10.1038/S41598-023-32378-X (2023).

31. S. Martin, D. Heavens, Y. Lan, S. Horsfield, M. D. Clark, R. M. Leggett, Nanopore adaptive sampling: a tool for enrichment of low abundance species in metagenomic samples. Genome Biology 23, 1–27, DOI 10.1186/S13059-021-02582-X (2022).

32. D. C. Wrenn, D. M. Drown, Nanopore adaptive sampling enriches for antimicrobial resistance genes in microbial communities. Gigabyte, 1–14, DOI 10.46471/GIGABYTE.103 (2023).

33. A. Viehweger, M. Marquet, M. Hölzer, N. Dietze, M. W. Pletz, C. Brandt, Nanopore based enrichment of antimicrobial resistance genes - a case-based study. GigaByte, 1–15, DOI 10.46471/GIGABYTE.75 (2023).

34. P. Marttinen, N. J. Croucher, M. U. Gutmann, J. Corander, W. P. Hanage, Recombination produces coherent bacterial species clusters in both core and accessory genomes. Microbial Genomics 1, DOI 10.1099/MGEN.0.000038 (2015).

35. J. C. D’Aeth, M. P. van der Linden, L. McGee, H. de Lencastre, P. Turner, J. H. Song, S. W. Lo, R. A. Gladstone, R. Sá-Leão, K. S. Ko, W. P. Hanage, R. F. Breiman, B. Beall, S. D. Bentley, N. J. Croucher, The role of inter-species recombination in the evolution of antibiotic-resistant pneumococci. eLife 10, DOI 10.7554/ELIFE.67113 (2021).

36. M. Bek-Thomsen, H. Tettelin, I. Hance, K. E. Nelson, M. Kilian, Population diversity and dynamics of Streptococcus mitis, Streptococcus oralis, and Streptococcus infantis in the upper respiratory tracts of adults, determined by a nonculture strategy. Infection and Immunity 76, 1889–1896, DOI 10.1128/IAI.01511-07 (2008).

37. C. Delahaye, J. Nicolas, Sequencing DNA with nanopores: Troubles and biases. PLoS ONE 16, DOI 10.1371/JOURNAL.PONE.0257521 (2021).

38. A. Løchen, J. E. Truscott, N. J. Croucher, Analysing pneumococcal invasiveness using Bayesian models of pathogen progression rates. PLoS Computational Biology 18, e1009389, DOI 10.1371/JOURNAL.PCBI.1009389 (2022).

39. N. J. Croucher, D. Walker, P. Romero, N. Lennard, G. K. Paterson, N. C. Bason, A. M. Mitchell, M. A. Quail, P. W. Andrew, J. Parkhill, S. P. Bentley, T. J. Mitchell, Role of Conjugative Elements in the Evolution of the Multidrug-Resistant Pandemic Clone Streptococcus pneumoniae Spain23F ST81. Journal of Bacteriology 191, 1480, DOI 10.1128/JB.01343-08 (2009).

40. M. Kolmogorov, D. M. Bickhart, B. Behsaz, A. Gurevich, M. Rayko, S. B. Shin, K. Kuhn, J. Yuan, E. Polevikov, T. P. Smith, P. A. Pevzner, metaFlye: scalable long-read metagenome assembly using repeat graphs. Nature Methods 17, 1103–1110, DOI 10.1038/s41592-020-00971-x (2020).

41. Y. Chen, Y. Zhang, A. Y. Wang, M. Gao, Z. Chong, Accurate long-read de novo assembly evaluation with Inspector. Genome Biology 22, 1–21, DOI 10.1186/S13059-021-02527-4 (2021).

42. N. J. Croucher, S. R. Harris, L. Barquist, J. Parkhill, S. D. Bentley, A high-resolution view of genome-wide pneumococcal transformation. PLoS Pathogens 8, DOI 10.1371/JOURNAL.PPAT.1002745 (2012).

43. M. Cretu Stancu, M. J. Van Roosmalen, I. Renkens, M. M. Nieboer, S. Middelkamp, J. De Ligt, G. Pregno, D. Giachino, G. Mandrile, J. Espejo Valle-Inclan, J. Korzelius, E. De Bruijn, E. Cuppen, M. E. Talkowski, T. Marschall, J. De Ridder, W. P. Kloosterman, Mapping and phasing of structural variation in patient genomes using nanopore sequencing. Nature Communications 8, 1–13, DOI 10.1038/s41467-017-01343-4 (2017).

44. E. Garrison, J. Sirén, A. M. Novak, G. Hickey, J. M. Eizenga, E. T. Dawson, W. Jones, S. Garg, C. Markello, M. F. Lin, B. Paten, R. Durbin, Variation graph toolkit improves read mapping by representing genetic variation in the reference. Nature Biotechnology 36, 875–881, DOI 10.1038/nbt.4227 (2018).

45. A. Dilthey, C. Cox, Z. Iqbal, M. R. Nelson, G. McVean, Improved genome inference in the MHC using a population reference graph. Nature Genetics 47, 682–688, DOI 10.1038/ng.3257 (2015).

46. S. T. Horsfield, G. Tonkin-Hill, N. J. Croucher, J. A. Lees, Accurate and fast graph-based pangenome annotation and clustering with ggCaller. Genome Research 33, gr.277733.123, DOI 10.1101/GR.277733.123 (2023).

47. Z. Iqbal, M. Caccamo, I. Turner, P. Flicek, G. McVean, De novo assembly and genotyping of variants using colored de Bruijn graphs. Nature Genetics 44, 226–232, DOI 10.1038/ng.1028 (2012).

48. N. L. Bray, H. Pimentel, P. Melsted, L. Pachter, Near-optimal probabilistic RNA-seq quantification. Nature biotechnology 34, 525–527, DOI 10.1038/NBT.3519 (2016).

49. T. Mäklin, T. Kallonen, J. Alanko, Ø. Samuelsen, K. Hegstad, V. Mäkinen, J. Corander, E. Heinz, A. Honkela, Bacterial genomic epidemiology with mixed samples. Microbial Genomics 7, 691, DOI 10.1099/MGEN.0.000691 (2021).

50. J. N. Alanko, J. Vuohtoniemi, T. Mä Klin, S. J. Puglisi, Themisto: a scalable colored k-mer index for sensitive pseudoalignment against hundreds of thousands of bacterial genomes. Bioinformatics 39, i260–i269, DOI 10.1093/BIOINFORMATICS/BTAD233 (2023).

51. G. Holley, P. Melsted, Bifrost: highly parallel construction and indexing of colored and compacted de Bruijn graphs. Genome Biology 21, 249, DOI 10.1186/s13059-020-02135-8 (2020).

52. H. Li, Minimap2: pairwise alignment for nucleotide sequences. Bioinformatics 34, 3094–3100, DOI 10.1093/bioinformatics/bty191 (2018).

53. A. Mavroidi, D. M. Aanensen, D. Godoy, I. C. Skovsted, M. S. Kaltoft, P. R. Reeves, S. D. Bentley, B. G. Spratt, Genetic Relatedness of the Streptococcus pneumoniae Capsular Biosynthetic Loci. Journal of Bacteriology 189, 7841, DOI 10.1128/JB.00836-07 (2007).

54. R. Sinclair Dokos, Update from Oxford Nanopore Technologies. London Calling 2022, (https://nanoporetech.com/london-calling-2022-collection) (2022).

55. U. B. Skov Sørensen, K. Yao, Y. Yang, H. Tettelin, M. Kilian, Capsular Polysaccharide Expression in Commensal Streptococcus Species: Genetic and Antigenic Similarities to Streptococcus pneumoniae. mBio 7, DOI 10.1128/MBIO.01844-16 (2016).

56. J. Hoskins, J. Alborn, J. Arnold, L. C. Blaszczak, S. Burgett, B. S. Dehoff, S. T. Estrem, L. Fritz, D. J. Fu, W. Fuller, C. Geringer, R. Gilmour, J. S. Glass, H. Khoja, A. R. Kraft, R. E. Lagace, D. J. LeBlanc, L. N. Lee, E. J. Lefkowitz, J. Lu, P. Matsushima, S. M. McAhren, M. McHenney, K. McLeaster, C. W. Mundy, T. I. Nicas, F. H. Norris, M. O’Gara, R. B. Peery, G. T. Robertson, P. Rockey, P. M. Sun, M. E. Winkler, Y. Yang, M. Young-Bellido, G. Zhao, C. A. Zook, R. H. Baltz, S. R. Jaskunas, J. Rosteck, P. L. Skatrud, J. I. Glass, Genome of the Bacterium Streptococcus pneumoniae Strain R6. Journal of Bacteriology 183, 5709, DOI 10.1128/JB.183.19.5709-5717.2001 (2001).

57. F. Iannelli, B. J. Pearce, G. Pozzi, The Type 2 Capsule Locus of Streptococcus pneumoniae. Journal of Bacteriology 181, 2652, DOI 10.1128/JB.181.8.2652-2654.1999 (1999).

58. S. J. Salter, C. Turner, W. Watthanaworawit, M. C. de Goffau, J. Wagner, J. Parkhill, S. D. Bentley, D. Goldblatt, F. Nosten, P. Turner, A longitudinal study of the infant nasopharyngeal microbiota: The effects of age, illness and antibiotic use in a cohort of South East Asian children. PLoS Neglected Tropical Diseases 11, e0005975, DOI 10.1371/JOURNAL.PNTD.0005975 (2017).

59. C. Troeger et al., Estimates of the global, regional, and national morbidity, mortality, and aetiologies of lower respiratory infections in 195 countries, 1990–2016: a systematic analysis for the Global Burden of Disease Study 2016. The Lancet Infectious Diseases 18, 1191, DOI 10.1016/S1473-3099(18)30310-4 (2018).

60. M. T. Nelson, C. E. Pope, R. L. Marsh, D. J. Wolter, E. J. Weiss, K. R. Hager, A. T. Vo, M. J. Brittnacher, M. C. Radey, H. S. Hayden, A. Eng, S. I. Miller, E. Borenstein, L. R. Hoffman, Human and Extracellular DNA Depletion for Metagenomic Analysis of Complex Clinical Infection Samples Yields Optimized Viable Microbiome Profiles. Cell Reports 26, 2227, DOI 10.1016/J.CELREP.2019.01.091 (2019).

61. T. Charalampous, G. L. Kay, H. Richardson, A. Aydin, R. Baldan, C. Jeanes, D. Rae, S. Grundy, D. J. Turner, J. Wain, R. M. Leggett, D. M. Livermore, J. O’Grady, Nanopore metagenomics enables rapid clinical diagnosis of bacterial lower respiratory infection. Nature Biotechnology 37, 783–792, DOI 10.1038/S41587-019-0156-5 (2019).

62. P. Turner, C. Turner, A. Jankhot, N. Helen, S. J. Lee, N. P. Day, N. J. White, F. Nosten, D. Goldblatt, A Longitudinal Study of Streptococcus pneumoniae Carriage in a Cohort of Infants and Their Mothers on the Thailand-Myanmar Border. PLoS ONE 7, DOI 10.1371/JOURNAL.PONE.0038271 (2012).

63. P. Turner, C. Turner, N. Green, L. Ashton, E. Lwe, A. Jankhot, N. P. Day, N. J. White, F. Nosten, D. Goldblatt, Serum antibody responses to pneumococcal colonization in the first 2 years of life: results from an SE Asian longitudinal cohort study. Clinical Microbiology and Infection 19, E551, DOI 10.1111/1469-0691.12286 (2013).

64. S. T. Horsfield, Nanopore Adaptive Sampling Analysis Scripts, 2024, DOI 10.5281/zenodo.10581200.

65. S. T. Horsfield, Graph-based Nanopore Adaptive Sampling Typing (GNASTy), 2024, DOI 10.5281/zenodo.10581132.

66. S. T. Horsfield, B. Fok, Y. Fu, P. Turner, J. A. Lees, N. J. Croucher, Nanopore Adaptive Sampling for pneumococcal surveillance using serotyping, 2024, (https://www.ebi.ac.uk/ena/browser/view/PRJEB72455).

67. S. T. Horsfield, PnuemoKITy-Nanopore_v1.0.1, 2024, DOI 10.5281/zenodo.10590659.

68. C. Yang, J. Chu, R. L. Warren, I. Birol, NanoSim: nanopore sequence read simulator based on statistical characterization. GigaScience 6, 1–6, DOI 10.1093/gigascience/gix010 (2017).

69. K. B̌rinda, C. Yang, J. Chu, J. Linthorst, W. Franus, NanoSim-H; a simulator of Oxford Nanopore reads; a fork of NanoSim. 2018, DOI 10.5281/zenodo.1341250.

70. E. Kovács, J. Sahin-Tóth, A. Tóthpál, M. van der Linden, T. Tirczka, O. Dobay, Co-carriage of Staphylococcus aureus, Streptococcus pneumoniae, Haemophilus influenzae and Moraxella catarrhalis among three different age categories of children in Hungary. PLoS ONE 15, DOI 10.1371/journal.pone.0229021 (2020).

71. E. M. Dunne, C. Murad, S. Sudigdoadi, E. Fadlyana, R. Tarigan, S. A. K. Indriyani, C. L. Pell, E. Watts, C. Satzke, J. Hinds, N. E. Dewi, F. F. Yani, K. Rusmil, E. Kim Mulholland, C. Kartasasmita, Carriage of streptococcus pneumoniae, haemophilus influenzae, moraxella catarrhalis, and staphylococcus aureus in Indonesian children: A cross-sectional study. PLoS ONE 13, DOI 10.1371/journal.pone.0195098 (2018).

72. S. Chochua, V. D’Acremont, C. Hanke, D. Alfa, J. Shak, M. Kilowoko, E. Kyungu, L. Kaiser, B. Genton, K. P. Klugman, J. E. Vidal, Increased Nasopharyngeal Density and Concurrent Carriage of Streptococcus pneumoniae, Haemophilus influenzae, and Moraxella catarrhalis Are Associated with Pneumonia in Febrile Children. PLOS ONE 11, e0167725, DOI 10.1371/JOURNAL.PONE.0167725 (2016).

73. J. M. Eizenga, A. M. Novak, J. A. Sibbesen, S. Heumos, A. Ghaffaari, G. Hickey, X. Chang, J. D. Seaman, R. Rounthwaite, J. Ebler, M. Rautiainen, S. Garg, B. Paten, T. Marschall, J. Sirén, E. Garrison, Pangenome Graphs. Annual Review of Genomics and Human Genetics 21, DOI 10.1146/annurev-genom-120219-080406 (2020).

74. G. Hickey, D. Heller, J. Monlong, J. A. Sibbesen, J. Sirén, J. Eizenga, E. T. Dawson, E. Garrison, A. M. Novak, B. Paten, Genotyping structural variants in pangenome graphs using the vg toolkit. Genome Biology 21, 35, DOI 10.1186/s13059-020-1941-7 (2020).

75. J. A. Sibbesen, J. M. Eizenga, A. M. Novak, J. Sirén, X. Chang, E. Garrison, B. Paten, Haplotype-aware pantranscriptome analyses using spliced pangenome graphs. Nature Methods 20, 239–247, DOI 10.1038/s41592-022-01731-9 (2023).

76. B. D. Ondov, T. J. Treangen, P. Melsted, A. B. Mallonee, N. H. Bergman, S. Koren, A. M. Phillippy, Mash: Fast genome and metagenome distance estimation using MinHash. Genome Biology 17, DOI 10.1186/s13059-016-0997-x (2016).

77. S. R. Harris, SKA: Split Kmer Analysis Toolkit for Bacterial Genomic Epidemiology. bioRxiv, 453142, DOI 10.1101/453142 (2018).

78. H. Fan, A. R. Ives, Y. Surget-Groba, C. H. Cannon, An assembly and alignment-free method of phylogeny reconstruction from next-generation sequencing data. BMC Genomics 16, DOI 10.1186/S12864-015-1647-5 (2015).

79. R. Munro, A. Payne, M. Loose, Icarust, a real-time simulator for Oxford Nanopore adaptive sampling. bioRxiv, 2023.05.16.540986, DOI 10.1101/2023.05.16.540986 (2023).

80. A. E. Darling, B. Mau, N. T. Perna, progressiveMauve: Multiple Genome Alignment with Gene Gain, Loss and Rearrangement. PLoS ONE 5, e11147, DOI 10.1371/JOURNAL.PONE.0011147 (2010).

